# interpreTFM: an interpretable framework for decoding transcriptomic foundation models

**DOI:** 10.64898/2026.11.16.767062

**Authors:** Yunfei Hu, Zihang Wen, Yichen Henry Liu, Weiman Yuan, Haoran Qin, Zhenhan Lin, David Klindt, Haichun Yang, Oana Carja, Xin Maizie Zhou

## Abstract

**Background:** What transcriptomic foundation models encode, and how well they describe cells and tissue on targeted spatial panels, is under-explored. It is unclear whether a concept-labeled sparse autoencoder (SAE) feature represents its concept or only the genes it detects.

**Results:** Here, we propose interpreTFM, which decomposes frozen transcriptomic foundation model activations with SAEs. We applied it to a CosMx lung-cancer section, four further CosMx donors, and a Xenium renal-cell-carcinoma section. On held-out tokens of known genes, sparse features marked 262–593 concepts per analysis layer, and SAEs of re-initialized scGPT and Geneformer did as well. On held-out genes, concept F1 was 0.01–0.12 for every representation and lowest for sparse features. A linear probe of hidden states predicted the concepts of unseen genes, but only in trained models. Per cell, sparse features reached 66–98% of single-gene population specificity (57–110% in the donors). In the analyzed CosMx section, sparse-feature niches matched author cell types at ARI 0.34–0.42, while expression-only niches reached 0.30; in the donors, their defining concepts recurred but their composition did not. In Xenium, the niches shared by all models were recovered from expression alone. Calibrated model interface calls were at chance in CosMx and tracked transcript spillover in Xenium.

**Conclusions:** On these panels, SAE features act as concept-labeled gene detectors, while learned concept information is distributed across hidden dimensions. Concept labels should be checked against held-out genes and re-initialized models. For cells and niches, the features give named descriptions, comparable across models and patients and bounded by the measured panel.

## Introduction

Single-cell RNA sequencing, spatial transcriptomics, and multiplexed imaging measure cellular states in their tissue context and have shown how tissue organization changes in disease, from tumors^1^ and Alzheimer’s disease^2^ to myocardial infarction^3^ and neuropathic pain.^4^ Each technology, however, provides only a partial, context-dependent view of the underlying regulatory mechanisms, and benchmarks of spatial clustering, alignment, integration, deconvolution, and cell-type annotation methods find that performance varies considerably across platforms and tissues.^5–8^ Methods that align and jointly represent tissue sections,^9–11^ integrate spatial multi-omics modalities,^12, 13^ or deconvolve spot-level measurements with single-cell references^14–16^ address parts of this problem by transferring information across sections, modalities, and references, but integrating fragmented signals remains difficult when measurements differ in gene coverage, noise structure, and spatial context. Pretrained foundation models promise such transfer at scale, which makes it necessary to know what biological information their representations encode and whether it still describes cells and tissue after transfer to a new kind of measurement.

Transcriptomic foundation models (TFMs) are large neural networks trained on millions of single-cell profiles to learn representations that transfer across datasets and tasks. Earlier representation-learning approaches include variational models such as scVI^17^ and totalVI,^18^ which learn low-dimensional embeddings for denoising and integration. More recently, transformer-based approaches such as Geneformer,^19^scGPT,^20^ CellPLM,^21^ SATURN,^22^ and C2S-scale^23^ have added large-scale pretraining, with contextual gene representations, relational modeling of cells, cross-species embeddings, and larger models. They are meant to provide reusable embeddings for cell annotation, perturbation modeling, and cross-cohort generalization. Yet a model that performs well does not show what it has captured. A model can annotate cell types accurately without revealing which biological programs it represents, in which layer they sit, or whether they still hold on a targeted spatial panel rather than the scRNA-seq data it was trained on.

Despite this progress, three barriers limit the practical utility of TFMs. First, reported performance often depends on fine-tuning, and independent studies report that frozen models sometimes fail to outperform simple baselines and can be sensitive to batch effects.^24, 25^ Second, targeted spatial panels introduce substantial domain shift relative to scRNA-seq pre-training corpora. Spatial measurements differ in sparsity, dynamic range, noise processes, and biological context, challenging model transfer unless the evaluation procedures explicitly account for such shifts. Third, the internal representations of TFMs remain largely opaque. Individual hidden dimensions need not correspond to individual biological processes, so it is difficult to determine whether model representations carry reusable biological structure or statistical features that serve only a particular downstream task.^24^ An evaluation therefore needs simple baselines for the frozen models, tests on the targeted panels themselves, and units of the representation, at each layer, that can be read biologically.

Mechanistic interpretability, developed mainly for language and vision models, offers tools for this problem. A central idea in that work is superposition:^26–29^ neural representations can encode more features than their dimensionality permits as independent directions, so individual dimensions may mix several concepts. Sparse dictionary learning, implemented through sparse autoencoders (SAEs,^30, 31^), can partially separate such mixtures^32^ by projecting dense activations into a higher-dimensional, sparse representation in which individual latents may align more selectively with interpretable concepts. SAEs learn a dictionary {*d*_*i*_} and sparse activations *f* (*x*) such that *x* ≈ *b* + ∑_*i*_ *f*_*i*_(*x*)*d*_*i*_, providing a decomposition of dense model states into a sparse set of learned features. These ideas have clarified internal computations in language and vision models and have enabled circuit-level analyses.^28, 33^ Recent work has applied SAEs to single-cell foundation models, reporting biologically annotated and steerable features,^34, 35^ comparing SAE concepts with individual neurons,^36^evaluating features against known concepts,^37^ and testing what spatial foundation models learn.^38^ A recent preprint built a layer-resolved atlas of Gene-former and scGPT features and annotated 29–59% of them by enrichment tests on each feature’s own top 20 genes;^39^ the labels were not tested on held-out genes, and no raw-neuron or untrained-SAE baseline was reported, so such annotations cannot distinguish concept detection from gene detection. To our knowledge, no evaluation has held out whole genes during feature selection or compared trained with untrained SAEs of the same width. This leaves two questions open: whether a concept-labeled sparse feature represents its concept or only the genes it detects, and how well such features describe cells and tissue organization on targeted spatial panels compared with the measured expression.

Here we introduce interpreTFM, a layer-resolved framework that examines the internal representations of transcriptomic foundation models and places them in tissue context. We decompose frozen hidden states with SAEs and evaluate the resulting features at two linked resolutions, one for each question. At the gene-token level, we test whether sparse features align with panel-observable ontology and pathway concepts on held-out genes, whether features from randomly reinitialized models do the same, and whether a linear readout of the hidden states recovers concept information about unseen genes. At the cell level, we ask whether the resulting concept-associated representations recover spatially coherent tissue organization and localize interpretable programs to cell–cell interfaces, compared with references built from the measured expression.

interpreTFM evaluates interpretability layer by layer. We extract layer-resolved activations from frozen scGPT, Geneformer, and C2S-scale models and train a separate sparse autoencoder for each model-layer combination. Every layer is scored with three measures. The transcriptomic interpretability score (TIS) tests against shuffled controls whether a hidden dimension preserves transcriptomic structure. Held-out concept alignment tests whether a sparse feature marks the genes of a named biological concept, against raw hidden dimensions and an untrained SAE of the same width, both for new observations of known genes and for genes never seen during feature selection. Population specificity, measured by the Gini specificity index (GSI) and marker-gene hit rate (MHR), tests whether a feature is as specific to one cell population as a marker gene, against raw hidden dimensions and single genes.

We apply this framework to targeted spatial transcriptomic sections from CosMx lung cancer and Xenium renal cell carcinoma (RCC), providing distinct tests of transfer across tissue context and panel composition. Using representative layers chosen in these screens, we project retained concept-associated features to cells and use them for spatial niche discovery. We then evaluate niche structure by spatial coherence, cell-type composition, block-level robustness, niche-associated concepts, and broad cell-group interfaces. Next, we place model-free ligand–receptor colocalization in the model niches and test the models’ own programs at cell–cell interfaces with a transcript-count-adjusted statistic calibrated on contact-free fields. Finally, we ask whether a macrophage’s sparse profile predicts the composition of its neighborhood as well as its measured expression, and whether the models’ tumor niches give a consistent reading of tumor immune infiltration. One section from each of four further CosMx donors, processed with the pipeline unchanged, tests whether the results recur across patients.

Across both datasets, sparse features behave as detectors of individual genes that inherit those genes’ concept labels: the labels do not extend to genes held out during selection, although a linear readout of the hidden states still recovers concept membership for such genes. At the cell and niche level, the features approach marker-gene specificity, give niches that agree across models, and describe cells and niches about as well as the measured expression, without exceeding it; in four further CosMx donors, the cell-level results and the niches’ concept-level definition recurred, but niche composition did not. Within the niches, model-free ligand–receptor co-localization places candidate contacts in tissue context, whereas the models’ own interface calls, once calibrated, are at chance level in CosMx and follow segmentation spillover in Xenium. interpreTFM makes these tests (held-out genes, untrained and re-initialized controls, and a calibrated interface statistic) available as an open-source, model-agnostic pipeline.

## Results

### A model-agnostic workflow converts foundation-model states into spatially interpretable programs

interpreTFM is a model-agnostic framework that connects the internal representations of transcriptomic foundation models to interpretable spatial tissue organization (Fig. 1). Layer-specific hidden activations are extracted from frozen scGPT, Gene-former, and C2S-scale models given spatially resolved single-cell expression profiles (Fig. 1a,b), and the transcriptomic interpretability score (TIS) tests whether they preserve coherent transcriptomic structure (Fig. 1c). A sparse autoencoder (SAE) trained per model layer decomposes the activations into sparse latent features (Fig. 1d), which are scored at the gene-token level for held-out associations with concepts observable on the measured gene panel, on new observations of known genes and on un-seen genes, against raw hidden dimensions and an untrained SAE, and at the cell level for population specificity against single genes (Fig. 1e). Selected concept-associated features are projected back to cells, giving a single-cell representation that carries concept labels and spatial coordinates (Fig. 1f) and supports three downstream analyses: interpretable features across layers, spatial niche discovery, and localization of interpretable programs at cell–cell interfaces (Fig. 1g-i). Cell-type labels enter only down-stream, for calibration and interpretation, and are not used to train the sparse features or to define their concept associations.

**Figure 1.**
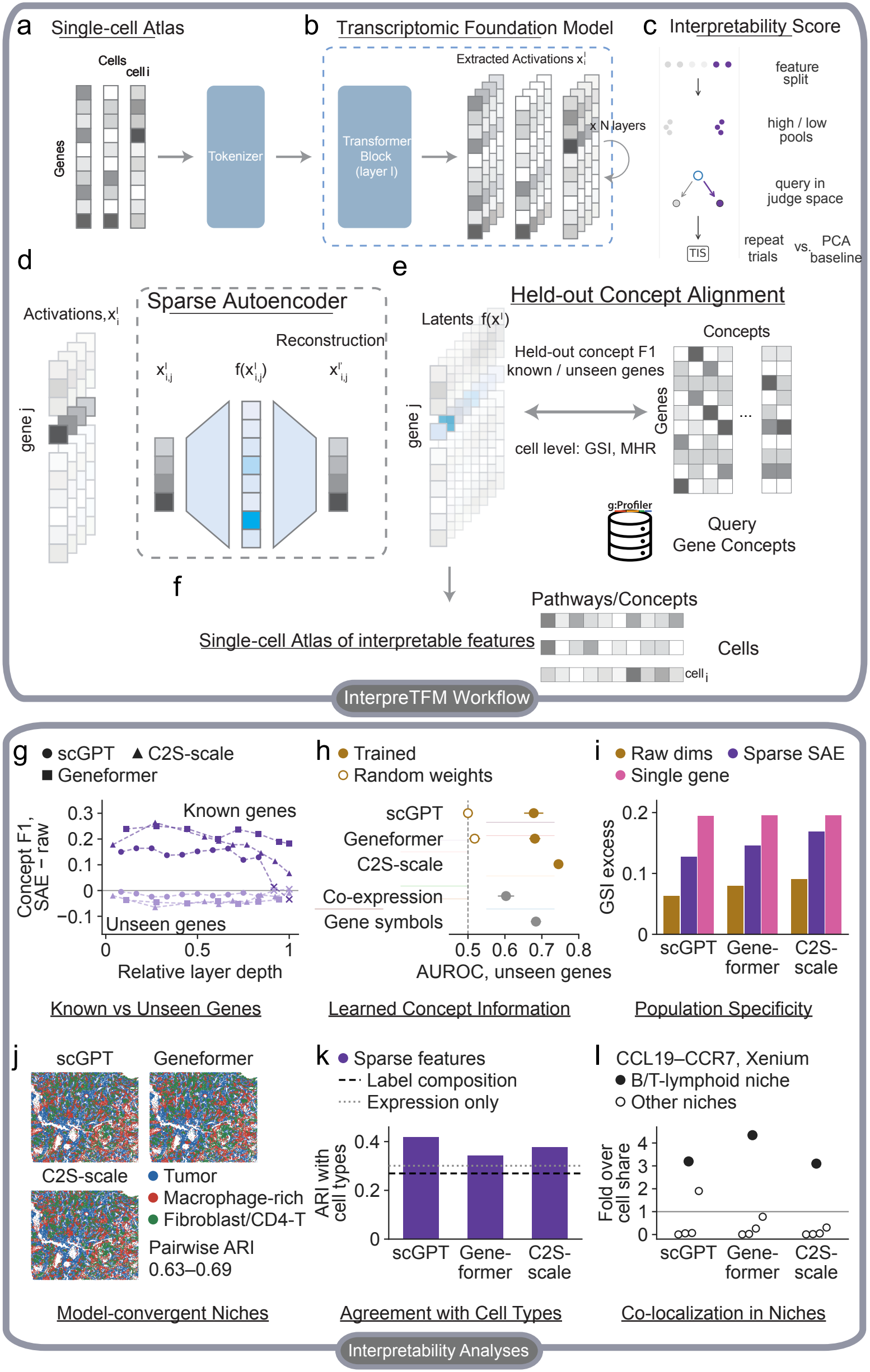
The interpreTFM workflow links foundation-model activations to interpretable spatial biology. **a**, Spatially resolved single-cell expression profiles and coordinates. **b**, Token-level hidden activations from selected layers of frozen transcriptomic foundation models. **c**, Transcriptomic interpretability score: whether cells with high and low activation of a hidden dimension differ coherently in measured expression, against shuffled and PCA controls. **d**, A separate sparse autoencoder (SAE) per model layer decomposes hidden states into sparse latent features. **e**, Features are scored by held-out gene-token association with panel-observable pathway and ontology concepts, on new observations of known genes and on unseen genes, against raw hidden dimensions and an untrained SAE, and at the cell level for population specificity against single genes. **f**, Selected concept-associated features are projected back to cells. **g–l**, Main results (CosMx lung-cancer section unless stated). **g**, Concept F1 of SAE features minus raw hidden dimensions per layer, on known genes (row split) and unseen genes (whole-gene split); crosses, layers with fewer than 10 retained features. **h**, Area under the ROC curve (AUROC) of a linear probe for the concept membership of unseen genes, for trained and randomly re-initialized models and for co-expression and gene-symbol baselines; dashed line, chance. **i**, Gini specificity index (GSI) excess of raw hidden dimensions, SAE features, and single genes. **j**, Niche maps of the three models and their pairwise adjusted Rand index (ARI). **k**, Niche–cell-type ARI against label-composition (dashed) and expression-only (dotted) niches. **l**, CCL19–CCR7 close pairs in the Xenium B/T-lymphoid niche (filled) and the other niches (open), as fold over the niche’s cell share. Details in Figs. 2, 3, and 5 and Supplementary Table S3. Cell-type labels are not used to train the sparse features (Methods).

### Sparse features act as concept-labeled gene detectors, and learned concept information about unseen genes is spread across hidden dimensions

We first asked whether a sparse feature labeled with a biological concept represents that concept or only the genes it detects, and scored every model layer (Fig. 2; Methods). TIS gain measures how coherently cells split by a hidden dimension differ in measured expression, beyond shuffled activations. Held-out concept alignment tests whether activation of a single sparse feature predicts membership of the corresponding gene in a pathway or ontology term, scored on held-out gene-token rows of genes seen during selection (row split) or on genes never seen (whole-gene split), with raw hidden dimensions and an untrained SAE of the same width as matched baselines.

**Figure 2.**
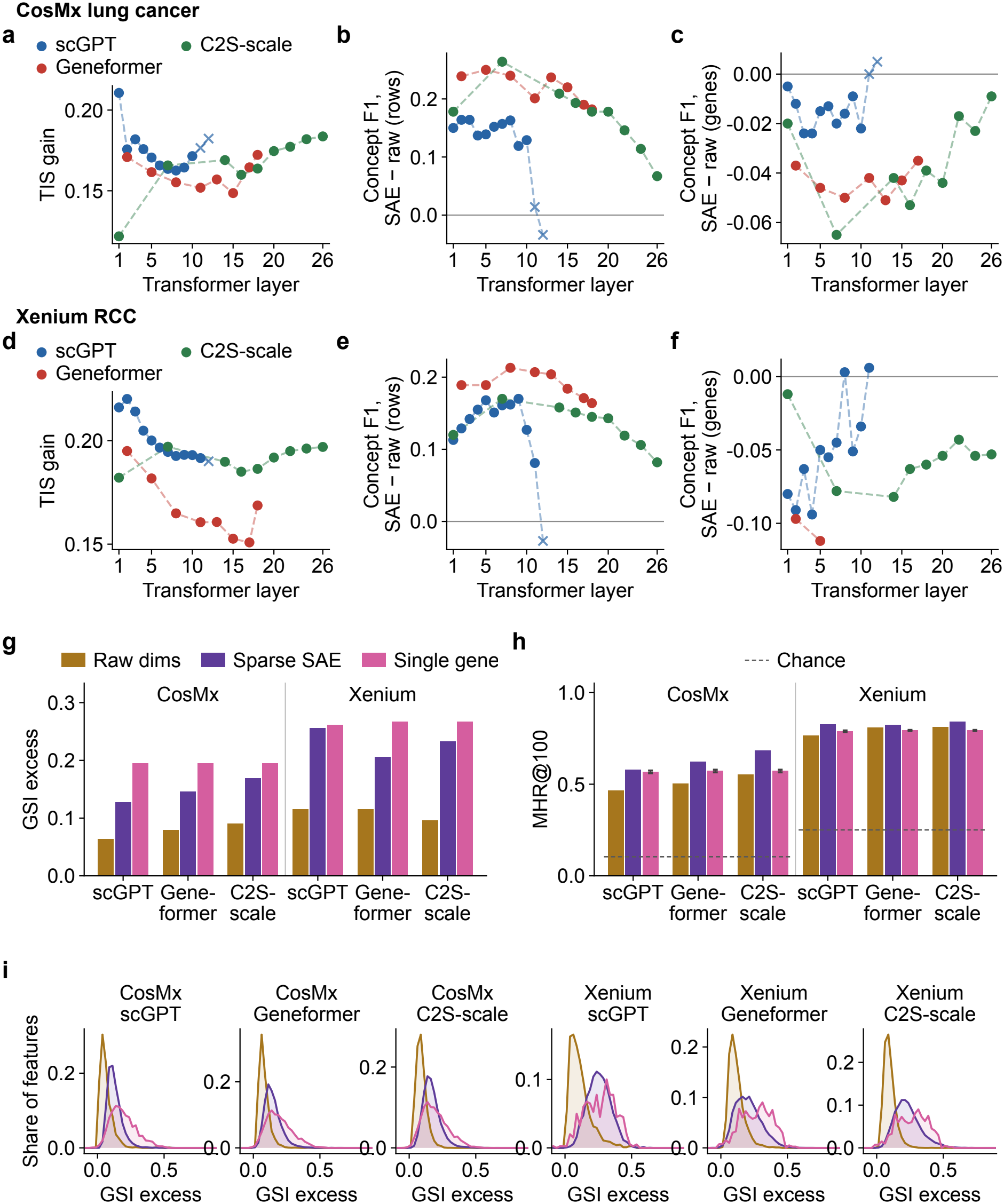
Concept alignment of sparse features depends on the held-out unit, and their cell-level specificity approaches, and in CosMx can exceed, that of single genes. **a–c**, CosMx layer screen for scGPT, Geneformer (GF), and C2S-scale (C2S): TIS gain, and concept F1 of sparse autoencoder (SAE) features minus raw hidden dimensions with held-out gene-token rows (row split) and held-out genes (whole-gene split). **d–f**, Xenium screen; whole-gene results are available for 50 of the 58 layers (not for CosMx Geneformer Layer 18, Xenium scGPT Layer 12, or Xenium Geneformer layers other than 2 and 5). Crosses, layers with fewer than 10 features retained by the spatial feature rule; lines connect layers within a model only. **g–i**, Population specificity at the representative layers for raw hidden dimensions, sparse SAE features, and single panel genes on the same cells: **g**, GSI excess averaged over active features; **h**, MHR@100 (dashed lines, chance rate; error bars on single genes, SD over five additional splits); **i**, distributions of per-feature GSI excess. Values in Supplementary Table S1. TIS, transcriptomic interpretability score; GSI, Gini specificity index; MHR, marker-gene hit rate.

The hidden states carried transcriptomic structure throughout model depth (Fig. 2; Supplementary Table S1): TIS exceeded independently shuffled controls at every layer, with mean gains of 0.122–0.211 in CosMx and 0.151–0.220 in Xenium across the three models, and the gain varied substantially with depth. TIS depends on the basis of the activations: the same activations rotated to principal components gave gains of only 0.008–0.047. The models’ cell embeddings at the analyzed layers separated the annotated cell types well above chance (balanced k-nearest-neighbor accuracy 0.28–0.44 in CosMx and 0.58–0.78 in Xenium, against 0.05 and 0.08 by chance), at or below the log-normalized expression of the same cells (0.44, and 0.88 for Xenium labels derived from that expression; Methods).

Concept alignment at the layer level was summarized as concept F1, the held-out F1 of the feature selected on validation data averaged over concepts with 5–100 panel genes (Methods). With held-out gene-token rows, sparse SAE features exceeded raw hidden dimensions at 56 of 58 evaluated layers (median difference 0.16; Fig. 2b,e) and an untrained SAE at all six analysis layers (the representative layers used downstream, one per model and dataset; see below), where 63–111 sparse features each marked the member genes of at least one concept with a test F1 of at least 0.5, together covering 262–593 concepts, against 0–70 concepts for raw hidden dimensions and 0–249 for the untrained SAE (Supplementary Table S1).

On genes held out during selection, concept F1 was 0.01–0.12 for every representation: sparse features scored below raw dimensions at 43 of 50 layers (by up to 0.11) and were the lowest of the three representations at all six analysis layers (Fig. 2c,f; Supplementary Table S1), and features selected for a concept fired on the specific genes seen during selection (CosMx C2S-scale Layer 14: validation F1 0.618, 0.025 on unseen genes). Row-level concept alignment reflects how well a dictionary isolates individual genes: a feature inherits its concept label from the genes it detects. Consistent with this, sparse SAEs trained on scGPT and Geneformer with randomly re-initialized weights reached the same row-split concept F1 as on the trained models (scGPT 0.330–0.366 versus 0.333; Geneformer 0.341–0.349 versus 0.355; CosMx Layer 2, the same cells for every form) and marked as many or more concepts with a test F1 of at least 0.5 (scGPT 350–643 versus 275; Geneformer 636–638 versus 559; Methods; Supplementary Table S1). In a steering check, adding a concept-labeled feature’s decoder direction to the hidden states moved the model’s output for the concept’s other member genes no more than the detected gene’s own identity direction did in any of the three models (differences *−*0.008 to +0.003), although more than a random feature in C2S-scale and scGPT (+0.056 and +0.030; Methods; Supplementary Table S2).

Concept information was nevertheless present in the representations: a linear probe on the mean hidden states of training genes ranked unseen genes by concept membership with AUROC 0.66–0.75, against 0.49–0.50 for random vectors and 0.59–0.61 for same-section co-expression and cell-type expression profiles, whereas the best single dimension reached at most 0.63, so the information is spread across dimensions (Supplementary Table S3). This information was learned in pretraining: with every weight randomly re-initialized, the readout of scGPT and Geneformer fell to chance (0.50 and 0.52; CosMx Layer 2), whereas re-initializing every weight except the gene-embedding table left it nearly unchanged (0.67 and 0.66), so at this layer it resides mainly in the learned gene embeddings (post hoc). Gene symbols carry comparable information, because many concepts group gene families: a readout of each symbol’s character trigrams reached 0.68 in CosMx and 0.64 in Xenium, and removing the part of each hidden state predictable from the symbols lowered every model’s readout by 0.06–0.09, that of C2S-scale, which reads gene names, slightly more in CosMx (0.088 versus 0.062–0.065; post hoc; Supplementary Table S3). Readouts of the sparse features were lower with standardized inputs (0.59–0.67); with centered inputs (post hoc, because standardization inflates rare sparse activations), they matched the raw hidden state within 0.01 at three analysis layers and were 0.025–0.033 lower at the other three, so the sparse decomposition partly preserved this information.

### Averaged per cell, sparse features approach the specificity of marker genes, and no single layer is optimal

Gene detectors can still describe cells, because cell populations are defined by the genes they express. For the second question, how well the features describe cells and tissue, the measured expression sets the upper reference, because the features are computed from the measured genes and cannot carry more information than the panel. We therefore asked how much of that information the features keep in named units, and how reproducible those units are across models, seeds and patients. We started with population specificity: whether a feature, averaged per cell, is as specific to one cell population as a marker gene (GSI excess and MHR@100, against raw hidden dimensions and single genes). At the cell level, sparse features were more population-specific than raw hidden dimensions (GSI excess 0.13–0.26 versus 0.06–0.12) and had higher marker-gene hit rates at all six analysis layers (Fig. 2g–i; Supplementary Table S1). Against single panel genes, the natural reference because populations are defined from expression, sparse features reached 66–98% of single-gene specificity (raw hidden dimensions, 32–47%) and slightly exceeded single-gene hit rates in Xenium (0.82–0.84 versus 0.79), where MHR@100 is near its ceiling for every representation; in CosMx the leading C2S-scale and Geneformer features recovered marker programs more completely than the best single genes (MHR@100 0.69 and 0.62 versus 0.57). In one section from each of four further donors of the same CosMx study, processed with the pipeline unchanged (Methods), sparse features again exceeded raw hidden dimensions on both measures in all 12 donor– model combinations and reached 57–110% of the single-gene GSI excess (raw hidden dimensions, 32– 49%), and C2S-scale features exceeded single genes on MHR@100 in every donor (Supplementary Table S4).

Different criteria favored different depths: row-level concept F1 was highest in early-to-intermediate layers, marker-gene hit rate peaked at C2S-scale Layer 14 in both datasets, and GSI tended to rise with depth (Fig. 2; Supplementary Table S1). We therefore did not re-select layers with a composite of these criteria; downstream analyses use representative layers chosen in a preliminary screen (a composite of TIS gain and concept alignment) and kept fixed throughout (CosMx scGPT Layer 2, Geneformer Layer 2, and C2S-scale Layer 14; Xenium scGPT Layer 3, Gene-former Layer 2, and C2S-scale Layer 14).

### Interpretable representations recover coherent, model-convergent niches in a lung tumor section, with tumor and stromal concepts recurring across patients

With the representative layers fixed, we went from single cells to tissue and asked whether cell representations built from the sparse features recover tissue organization, and how they compare with niches built from the measured expression. All three CosMx representations partitioned the section into the same three spatially coherent niches: a tumor niche (tumor 13, 84–89% of cells), a macrophage-rich niche (macrophage 49–52%, CD4 memory T 11–13%, fibroblast 10–14%), and a fibroblast/CD4-T stromallymphoid niche (fibroblast 33–40%, CD4 memory T 21–23%, pDC 5–6%) (Fig. 3; Supplementary Figure S1 and Table S5). Niche-versus-cell-type adjusted Rand index (ARI) was 0.419, 0.342, and 0.377 for scGPT, Geneformer, and C2S-scale, above both a label-composition baseline (0.269) and an expression-only baseline on the same sweep grid (0.301); on normalized mutual information (NMI) only Gene-former (0.292) fell below the expression-only winner (0.317). The partitions were stable and spatially contiguous (seed-to-seed ARI ≥0.989; neighborhood purity 0.82–0.85 versus 0.61–0.63 under the tile-shuffle null) and agreed with each other (ARI 0.63–0.69) more than with expression-only niches at their own settings (mean 0.665 vs 0.471). The convergence is onto a coarse tumor/myeloid/stroma-lymphoid split: each partition also agreed with the label-composition baseline at ARI 0.46–0.52, and expression-only niches at the three settings agreed with each other at 0.74–0.85. At finer resolution (four to five niches), model-feature niches were more reproducible across seeds than expression-only niches, both in this section (median seed ARI 0.78–0.91 vs 0.71) and in all 12 donor– model combinations of the four further donors (Supplementary Table S4e). The concept labels of the niche-defining features concurred across models for niches matched by composition (top-25 concept-label Jaccard 0.14–0.22 versus 0.014–0.021 for unmatched niches), although at most 11 of 25 labels were shared in any matched niche (Supplementary Note S1).

**Figure 3.**
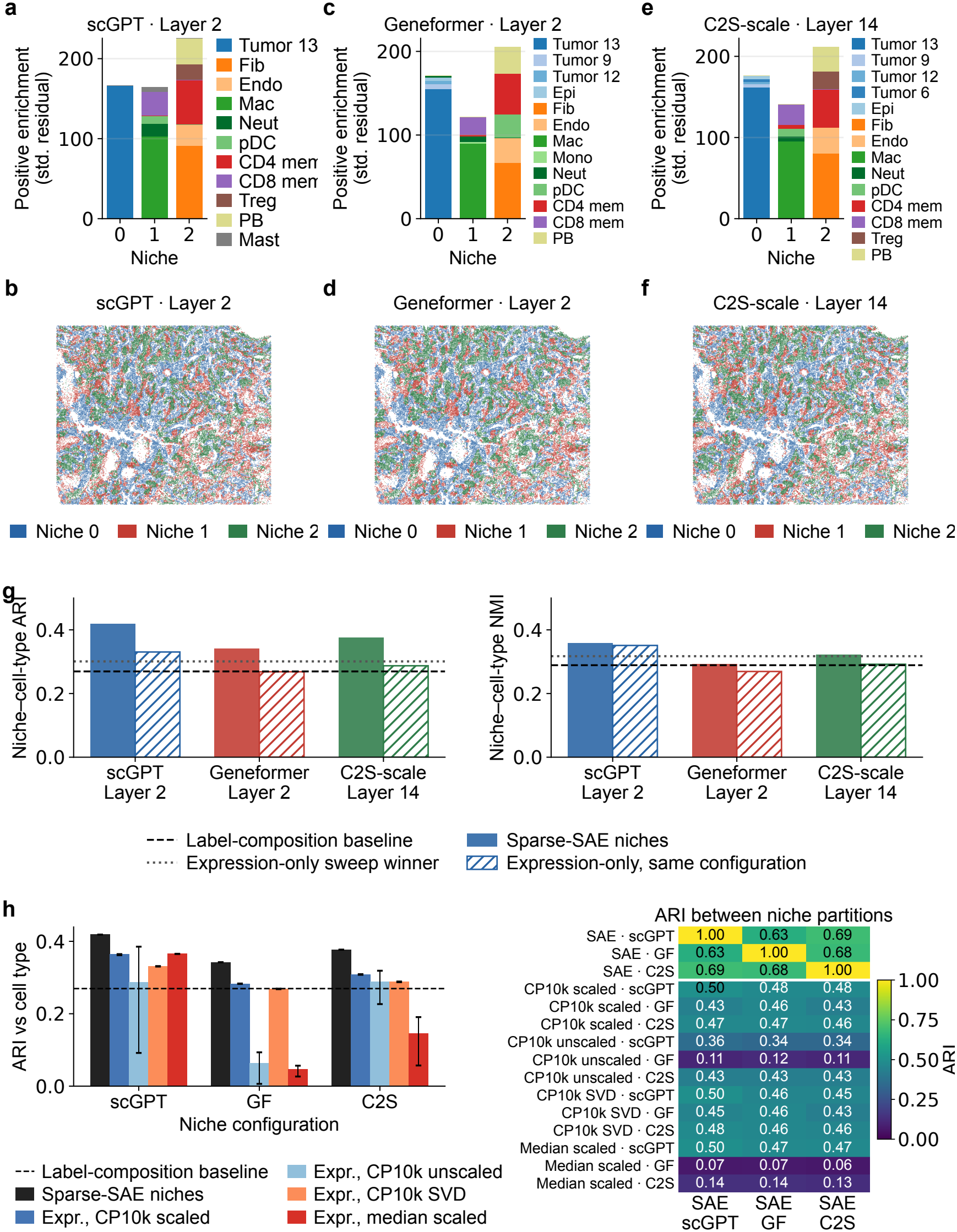
Interpretable representations recover spatially coherent, model-convergent CosMx lung-cancer niches. **a,b**, scGPT Layer 2; **c,d**, Geneformer Layer 2; **e,f**, C2S-scale Layer 14, at the sweep-selected configurations (Methods). Bars (a,c,e), the five largest positive standardized cell-type residuals per niche above 1.5; stack heights summarize residuals, not cell-type proportions. Maps (b,d,f), niche of each cell. Signed residual heatmaps in Supplementary Figure S1. **g**, Niche-versus-cell-type adjusted Rand index (ARI) and normalized mutual information (NMI) of each model and of expression-only niches at the same configuration; horizontal lines, label-composition baseline (Squidpy, *k* = 3) and expression-only sweep winner. **h**, ARI of sparse-feature and expression-only niches (four expression preprocessings at the same configurations) with author cell types (left) and with each other (right); bars, mean over clustering seeds 0, 1, and 2 (the sparse-feature bar is the single selected partition); whiskers, range; dashed line, label-composition baseline; color limits from 0 to the largest |ARI| (1, the self-comparisons), as in Fig. 4i. Tumor 6, 9, 12, and 13, author tumor clusters 6, 9, 12, and 13; Epi, non-tumor epithelial; Fib, fibroblast; Endo, endothelial; Mac, macrophage; Mono, monocyte; DC, dendritic cell; pDC/mDC, plasmacytoid/myeloid DC; Neut, neutrophil; Mast, mast cell; B/Pl, B/plasma; PB, plasmablast; NK, NK cell; Treg, regulatory T; CD4/CD8 mem and naive, memory and naive CD4/CD8 T cells; GF, Geneformer; C2S, C2S-scale.

The niches’ cell-type composition was specific to this section. With the pipeline applied unchanged to one section from each of four further donors, each niche’s cell-type residual profile matched its best donor counterpart at cosine 0.18–0.72, against 0.98–1.00 for the same niche across the three models within this section (post hoc); with the per-donor tumor subtypes merged, the tumor niche came closer (0.79–0.92) and the other two niches varied by donor (0.43–0.91). Expression-only and label-composition niches were as section-specific (0.02–0.80; post hoc), so section specificity is not particular to the foundation models, and in the donors the agreement of model niches with author cell types exceeded that of expression-only niches clearly only in Lung6 (two of three models), was equal in Lung5, and was equal or lower in Lung9 and lower in Lung12 (Supplementary Table S4b). The niches’ concept-level definition did recur: for the tumor and stromal-lymphoid niches, a donor niche shared top-25 concept labels with this section above chance (*P <* 0.05 against random concept sets) in all four donors and all three models (Jac-card 0.28–0.72 vs 0.16–0.18 expected; Supplementary Table S4f); the macrophage-rich niche did so in all four donors for Geneformer but in two for scGPT and C2S-scale.

### Interpretable representations recover three RCC territories that are shared across models and present in the measured expression

We repeated the niche analysis in the Xenium RCC section, a more stringent test on a second platform and tissue: a much larger section, a smaller panel, and no tumor-aware annotation (Fig. 4; Supplementary Tables S5 and S6). Kidney-lineage labels transferred from a tumor-naive reference were supported by their own panel markers only for the broad compartments (own marker in 85–95% of endothelial, immune, and smooth muscle/pericyte cells), whereas the tubule-subtype labels were contradicted (own marker in 0.2–1.8% of thick ascending limb (TAL), principal (PC), and intercalated (IC) cells) or untestable (Supplementary Table S7). Cells were therefore annotated from the panel (13 marker-based types; 8.7% low quality; Methods; Supplementary Figure S2), with the transferred labels as a secondary set. Each representation was partitioned into five niches (Methods), which agreed only weakly with either label set, no more than expression-only niches or the label-composition baseline did (Supplementary Table S5).

**Figure 4.**
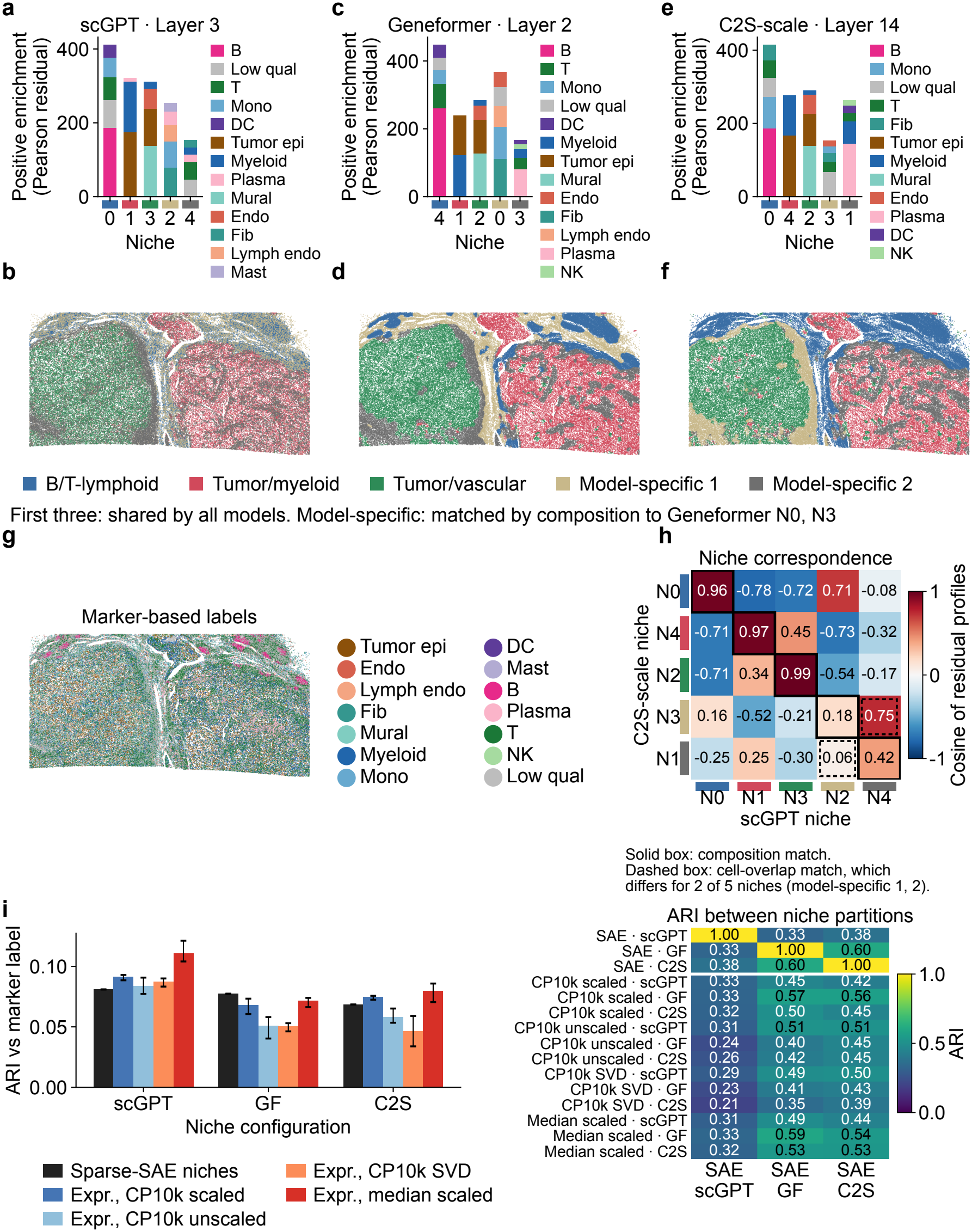
Model-derived niches resolve recurrent territories in a Xenium renal cell carcinoma (RCC) section. **a,b**, scGPT Layer 3; **c,d**, Geneformer Layer 2; **e,f**, C2S-scale Layer 14, at the best-scoring five-niche configurations (Methods), showing positive enrichment residuals (five largest per niche at ≥ 1.5) and the niche map. Labels are marker-based (Methods; Supplementary Figure S2). Niche numbers are model-specific; niches are ordered and colored by matched group: the three shared niches (B/T-lymphoid, tumor/myeloid, tumor/vascular) have the same color in every model, and each model’s two other niches are shown in neutral colors (model-specific 1 and 2) according to their composition match to Geneformer niches 0 and 3. **g**, Whole-section orientation by marker-based label. **h**, Cosine correspondence of C2S-scale and scGPT niche residual profiles; solid boxes, composition matching; dashed boxes, cell-overlap matching, which pairs the two model-specific niches the other way; expression signatures per niche in Supplementary Figure S3. **i**, Adjusted Rand index (ARI) of foundation-model and expression-only niches with the marker-based labels (left) and with each other (right); bars, mean over clustering seeds 0, 1, and 2 (the foundation-model bar is the single selected partition); whiskers, range; color limits as in Fig. 3h. Three-niche solutions in Supplementary Figure S4. Tumor epi, tumor-like epithelial; Endo, endothelial; Lymph endo, lymphatic endothelial; Fib, fibroblast; Mono, monocyte; DC, dendritic cell; Mast, mast cell; B, B cell; Plasma, plasma cell; T, T cell; NK, NK cell; Low qual, low quality (*<*20 transcripts or *<*10 genes); GF, Geneformer; C2S, C2S-scale.

Whole-section maps, marker-label residuals, and expression signatures (Supplementary Figure S3) nevertheless identified the same three shared niches in every model: a B/T-lymphoid niche (T cells 27–32%, B cells 16–25%, monocytes 7–9%; B/lymphoid-follicle signature), a tumor/myeloid niche (tumor-like epithelial 26–35%, myeloid 31–34%; epithelial, proliferation, IFN-*γ*-chemokine, and macrophage signatures), and a tumor/vascular-mural niche (tumor-like epithelial 23–26%, mural 16–18%, endothelial 10–12%; vascular and stromal signatures). Each shared niche drew at least 87% of its cells from one territory of the three-niche solutions (Supplementary Figure S4), and matched shared niches had residual-profile cosines of 0.91–0.99 in every model pair. The other two niches per model resolved substructure that differed between models: a fibroblast/monocyte/lymphatic-endothelial niche at the lymphoid border (scGPT, Geneformer), a plasma-rich niche (plasma 9–15%; Geneformer, C2S-scale), and a T-cell-rich niche spanning two territories (scGPT, C2S-scale); matched model-specific niches had cosines of 0.18–0.88, and the partitions agreed at ARI 0.33–0.60. The partitions were stable (seed-to-seed ARI ≥0.996) and, at four to five niches, more reproducible across seeds than expression-only niches (median 0.88–0.93 vs 0.69), as in CosMx; the Geneformer and C2S-scale niches were spatially contiguous (neighborhood purity 0.88 and 0.87 versus 0.68 and 0.70 under the tile-shuffle null). The expression-only five-niche solution contained the same three shared niches (best-match residual cosine 0.88–0.99; post hoc; Supplementary Table S5), so these territories are a property of the section’s expression that the sparse features preserve.

The marker-based labels come from one clustering of a 399-gene panel without pathology review, so “tumor-like epithelial” does not establish malignancy. The scGPT niches were barely more contiguous than under the tile-shuffle null (purity 0.53 vs 0.48), its T-cell-rich niche being interspersed through both tissue lobes. The shared niches occupy the two lobes and the strip between them, and the tumor/myeloid niche has more transcripts per cell than the other niches (median 119–181 vs 43–90): a partition by transcript-count tercile agreed with the niches at ARI 0.116, 0.048, and 0.060 (scGPT, Geneformer, C2S-scale), against 0.081, 0.077, and 0.068 for the marker-based labels, so transcript depth accounts for at least as much of the scGPT partition as cell type, though not of the other two; all six values are below 0.12.

### Model niches place ligand–receptor co-localization in tissue context, whereas the models’ calibrated interface calls show no detectable contact-specific signal

From territories we turned to contacts between cells. We next asked where ligand- and receptor-expressing cells meet within the model niches of both sections, and whether the models’ own programs mark those contacts (Figs. 5 and 6). Co-localization was measured without the models: for every CellChatDB ligand–receptor (LR) pair with all genes on the panel (397 CosMx, 25 Xenium), we counted ligand-positive–receptor-positive cell pairs within 100 coor-dinate units (about 18 µm in CosMx, contact range; 100 µm in Xenium, neighborhood range) and compared them with a null that permutes gene positivity within each cell type, so that the layout of the annotated cell types cannot by itself produce a hit (Methods; Supplementary Table S12). The models supplied the niche context, and their programs were tested at cell-group interfaces with a calibrated statistic.

**Figure 5.**
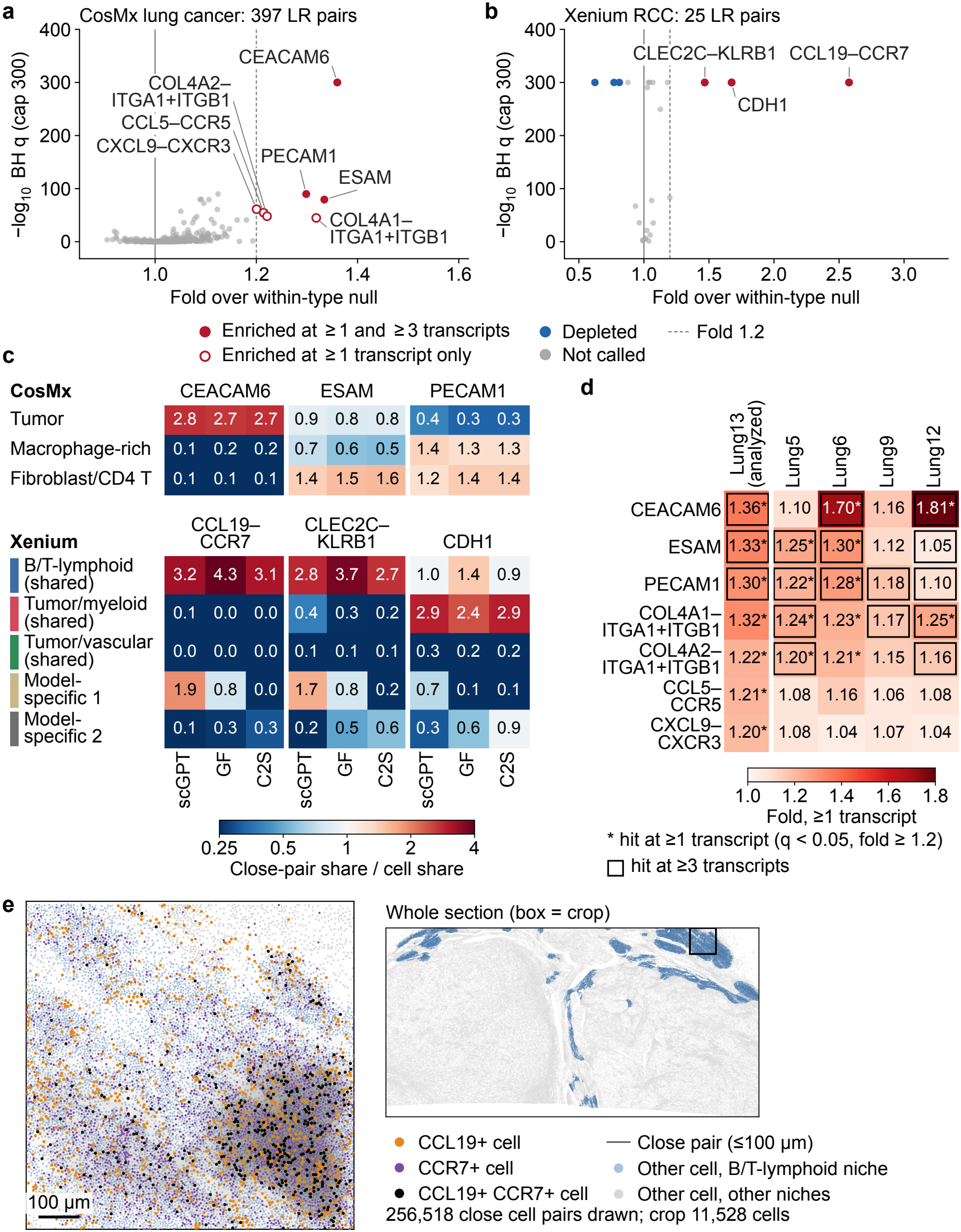
Model niches place ligand–receptor co-localization in tissue context. **a,b**, Model-free co-localization screen of every CellChatDB ligand–receptor (LR) pair with all genes on the panel (**a**, CosMx lung cancer, 397 pairs; **b**, Xenium renal cell carcinoma (RCC), 25 pairs): fold of ligand-positive–receptor-positive close pairs (distinct cells within 100 coordinate units: about 18 µm in CosMx, 0.18-µm pixels; 100 µm in Xenium; positive, *≥*1 transcript) over a within-type null (positivity permuted within cell type, 100 permutations), against *−* log_10_ Benjamini–Hochberg (BH) adjusted *q* (capped at 300). Hits (*q <* 0.05, fold *≥*1.2) are labeled; filled, enriched at *≥*1 and *≥*3 transcripts; open, at *≥*1 transcript only (untestable at *≥*3); blue, depleted (*q <* 0.05, fold ≤1/1.2); dashed line, fold 1.2. Homophilic pairs are named by their gene (CEACAM6 = CEACAM6–CEACAM6); CLEC2C–KLRB1 is the CellChatDB name (ligand gene CD69). **c**, Niche placement of the robust hits: share of a hit’s close pairs falling in a niche divided by the niche’s share of cells (1, no enrichment), per model (scGPT; GF, Geneformer; C2S, C2S-scale). CosMx: tumor, macrophage-rich, and fibroblast/CD4-T niches. Xenium: five niches per model labeled by composition-matched group (color strip as in Fig. 4); model-specific 1 and 2 are matched by composition to Gene-former’s fibroblast/monocyte/lymphatic-endothelial and plasma-rich niches (the T-cell-rich niche is model-specific 2 in scGPT and model-specific 1 in C2S-scale). **d**, Cross-donor replication of the seven CosMx hits: fold at ≥1 transcript in the analyzed section (Lung13) and one section from each of four further donors, screen unchanged; *, hit; box, hit at ≥3 transcripts. **e**, Descriptive Xenium example: CCL19-positive (orange), CCR7-positive (purple), and double-positive (black) cells and their close pairs (gray lines; 256,518 cell pairs within the crop) in an 800×800-µm crop; other cells light blue when all three models place them in the B/T-lymphoid shared niche, gray otherwise. Crop: the 500-µm tile with the median number of close pairs among tiles with ≥300 cells and ≥50% of cells in this consensus niche, widened to 800 µm within the section. Inset, whole section (consensus niche blue; box, crop). Display only.

**Figure 6.**
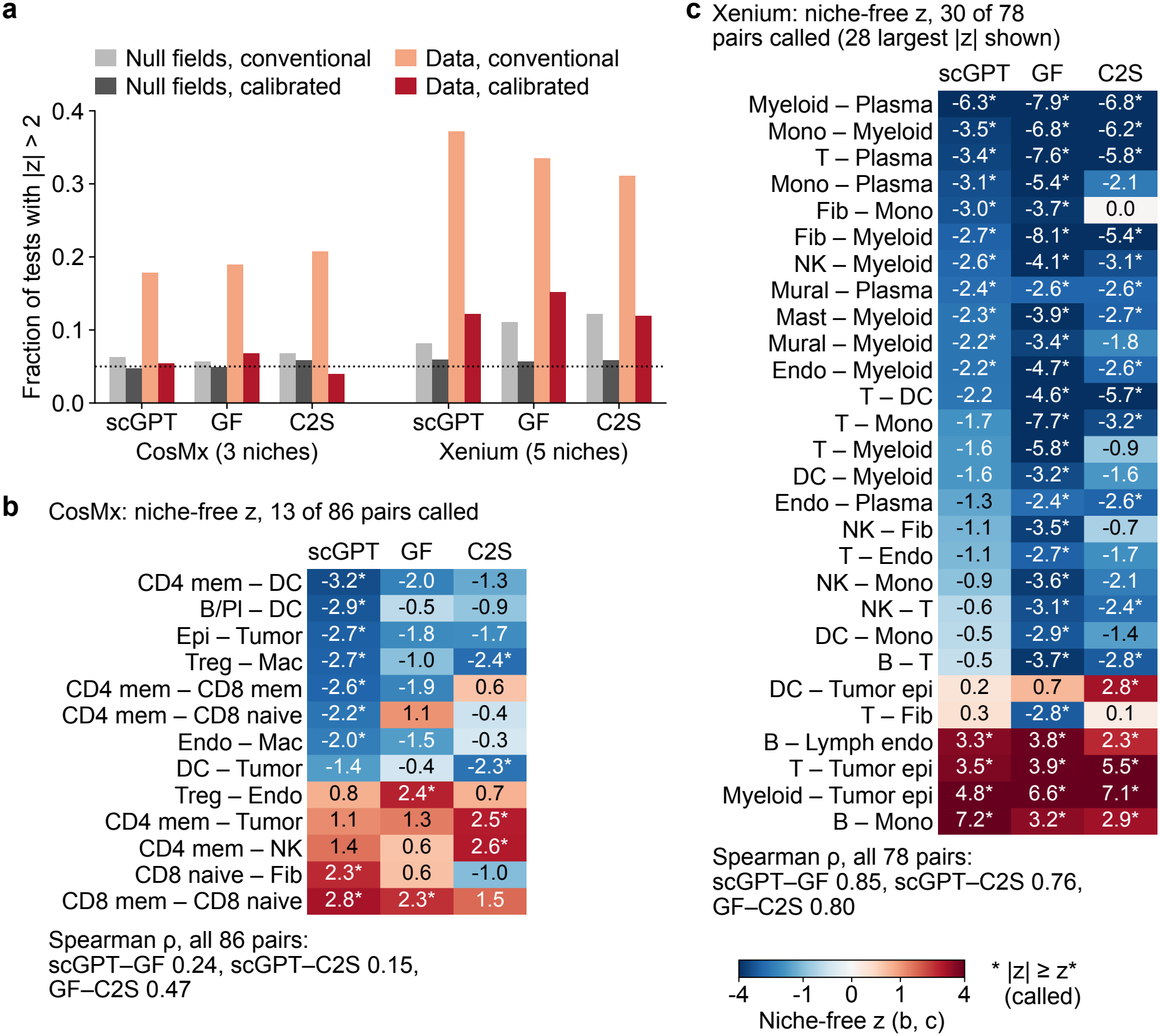
A calibrated interface test controls false-positive interface calls in both datasets. **a**, Fraction of niche-by-pair tests with |*z* |*>* 2 per model on contact-free smooth null fields matched on neighbor correlation (gray) and on the data (orange, red), under a conventional (uncalibrated) 400-unit tile null and under the calibrated 120-unit block null (CosMx 72 and 22 µm; Xenium 400 and 120 µm); CosMx at three niches (125–132 tests), Xenium at five niches (376–382 tests; marker-based labels). Dotted line, 0.05. **b,c**, Niche-free calibrated *z* of the cell-type pairs called in at least one model (**b**, CosMx, 13 of 86 pairs; **c**, Xenium, 30 of 78 pairs, the 28 with the largest |*z*| shown); *, |*z*| ≥*z*^*∗*^ (niche-free *z*^*∗*^: CosMx 2.00; Xenium 2.23–2.30); color, sign(*z*) log_10_(1 + |*z*|) with ticks in *z*. Spearman *ρ* between models over all pairs. Xenium contact-level calls track partner-marker transcripts at the contact (segmentation spillover) and are not read as interface biology. Full per-model layouts in Supplementary Figures S6– S11. Abbreviations as in Figs. 3 and 4; GF, Geneformer; C2S, C2S-scale.

In CosMx, three homophilic adhesion pairs were enriched at both transcript thresholds, CEACAM6 (fold 1.36), ESAM (1.33), and PECAM1 (1.30), and four collagen–integrin and chemokine–receptor pairs of distinct ligand and receptor genes (fold 1.20–1.32) at ≥ 1 transcript (Fig. 5a). Their close pairs concentrated in specific niches: CEACAM6 2.8-fold in the scGPT tumor niche, and PECAM1 1.3–1.4-fold in the macrophage-rich or stromal niche (Fig. 5c). In four further donors, every hit except CXCL9– CXCR3 was enriched in the same direction (fold *>* 1, *q <* 0.05); the adhesion and collagen–integrin pairs had folds of 1.05–1.81 (1.22–1.36 in this section) and the chemokine pairs 1.04–1.16 (1.20–1.21), and in two donors CEACAM6 close pairs were 3.9–5.0-fold over-represented in the tumor niche of all three models; collagen–integrin contacts were mostly between cells of the same type (Fig. 5d; Supplementary Table S13). In Xenium, CCL19–CCR7 was the strongest pair (fold 2.6; 4.5 at ≥3 transcripts), and its close pairs concentrated in the B/T-lymphoid niche (3.1–4.3 times its cell share; Fig. 5b,c,e), consistent with chemokine-guided lymphoid aggregation. CLEC2C– KLRB1 (CellChatDB name; ligand gene CD69; fold 1.5) also localized to the B/T-lymphoid niche and CDH1–CDH1 (1.7) to the tumor/myeloid niche (2.4– 2.9 times), whereas EDN1–EDNRB was not enriched section-wide (fold 1.01; see below) and two ACKR1 chemokine pairs and SELL–CD34 were depleted.

Because 100 coordinate units cover about 7 neighboring cells in CosMx but 287 in Xenium, the screen was repeated at matched neighbor counts: at 15.4 µm the three Xenium hits remained hits in the same niches, whereas at 117.5 µm the CosMx folds were lower (1.10–1.27 in this section and 1.07–1.50 in the donors), although every CosMx target stayed enriched in every donor (*q <* 0.05; Supplementary Table S20). Matched-niche CellChat pathway profiles agreed across models (mean Spearman, CosMx 0.88– 0.94 vs 0.79–0.80 for unmatched niches; Xenium 0.87– 0.91 vs 0.69–0.75; Supplementary Figure S5); CCL signaling was specific to the B/T-lymphoid niche in two of three models, and in CosMx the tumor niche carried pathways specific to it in all three (CEACAM, CDH/CDH1, EGF, ICAM, KIT, and CCL).

The models’ programs added no detectable contact-specific signal. Strict crosstalk scores the transcript-count-adjusted program on each cell-group interface of a niche against a within-stratum spatial null, with a threshold *z*^*∗*^ calibrated on contact-free smooth random fields (Methods). Under a conventional, uncalibrated 400-unit tile null, contact-free fields produced |*z*| *>* 2 in 5.7–6.8% of CosMx and 8.2–12.2% of Xenium pairs, against 4.8–5.9% under the calibrated null (Fig. 6a; Supplementary Table S9). In CosMx the calibrated statistic called 7, 9, and 5 of 125–132 niche-by-pair tests for scGPT, Gene-former, and C2S-scale, close to the 6.2–7.3 expected by chance, and no enriched pair was called by all three models (Fig. 6b; Supplementary Figures S6– S8 and Table S8). In Xenium, 42, 55, and 42 of 376– 382 tests were called, and the niche-free scores agreed across models (Spearman 0.76–0.85; Fig. 6c), whereas niche-level *z* agreed less across composition-matched niches (Spearman 0.21–0.37; Supplementary Figures S9–S11). Enrichments and depletions alike tracked partner-marker transcripts at the contact (T–tumor-like epithelial niche-free *z* −14 to −35 in the lowest vs +20 to +40 in the highest tercile; 84% of niche-level calls reversed sign between terciles), the pattern expected from segmentation spillover,^40, 41^ and removing the features that detect either partner’s markers erased the tumor-interface enrichments in all three models (Supplementary Table S11).

For case views of the interfaces, Figs. 7a and 8 combine contact-adjusted residual maps, distance diagnostics, and curated expression/proximity overlays from the same section (CosMx case crops in Supplementary Figure S12; Supplementary Tables S14–S16). The residual maps show where program burden on the primary interfaces (CosMx endothelial– fibroblast; Xenium endothelial–tumor-like epithelial) remains high or low after accounting for local pair-edge abundance (edge counts in Supplementary Table S14). Distance diagnostics against a coordinate-shuffled target-cell null showed no organization beyond contact geometry in either dataset (Supplementary Figure S13a; Fig. 8b), so they serve as the contact-geometry baseline for the residual maps.

**Figure 7.**
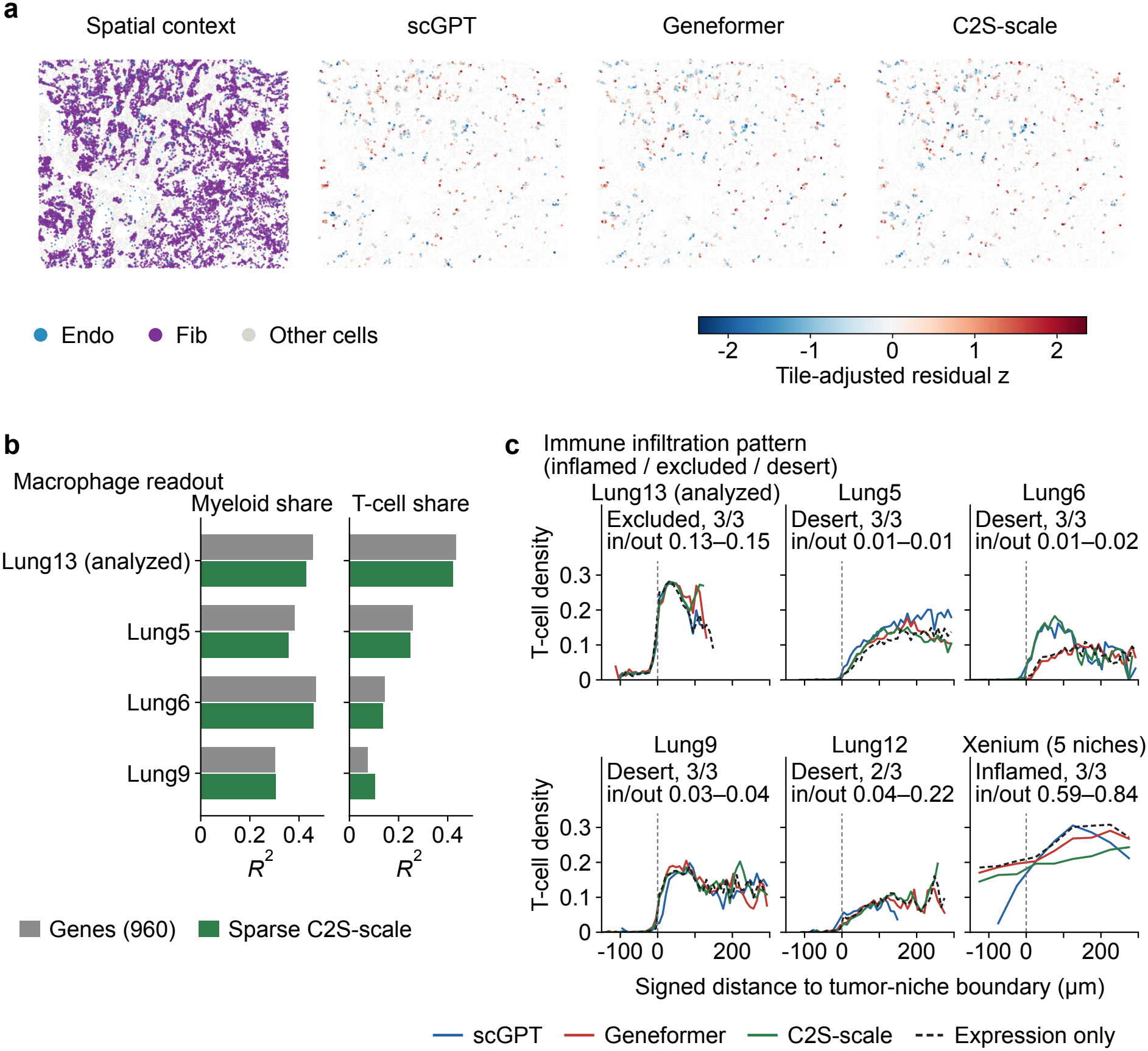
Contact-adjusted model fields, a macrophage neighborhood readout, and tumor immune infiltration patterns. **a**, CosMx contact-adjusted residual map of the endothelial–fibroblast program (standardized residuals after subtracting the fitted tile-level contact expectation; Methods; edge counts in Supplementary Table S14); color limit ±2.36; positive values, program burden above the expectation from contact density alone. Endo, endothelial; Fib, fibroblast. **b**, Macrophage readout: ridge *R*^2^ (penalty by inner cross-validation) for the myeloid and T-cell share of each macrophage’s 75-µm neighborhood, predicted from its 960 panel genes (gray) or its sparse C2S-scale profile (green), in the analyzed CosMx section (Lung13) and the three further donors with ≥3,000 macrophages (Lung5_Rep1, Lung6, and Lung9_Rep1, shown as Lung5, Lung6, and Lung9; Supplementary Table S4). **c**, Immune infiltration pattern (inflamed / excluded / desert): T-cell density (share of cells that are T cells, in 50-unit bins: 9 µm in CosMx, 50 µm in Xenium) against signed distance to the tumor-niche boundary (µm; negative inside; bins with *<*50 cells not drawn), per section, for each model’s niches (scGPT, Geneformer, C2S-scale) and for expression-only niches (dashed; post hoc). Text, the pattern from the inside/outside T-density ratio (*>*0.5 inflamed, 0.1–0.5 excluded, *<*0.1 or outside density *<*1% desert), the number of models agreeing, and the models’ ratio range; expression-only niches gave the majority pattern in every section. Tumor niche: CosMx, the niche with the highest tumor-cell share; Xenium (five niches, marker-based labels), the niche with the highest tumor-like epithelial share (the tumor/myeloid shared niche in all three models). Descriptive.

**Figure 8.**
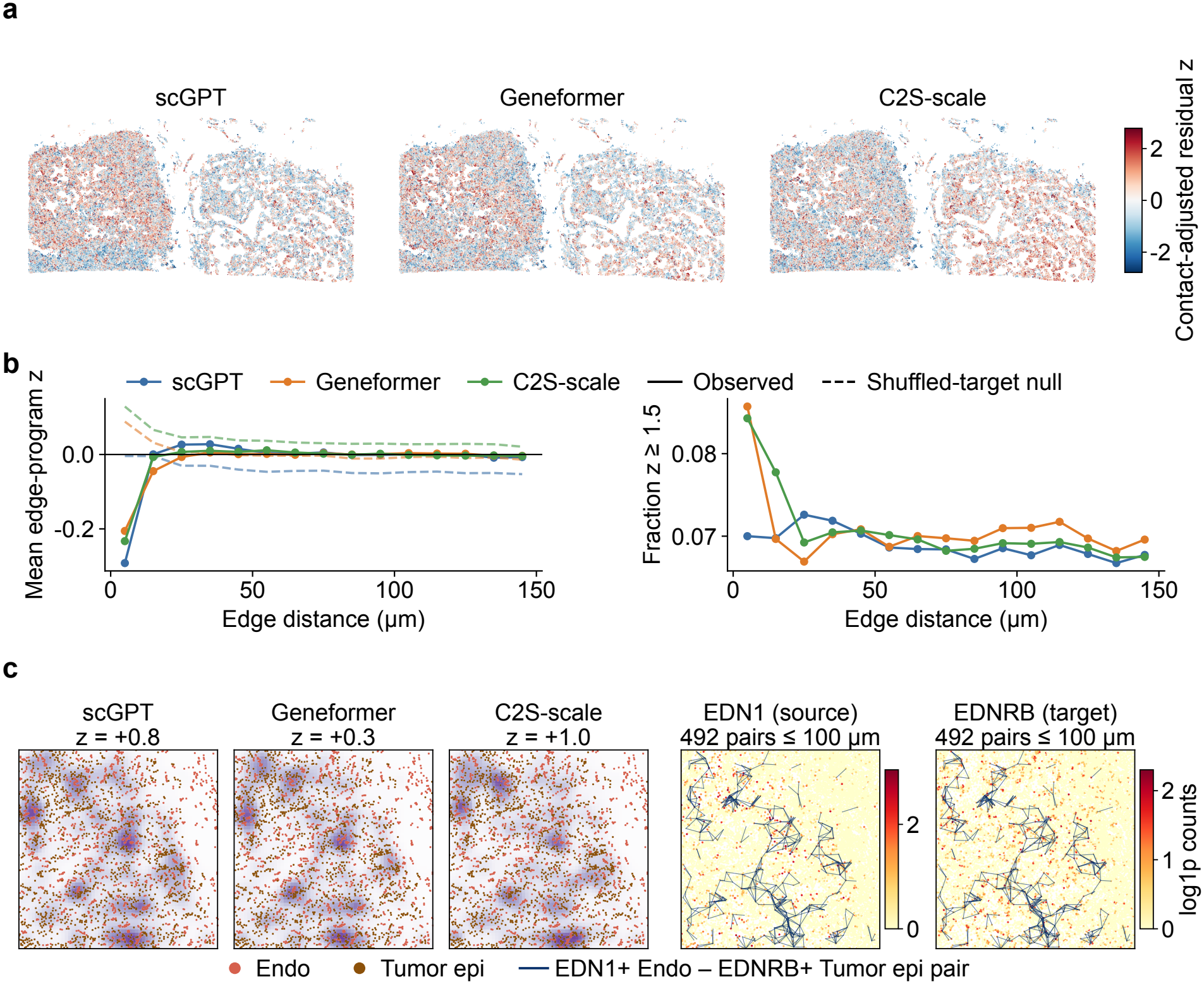
Xenium renal cell carcinoma (RCC) endothelial–epithelial case study with marker-based labels. **a**, Contact-adjusted residual maps of the count-adjusted program score on endothelial–tumor-like epithelial edges within 120 µm for the three models (Methods; edge counts in Supplementary Table S14). **b**, Mean edge-program score by cell–cell distance (µm) for the same edges against a coordinate-shuffled target-cell null (right, share of edges with *z* ≥1.5, observed only); bins with fewer than 50 edges omitted. **c**, EDN1/EDNRB crop: positive edge-load fields for the three models (titles, niche-free calibrated z for the pair; none passes *z*^*∗*^) and EDN1 (endothelial source) and EDNRB (epithelial target) expression, with all 492 EDN1-positive endothelial–EDNRB-positive epithelial pairs within 100 µm drawn as links. Gene-pair proximity tests for this crop in Supplementary Figure S13c. Abbreviations as in Fig. 4.

In Xenium, the endothelial–tumor-like epithelial EDN1/EDNRB crop, selected for display before the proximity tests, showed modest co-localization at 100 µm: EDN1-positive endothelial–EDNRB-positive epithelial pairs were 1.19-fold more frequent than under a target-position null (*P* = 0.001) and 1.18-fold than under a within-type null (*P* = 0.011), with similar folds on the transferred labels (Fig. 8c; Supplementary Figure S13c and Table S16). At 50 µm the excess was not significant (fold 1.01 and 1.14; *P* ≥0.12), and section-wide the panel screen gave fold 1.01, so the co-localization holds within the examined region only. The pair was not called in any model (niche-free *z* 0.31–0.99; *z*^*∗*^ 2.2–2.3) and remains an expression-supported candidate without convergent positive model-field evidence.

### Sparse cell profiles and model niches read out macrophage neighborhoods and tumor immune infiltration about as well as the measured expression

The interface statistic looked for program load specific to contacts; a weaker test is whether a cell’s profile reflects the composition of its neighborhood. Finally, we asked whether a cell’s sparse profile carries such information, compared with its measured expression. An exploratory readout showed that a CosMx macrophage’s own expression predicted the T-cell share of its 75-µm neighborhood (ridge regression; *R*^2^ 0.43; myeloid share 0.46; Fig. 7b; Supplementary Table S4). Part of the T-cell signal is consistent with contact or spillover: CD3 immunofluo-rescence inside macrophages correlated with the local T-cell share (*ρ* = 0.40), and its removal lowered the intrinsic-gene *R*^2^ to 0.29 (Supplementary Table S17). Sparse C2S-scale cell profiles predicted the T-cell share about as well as the 960 genes (*R*^2^ 0.42 vs 0.43) and the myeloid share slightly less well (0.43 vs 0.46), adding them to the genes gave no gain, and the 170 concept-labeled spatial features alone reached 0.27 (Supplementary Table S18): the sparse profile carries about as much local-neighborhood information as the full panel. In the three further donors with at least 3,000 macrophages, the myeloid share stayed predictable from genes and sparse profiles (*R*^2^ 0.30–0.47 and 0.31–0.46; scGPT and Geneformer profiles 0.25–0.43), whereas the T-cell share was predicted less well (genes, *R*^2^ 0.07–0.26; Fig. 7b; Supplementary Table S4). The macrophage features that varied most with distance to the tumor carried keratin and cornified-envelope labels and tracked the macrophage’s own PanCK fluorescence, a further sign of tumor contact or spillover (Supplementary Note S2).

The models’ tumor niches also gave a consistent reading of tumor immune infiltration, measured as the ratio of T-cell density inside to outside the tumor niche (Methods): 0.13–0.15 in the analyzed CosMx section, 0.005–0.038 in three further CosMx donors, where T cells were nearly absent from the tumor niche, 0.045–0.22 in Lung12, and 0.59–0.84 for the tumor/myeloid shared niche in Xenium (Fig. 7c; Supplementary Table S4g). Expression-only niches gave the same pattern in every section (post hoc); the agreement is partly built in, because tumor niches follow tumor-cell density. A tumor cell’s own expression predicted the local T-cell share only weakly (*R*^2^ 0.14 in this section, 0.14–0.24 in the donors; Supplementary Note S3).

## Discussion

interpreTFM evaluates what transcriptomic foundation-model hidden states encode on targeted spatial panels, separating the detection of individual genes from the generalization of concepts, and turns the hidden states into named features for cell- and tissue-level analysis. In both datasets, sparse features marked more concepts than raw hidden dimensions or an untrained SAE when tested on new observations of known genes, but scored below raw hidden dimensions on genes held out during feature selection, so they acted as gene detectors that carry concept labels; at CosMx Layer 2, sparse features of randomly re-initialized scGPT and Geneformer marked as many concepts or more. Steering results were consistent with this and with language-model benchmarks in which steering with SAE features did not outperform simple baselines.^42^ Concept information was nonetheless present and learned: a linear readout of the hidden states recovered the concept membership of unseen genes above co-expression and fell to chance in randomly re-initialized models (gene symbols alone gave a similar readout, so part of this information reflects gene families), and the sparse decomposition partly preserved it; no single feature carried it. Recent SAE studies have annotated single-cell foundation-model features with biological concepts;^34, 37, 38^ a whole-gene held-out split, an untrained SAE and a randomly re-initialized model test whether such annotations generalize beyond the genes a feature detects and depend on model training. Because features of re-initialized models carried as many labels here, a concept label assigned without these tests does not by itself show that the model learned the concept. The criteria also favored different depths, so a layer should be chosen for the criterion of interest.

At the cell and tissue level, the features summarized the measured biology in a named and reproducible form. Averaged per cell, they roughly doubled the population specificity of raw hidden dimensions and approached that of single marker genes, and in CosMx the leading features recovered marker programs more completely than the best single genes; these cell-level results held in four further donors. In the analyzed CosMx section, niches built from sparse features agreed with author cell types better than label-composition and expression-only niches, converged across models, and at finer resolution were more reproducible across seeds than expression-only niches; across patients their composition did not recur, but the concept labels that defined the tumor and stromal-lymphoid niches did. In the analyzed section and three further donors, macrophage sparse profiles predicted the myeloid composition of the neighborhood about as well as the full panel, and the models’ tumor niches gave consistent immune infiltration patterns, as did expression-only niches (an agreement partly built in, because tumor niches follow tumor-cell density). In Xenium the same territories, and in every section the same immune infiltration pattern, were recovered from expression alone, and no neighborhood readout exceeded the full gene panel. The representations are computed from the measured genes, so they cannot carry more information about a cell or its neighborhood than the panel itself; where the question concerned relationships between genes, the representations added to co-expression, and where it concerned a cell’s measured state, they matched expression while making it readable as named, reproducible features. The models’ cell embeddings likewise separated cell types at or below the log-normalized expression, in line with independent evaluations of frozen single-cell foundation models.^24, 25^

Spatial interface analysis needed calibration and quality control before interpretation. A conventional tile null produced 6–12% false-positive interface calls on contact-free fields, against about 5% for the calibrated statistic. Under calibration, CosMx calls fell to the expected rate; Xenium calls, although reproducible across models, tracked partner-marker transcripts at the contact, the signature of segmentation spillover,^40, 41^ and the tumor-interface enrichments disappeared when marker-detecting features were removed. Simple marker checks caught two further data artifacts: labels transferred from a normal-kidney reference assigned tubule identities to cells, mostly immune, that lacked their markers, and macrophage features that varied with tumor distance carried tumor keratin signal. Model-free co-localization placed in model niches was more stable: adhesion and collagen–integrin co-localization recurred directionally across CosMx donors, and CCL19–CCR7 localized to the Xenium lymphoid niche. Program-level interface statistics need a calibrated null and partner-marker checks, and a call marks a candidate context; this study did not test signaling direction, receptor activation or causality.

Each platform was analyzed in one section; a check in four further CosMx donors reproduced the cell-level comparisons, the macrophage read-out and the niches’ concept-level definition, but not niche composition, so tissue-level conclusions concern within-section organization. Interpretation depends on targeted-panel coverage, annotations and the fixed representative layers. The whole-gene split removes gene overlap between evaluation partitions, but cells are shared between them. The calibrated null controls per-pair false positives near 5% but not the false-discovery rate across pairs, and its power is limited: a planted contact effect of 5% of the edge program load was detected in 38% of tests and one of 1% in none, so the absence of CosMx calls rules out only contact effects of several percent. Xenium annotation relied on marker-based clustering of a 399-gene panel without pathology review. The foundation-model weights remained frozen, whereas the SAEs and niche models were fitted on the study sections, so the workflow is not a zero-shot evaluation. Independent sections, cell-disjoint evaluation, pathology-reviewed annotations and functional assays would address these limitations.

## Conclusions

**interpreTFM** provides a framework for evaluating sparse features from transcriptomic foundation models and relating them to spatial tissue organization. In the sections examined, the features detected individual genes and inherited their concept labels, and concept information about unseen genes remained linearly recoverable from the hidden states as a whole. At the cell and niche level, the features approached marker-gene specificity and recovered tissue territories that agreed across models; across CosMx donors, the niches’ defining concepts recurred but their composition did not, and no neighborhood readout exceeded the full gene panel. Calibrated interface calls of the models were near the chance rate in CosMx and followed partner-marker transcripts (segmentation spillover) in Xenium, whereas model-free co-localization of all but one ligand–receptor hit recurred directionally across CosMx donors. These results support layer-resolved evaluation and hypothesis generation; mechanistic implications remain to be established.

## Methods

### Study design and spatial transcriptomic datasets

We evaluated whether hidden representations from three pretrained single-cell foundation models could be converted into sparse, biologically annotated features and used for spatial tissue analysis. All analyses were performed independently for each spatial dataset, with no pooling of cells or model-derived representations across datasets. The workflow was applied to two imaging-based spatial transcriptomic datasets: a CosMx Spatial Molecular Imager lungcancer section containing 81,236 cells and 960 measured genes,^43, 44^ and a Xenium renal cell carcinoma (RCC) section containing 465,220 cells and 399 genes after model-panel compatibility filtering. One section from each of four further donors of the same CosMx study was used to test whether the CosMx results recur across patients (see Cross-donor replication). Both datasets contained two-dimensional cell coordinates and single-cell expression matrices. Coordinates are pixels of 0.18 µm in CosMx and micrometers in Xenium; radii and block sizes are given in these coordinate units, with the CosMx equivalent in micrometers.

The CosMx dataset included author-provided cell annotations in `author_cell_type`, spanning tumor, stromal, endothelial, myeloid, and lymphoid categories. The Xenium dataset has no tumor-aware cell annotation. Kidney-lineage labels were first transferred from the GSE183273 single-cell kidney reference (`subclass.l1; TAL1` and `TAL2` merged into `TAL`) with TACCO optimal transport on raw counts of the shared genes, assigning each cell the reference class with the highest score. Genes absent from the required model-gene union and all-zero cells were excluded before activation extraction. Because the transferred tubule-subtype labels were contradicted by their own panel markers (Results), Xenium cells were annotated from the panel for all primary analyses. Cells with fewer than 20 transcripts or fewer than 10 detected genes were flagged as low quality (40,395 of 465,220 cells, 8.7%) and not forced into a type. The remaining cells were median-library-size normalized, log1p-transformed, scaled (maximum 10), reduced to 30 principal components, connected in a 15-nearest-neighbor graph, and clustered by Leiden at resolution 1.0 (seed 0; 20 clusters). Each cluster was scored against curated marker sets restricted to panel genes (set score = mean z-scored cluster expression over the set’s genes) and took the top set’s label if that score was at least 0.75 SD and at least 0.25 SD above the runner-up; all 20 clusters passed, giving 13 types (Supplementary Figure S2). No cluster scored as normal proximal or distal tubule; the two epithelial clusters are labeled tumor-like epithelial, a name that does not establish malignancy. The transferred labels were retained as a secondary label set, and their marker support is reported in Supplementary Table S7. Neither the CosMx cell-type annotations nor either Xenium label set was used to train the foundation models, sparse autoencoders, or feature-concept associations; these labels were introduced only for downstream calibration, spatial summarization, and biological interpretation.

For downstream spatial summaries, fine annotations were additionally collapsed into broad cell-group categories. CosMx groups preserved major tumor, immune, endothelial, fibroblast, and plasma/B-cell compartments. Xenium marker-based types were used at their own resolution for the interface analyses. These groups were used for cell-composition and cell–cell-interface summaries and did not define the model-derived spatial niches.

### Foundation models and layer-resolved activation extraction

We analyzed layer-resolved hidden representations from three pretrained single-cell foundation models: scGPT,^20^ Geneformer,^19^ and C2S-scale,^23^ using the scGPT whole-human checkpoint, Geneformer V2-316M (18 layers; input of up to 4,096 genes), and C2S-Scale-Gemma-2-2B (26 layers). All pretrained model parameters were held fixed during activation extraction, and no model weights were fine-tuned on either spatial dataset. We evaluated 29 model-layer combinations: all 12 scGPT transformer blocks (`layer_0.norm2 through layer_11.norm2`), eight Geneformer layers (2, 5, 8, 11, 13, 15, 17, and 18), and nine C2S-scale layers (1, 7, 14, 16, 18, 20, 22, 24, and 26). Layer numbers reported in the text and figures are one-based, whereas stored layer identifiers retain the native zero-based indexing used during activation extraction; for example, `layer_1.norm2` corresponds to display Layer 2. Hidden-state dimensions were 512 for scGPT, 1,152 for Geneformer, and 2,304 for C2S-scale.

Inputs were constructed according to each model’s native tokenization and input representation. For scGPT, normalized expression values for nonzero genes were represented as gene tokens with a leading classification token, and activations were extracted from the `norm2` output of each evaluated transformer block. Geneformer received expression-ranked gene tokens and activations were extracted from the corresponding transformer hidden states at the evaluated layers. C2S-scale received its native gene-ranked cell representation, and hidden states corresponding to gene tokens were retained. Activations and row-level metadata were stored in a common sharded format.

Cell-level and gene-token-level representations were kept separately. Cell-level activation matrices were constructed from the classification-token representation for scGPT and Geneformer and from the mean hidden representation across gene-token rows for each cell for C2S-scale. Gene-token activation rows were retained separately for feature-concept association and subsequent projection of concept-associated features to cells. Special tokens were excluded from gene-token-level analyses and projection (`60694` and `60695` for scGPT; `<cls>` and `<eos>` for Geneformer). As a check of the inputs, each model’s cell-level activation matrix at the analyzed layer and the log-normalized expression (log1p CP10k) of the same cells were standardized, reduced to 50 principal components and classified by k-nearest neighbors (k = 15, cosine distance, five-fold stratified cross-validation) against the author labels (CosMx) and the marker-based labels (Xenium; 60,000 cells sampled with stratification), for populations of at least 30 cells; balanced accuracy is reported.

### Transcriptomic interpretability score

The transcriptomic interpretability score (TIS) follows the Machine Interpretability Score (MIS),^45^ which scores a unit by whether a similarity model, in place of human raters, can tell its high-activating from its low-activating examples; TIS uses measured expression as the similarity space. For each activation dimension *d* of a model layer, cells in the upper and lower activation quartiles formed high- and low-activation pools. In each of *T* = 800 trials, a query cell *q*_*t*_ was drawn from all analyzed cells, *m* = 32 reference cells were drawn without replacement from each pool after excluding the query, and the query was assigned to the pool whose mean expression profile 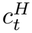or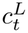 it resembled more:

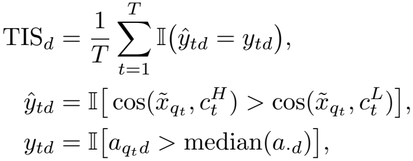

where *a*_*q t d*_ is the query’s activation along *d* and 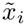is the expression profile of cell *i*, library-size normalized to 10^4^ counts, log-transformed, and L2-normalized, so that similarity is computed on measured expression. A high TIS indicates that the dimension’s variation is reflected in the measured transcriptome; it does not assign a concept or pathway to the dimension.

To estimate the contribution of the model representation beyond chance activation structure, each activation dimension was independently permuted across cells and the complete TIS procedure was repeated. Layer-level TIS gain was defined as the mean TIS across activation dimensions minus the corresponding mean obtained after permutation. The same activations rotated to their principal components (a dimension-matched change of basis) were also evaluated as a descriptive control.

### Sparse autoencoders and feature evaluation

To decompose dense foundation-model activations into sparse latent representations, a separate sparse autoencoder (SAE) was trained for every model-layer combination following sparse-dictionary-learning formulations.^28, 31, 33^ Each SAE operated on the hidden-state activations extracted from a single model layer and was trained independently of the SAEs for other layers and models, without labels, on all gene-token rows of its section; the held-out splits defined below apply to feature selection and evaluation. The latent dictionary size was set to eight times the corresponding hidden-state dimension, yielding 4,096 features for scGPT, 9,216 features for Geneformer, and 18,432 features for C2S-scale.

For an activation vector *x*_*i*_ ∈ ℝ^*h*^, where *h* is the hidden-state dimension of the corresponding foundation model, the SAE produced nonnegative latent features *z*_*i*_ and a reconstruction 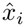,

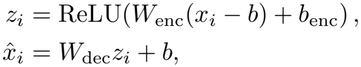

where *b*_enc_ is an encoder bias and *b* is a bias shared by the input and the output. Training minimized the mean squared reconstruction error together with an *l*_1_ penalty on latent activations to encourage sparse feature usage:

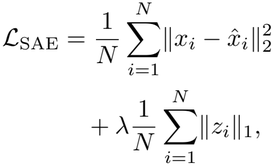

where N denotes the number of activation vectors used for training and *λ* controls the strength of the sparsity penalty. Activations were rescaled to mean norm 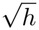 for training, and the fitted parameters were mapped back to the original activation scale. Decoder columns were constrained to unit Euclidean norm,

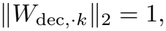

to remove the scale degeneracy between latent activations and decoder vectors. SAEs were optimized using constrained Adam with warm-up and dead-feature re-sampling, and the checkpoint with the lowest training objective was retained. The *l*_1_ weight was chosen from *λ*∈ {0.1, 0.3, 1, 3, 10} by a fixed rule: among weights with at most 2% of features active per token and a dead-feature rate of at most 0.25, the one with the lowest fraction of variance unexplained. Applied at the six representative layers, the rule selected *λ* = 1 at each of them (0.5–1.7% of features active per token, fraction of variance unexplained 0.06–0.11, no dead features), and *λ* = 1 was used at all layers. For comparison, an SAE of the same width with random, untrained weights was evaluated in the same way.

#### Concept universe and held-out feature-concept association

Biological concepts were generated separately for each measured gene panel using g:Profiler^46^ with human as the organism, false-discovery-rate correction at 0.05, and Gene Ontology biological process, molecular function, and cellular component terms together with Reactome and KEGG pathways. Terms were required to overlap at least two measured genes. For each dataset, these annotations were represented as a binary gene-by-concept matrix, in which each measured gene was labeled as a member or non-member of each retained biological concept. Because the concept universe was restricted to genes measured on each targeted panel, the analysis tests alignment with biological concepts observable on that panel and should not be interpreted as genome-wide pathway discovery.

Feature-concept association was evaluated on gene-token rows, each the context-dependent representation of one measured gene in one cell. For a given feature *k*, concept *c*, and activation threshold *τ* , a gene-token row was predicted to be concept-positive when its feature activation exceeded the threshold,

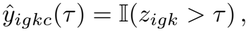

where *z*_*igk*_ denotes the activation of SAE feature *k* for gene *g* in cell *i*. The corresponding reference label was

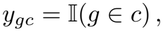

where *y*_*gc*_ indicates whether gene *g* is annotated as a member of concept *c*. Occurrences of the same gene can differ in activation but share the gene-level label, so alignment measures association with annotated gene-set membership, not pathway activity in a particular cell.

SAE features were evaluated using a held-out thresholded concept-alignment design^47^ with two held-out units. In the row split, gene-token rows were assigned deterministically to 70% training, 20% validation, and 10% test partitions using a hash of the seed, shard, row, and token identity, so rows corresponding to the same gene or originating from the same cell could occur in different partitions; this split assesses generalization to held-out observations of known genes. In the whole-gene split, measured genes were assigned to 70%, 20%, and 10% partitions and all rows of a gene followed it, so test genes were never seen during feature selection; cells remain shared between partitions. The Xenium Geneformer activation store recorded every cell twice at seven layers; duplicate rows were removed before evaluation.

Thresholds *τ* ∈ {0, 0.15, 0.30, 0.60} were applied to the latent activation scale (not as probabilities); predictions were compared with gene-level membership by F1, and associations were selected on the validation partition and evaluated on the test partition.

For comparison, the same concept-alignment analysis was applied to individual raw hidden-state dimensions, each split into its positive and negative parts so that all units are nonnegative, and to the un-trained SAE. All representations used the same rows, partitions, labels, thresholds, and metric. Concept alignment was summarized per concept, at threshold 0.15: for each concept with 5–100 panel genes and with positive rows in both the validation and test partitions, the unit with the highest validation F1 was selected (ties broken by the lowest index) and scored by its test F1. Concept F1 is the mean over concepts. Each concept counts once, so a broadly active unit cannot dominate the summary.

Randomly re-initialized models. To test whether concept alignment depends on model training, scGPT and Geneformer were re-initialized either completely or except for the gene-embedding table (each module’s own initialization, seed 0); activations at CosMx Layer 2 were extracted for a subsample of 30,356 and 40,480 cells, the same cells as for the trained models; and sparse SAEs were trained, selected and scored as above (the selection rule chose *l*_1_ weight 1, and 3 for the two re-initialized scGPT forms).

#### Gene-level readout of concept membership

To test whether concept membership of unseen genes could be decoded from a representation at all, each measured gene was represented by its mean hidden state (or mean SAE activation) over all of its tokens. Genes were divided into five whole-gene splits (10% test genes each). For each concept with 5–100 panel genes, an *l*_2_-regularized logistic regression (inverse regularization strength 1, balanced class weights) was fitted on the remaining genes and scored by test-gene AUROC. Inputs were standardized using the training genes; because standardization inflates the rare activations of sparse SAE features, a center-only version is reported as a post hoc sensitivity analysis. The best single dimension, chosen by training-gene AUROC, was scored in the same way. Controls were random Gaussian vectors of matched dimension, each gene’s co-expression profile in the same section (correlations with all panel genes, or their first 50 principal components), its mean expression in each cell type, and its mean expression and detection rate. Post hoc controls used the same splits: the mean hidden states of the reinitialized scGPT and Geneformer models described above; each gene symbol’s binary profile of character trigrams, and its first 50 principal components; and each representation after removing the part linearly predictable from these 50 components (least squares fitted on the training genes).

#### Population specificity of cell-level features

Cell-level profiles were the mean activation of each SAE feature, or of each raw dimension split into its positive and negative parts, over a cell’s gene tokens; single genes (log-normalized expression) were scored in the same way. Following scPISCES, the Gini specificity index (GSI) of a feature is the Gini coefficient of its mean activation across cell populations with at least 30 cells; GSI excess subtracts the mean GSI of the same feature over five label shuffles and is averaged over active features. For the marker-gene hit rate (MHR@100), cells were split in half within each population: in the first half, markers were the 100 genes with the largest mean expression difference between a population and all other cells, and the three features most correlated with population membership were selected; in the second half, MHR is the share of the 50 genes most correlated with each selected feature that are markers, averaged over populations. All representations were scored on the same cells: 75,745– 81,067 CosMx and 63,068–84,895 Xenium cells read from evenly spaced activation shards, excluding cells split across shard boundaries.

#### Causal steering check

To test whether a concept-labeled feature carries information about its concept beyond the genes it detects, we added its decoder direction to the hidden states of the representative CosMx layers in held-out cells and compared the resulting change in the model’s output between the concept’s other member genes, on which the feature does not fire, and non-member panel genes (AUROC). The same intervention with a random feature of similar firing frequency, a random direction, and the identity direction of the feature’s top detected gene, all at the same norm, served as controls; the full protocol is given in the Supplementary Methods and the results in Supplementary Table S2.

#### Cell-level projection and representative-layer selection

For downstream spatial analyses, sparse-SAE features were retained by a fixed rule: for each concept with 5–100 panel genes, the feature with the highest validation F1 at threshold 0.15 in the row split was kept if that F1 was at least 0.40, and features were then deduplicated. A layer required at least 10 retained features; layers below this are marked in Fig. 2, and the counts at the analysis layers are given in Supplementary Table S1. Selection used validation rows only.

Selected SAE features were projected from gene-token rows to cells by mean aggregation across a cell’s mapped gene tokens. The retained sets contained 194, 261, and 170 features for CosMx scGPT, Geneformer, and C2S-scale, respectively, and 118, 125, and 94 features for their Xenium counterparts (Xenium Gene-former after removal of duplicated token rows).

The representative layers were chosen in a preliminary screen by a composite of TIS gain and concept alignment: scGPT `layer_1.norm2`, Geneformer `layer_1`, and C2S-scale `layer_13` for CosMx, and scGPT `layer_2.norm2`, Geneformer `layer_1`, and C2S-scale `layer_13` for Xenium. They were kept fixed for all analyses reported here.

### Spatial niche discovery and niche-associated concept analysis

For each cell *i*, let *u*_*i*_ ∈ ℝ^*p*^ denote its selected interpretable-feature vector and *s*_*i*_ ∈ ℝ^2^ its spatial coordinate. For a radius *r*, the spatial neighborhood of cell *i* was

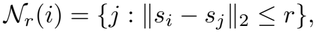

and the corresponding microenvironment vector was the uniform neighborhood mean

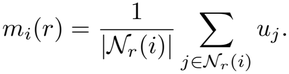

Candidate niches were clustered either from *m*_*i*_(*r*) alone (`m`) or from the concatenated cell-and-microenvironment representation [*u*_*i*_, *m*_*i*_(*r*)] (`xm`). Features were standardized, reduced to at most 50 principal components, and clustered by k-means or a full-covariance Gaussian mixture model. The screen covered radii of 100, 120, and 150 coordinate units (CosMx pixels of 0.18 µm, 18–27 µm; Xenium 100–150 µm), three to five clusters, and random seeds 0, 1, and 2. Configurations with a cluster smaller than 150 cells were rejected.

For configuration *c*, niche stability was measured by seed-to-seed ARI, microenvironment coherence by a spatial-neighborhood purity z-score against a tile-shuffle null, and niche separability by cosine distance between niche centroids. The ranking score was

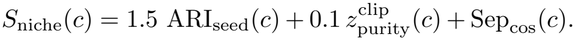

Neighborhood purity was compared with a spatial tile-shuffle null in which niche labels were permuted within tiles of width four times the neighborhood radius. Fragmentation was reported as a quality-control statistic but was not included in *S*_niche_.

For CosMx, the configuration with the highest *S*_niche_ was used for each model: scGPT radius 100, `xm`, k-means, *k* = 3; Geneformer radius 120, `m`, k-means, *k* = 3; and C2S-scale radius 100, `m`, k-means, *k* = 3. For Xenium we report five niches per model to resolve substructure within the three large territo-ries produced by the highest-scoring configurations (*k* = 3), using the configuration with the highest *S*_niche_ among *k* = 5 configurations: scGPT radius 100, `xm`, Gaussian mixture; Geneformer radius 120, `m`, k-means; and C2S-scale radius 100, `xm`, k-means. For all selected *k* = 5 configurations the purity z-score exceeded its clip, so their ranking was determined by seed-to-seed ARI and separability. Seed-to-seed ARI, purity, and its tile-shuffle null for these configurations, and the three-niche solutions, are given in Supplementary Table S5. As an expression-only base-line, the same sweep and ranking score were applied to 50 truncated singular-value-decomposition (SVD) components of log-normalized expression, matching the at most 50 principal components used for model features, (winners: radius 120, `m`, Gaussian mixture, *k* = 3 for CosMx; radius 100, `m`, Gaussian mixture, highest-scoring *k* = 5 configuration for Xenium), and expression-only niches were also computed at each model’s selected configuration.

Cell-type enrichment was summarized by standardized residuals from the niche-by-cell-type contingency table.

For niche-associated concept analysis, interpretable-feature values inside and outside each selected niche were compared by Wilcoxon rank-sum test, with Benjamini-Hochberg correction across tested features. Effect size was the difference in mean feature value, and the difference in active-cell fraction was also reported. Features were ranked by mean difference (ties broken by the difference in active-cell fraction). Broad biological theme labels were assigned from concept names for visualization and were not used as supervised targets.

As a label-composition baseline, Squidpy-like neighborhood composition vectors were constructed by averaging one-hot cell-type labels over spatial neighbors, reducing them to 15 principal components, and clustering by k-means with *k* equal to the number of model niches (3 for CosMx, 5 for Xenium) and 20 initializations. This baseline directly encodes cell-type composition and is therefore expected to favor agreement with the same cell-type labels. Partitions were compared between models by ARI, which does not depend on niche numbering; for niche-level comparisons, niches were matched across models by composition (Hungarian assignment on the maximum absolute label-fraction difference).

Concept-label concordance of matched niches used the top-25 positive-effect concept labels per niche, keyed by concept identifier; for each model pair, the mean Jaccard index over the composition-matched niches (three per model pair in CosMx, five in Xenium) was compared with the mean over the un-matched niche pairs (six and twenty). The transcript-depth check for Xenium computed the ARI between each model’s niches and a three-way partition of all cells by transcript-count tercile and compared it with the ARI against marker-based labels.

### Ligand–receptor co-localization and spatial crosstalk scoring

#### Ligand–receptor co-localization screen

The model-free screen (exploratory analysis C3; Supplementary Table S19) tested every CellChatDB human interaction whose ligand gene(s) and all receptor-subunit genes were on the panel (397 CosMx and 25 Xenium pairs). A gene was positive at 1 transcript and a complex when every subunit was positive; pairs with fewer than 100 positive cells on either side were not tested. The statistic was the number of ligand-positive–receptor-positive pairs of distinct cells within 100 coordinate units (about 18 µm in CosMx, contact range; 100 µm in Xenium, neighborhood range); the null permuted positivity within cell type (CosMx author labels; Xenium marker-based labels, with low-quality cells as a type), independently for ligand and receptor (100 permutations). Fold = (obs + 1)*/*(null mean + 1); two-sided normal P values were Benjamini–Hochberg adjusted per dataset. A hit required *q <* 0.05 and fold ≥1.2; a robust hit was a hit at both ≥1 and ≥3 transcripts. For each robust hit, the share of its close pairs inside each niche was divided by the niche’s share of cells. The screen was repeated at matched neighbor counts, at the radius where the mean number of other cells within reach equals that of the other dataset at 100 units (Xenium 15.4 µm, about 7 cells; CosMx 652.8 pixels, 117.5 µm, about 287 cells), with the four donors at the CosMx matched radius; the statistic, null, and hit rules were unchanged.

CellChat provided an external, expression-based set of candidate ligand–receptor (LR) pairs. It was run within each model’s selected niches on 3,000 sampled cells per group (seed 17). Matched-niche pathway profiles were the summed communication probability per pathway, normalized to fractions, and compared across models by Spearman correlation for composition-matched versus unmatched niches. CellChat rows of a cell pair in a niche were crosstabulated with that pair’s calibrated call; because the program score averages all retained features, every row of the pair receives the pair’s call, and concept-label support of the rows is reported as a descriptor (Supplementary Tables S10 and S19). All LR pairs come from CellChatDB.

#### Interface program scoring

Cell-level interpretable program load was computed from the selected feature matrix *U* ∈ ℝ^*n×p*^. For each feature, values were clipped at the 99.5th percentile, standard-ized, and restricted to the positive component:

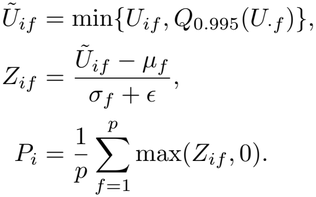

Here *P*_*i*_ summarizes nonnegative model-derived pro-gram burden for cell *i*. Because *P*_*i*_ can track transcript count within cell types, it was residualized within each exact cell type on log transcript count, log detected genes, and log cell area (CosMx: bounding-box width × height), and the residual 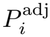 was used for testing. Undirected spatial edges were constructed within a fixed radius *r*,

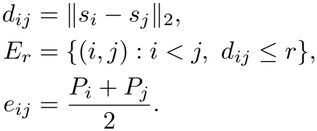

with *r* = 120 coordinate units in both datasets (about 22 µm in CosMx and 120 µm in Xenium). Eligible cross-group edges had both endpoints in the same niche and were grouped by unordered broad cell-group pair *g* and niche *n*. For supported edge groups,

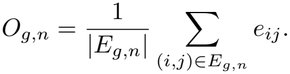

The local spatial null used *B* = 100 permutations of the adjusted program score within strata defined jointly by fixed 120-unit spatial blocks (about 22 µm in CosMx; 120 µm in Xenium), niche identity, and exact cell-type annotation, an exact within-stratum permutation test. Tissue coordinates, graph topology, and annotations remained fixed. For permutation *b* = 1, … , *B*, let 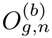 be the null score. The strict crosstalk statistic was

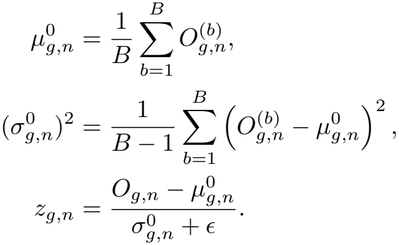

The empirical upper-tail value was

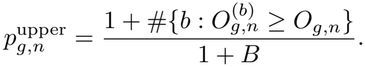

Edge scores and *O*_*g,n*_ were computed from *P* ^adj^. Positive *z*_*g,n*_ indicates program enrichment on a cell-group interface relative to the spatial null; negative *z*_*g,n*_ indicates depletion relative to the same null, not biological avoidance.

The call threshold was calibrated on contact-free negative controls. Smooth random fields (Gaussian-bump fields at length scales of 60, 150, and 400 units mixed with independent noise) were generated with their neighbor correlation matched to that of the adjusted score and passed through the same null (two fields per length scale, 50 permutations). The threshold *z*^*∗*^ was the 95th percentile of |*z*| over these fields, pooled over pairs and floored at 2, and a pair was called enriched if *z*_*g,n*_ *≥ z*^*∗*^ and depleted if *z*_*g,n*_ ≤ −*z*^*∗*^. This controls the per-pair false-positive rate near 5% under autocorrelated contact-free fields but is not a false-discovery-rate control across pairs. Power was estimated in CosMx (scGPT, niche-free statistic) by adding a contact effect to the adjusted score of the cells of one type that touched a cell of the partner type, sized to raise the pair’s edge-mean statistic by 0.25–5% of its mean edge program load, for 12 cell-type pairs spanning the range of edge counts (100 permutations, two permutation seeds). The same fields were passed through a conventional, uncalibrated 400-unit tile null (72 µm in CosMx; 400 µm in Xenium) with 20 permutations to estimate its false-positive rate (Supplementary Table S9). The effect size was 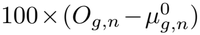 divided by the mean unadjusted edge program load of the same edges. A niche-free version of the statistic (pairs and strata without niche) was computed for each model to compare calls across models, because niche numbers cannot be aligned between models; case-study crops were evaluated with this niche-free statistic. For CosMx, a partner-marker check defined the top-10 panel genes of each author cell type (one-vs-rest Wilcoxon on log1p CP10k) and, for each cell pair called in at least one model, computed the niche-free z within terciles of the contacts’ partner-marker transcript counts; the gradient across terciles was summarized by its median over pairs with a block bootstrap over pairs (2,000 resamples), following the Xenium analysis. For spillover-robust (marker-excluded) calls, every retained feature whose fired-on genes (genes whose tokens activate it, *z >* 0.15, in at least 20% of their occurrences, plus the smallest set of top genes covering 80% of its activation mass) included a top-10 marker of either cell type of a pair was removed from the program score, and the adjusted score, the niche-free block null and *z*^*∗*^ were recomputed; a pair was spillover-robust if |*z*|≥*z*^*∗*^ with the sign of its original call in at least two of three models (Supplementary Table S11).

### Contact-adjusted interface validation and expression-proximity overlays

For selected case-study interfaces, we examined program burden after accounting for local cell-pair abundance. These separately generated diagnostics used radii of 138 coordinate units for CosMx (about 25 µm) and 120 µm for Xenium, with contact-model tile widths of 500 units (90 µm) and 300 µm, respectively. Tile-level program summaries were regressed on local pair-edge abundance. For each edge midpoint, the fitted tile-level contact expectation was subtracted from its program load. The residuals were then centered and scaled by their empirical standard deviation for display. The residual measures departure from the fitted contact expectation and does not adjust for other spatial confounders.

We also summarized edge-program scores as a function of cell–cell distance and compared the observed curves with a coordinate-shuffled target-cell null. Distance bins with fewer than 50 edges were dropped. Departures from this null would indicate organization beyond contact geometry.

For case-study crops, model interaction fields were overlaid with same-section expression/proximity support for nominated LR systems or marker-pair contexts. Links were drawn between nearby source and target cells when the relevant lig- and and receptor or receptor-complex component were detected. The three model-field columns aggregate all term-supported candidate LR rows for the selected cell pair, whereas the expression columns focus on curated genes used to interpret each crop: AREG/EGFR, COL9A2/ITGA1, and JCHAIN/PECAM1 in CosMx; EDN1/EDNRB in Xenium. JCHAIN/PECAM1 is the marker-pair context, and ITGA1 represents one component of the nominated ITGA1–ITGB1 receptor complex. In the CosMx crops, the drawn links are a capped, score-ranked display subset and do not count qualifying cell pairs; in the Xenium crop, all ligand-positive source–receptor-positive target cell pairs within 100 µm are drawn. Gene-pair proximity in each crop was tested as the number of ligand-positive source-type and receptor-positive target-type cell pairs within 50, 100, and 150 coordinate units (CosMx 9, 18, and 27 µm) against two nulls with 1,000 permutations each: a target-position null that resamples target positions from all cells in the crop, and a composition-preserving within-type null that shuffles gene positivity among cells of the same type, independently for source and target. Model support of a crop’s cell pair was read from the niche-free calibrated statistic (*z > z*^*∗*^).

### Macrophage readout, cross-donor replication, and exploratory analyses

The exploratory macrophage readout (CosMx) was defined as follows. For each macrophage, the targets were the T-cell and myeloid shares of its 75-µm neigh-borhood; predictors were its own panel expression (all 960 genes; intrinsic genes only; with CD3 immunofluorescence regressed out as a sensitivity row) or its sparse C2S-scale cell profile (all latents active in at least 1% of macrophages; the 170 spatial features; genes plus latents). Ridge regression was evaluated by blocked five-fold cross-validation with 500-µm spatial blocks. Because the latents outnumber the macrophages, Supplementary Table S18 reports a sensitivity setting (penalty chosen per target from a grid scaled by the training size); for the latent rows the penalty sat at the grid edge, so those values are lower bounds. The genes-versus-sparse-profile comparison in Figure 7b and the cross-donor comparison choose the penalty by inner cross-validation.

#### Cross-donor replication

One section from each of four further donors of the CosMx non-small-cell lung cancer (NSCLC) study (Lung5_Rep1, Lung6,Lung9_Rep1, and Lung12; 71,304–98,002 cells; the same 960-gene panel) was processed with the pipeline of the analyzed section unchanged: activations were extracted with the same code at the three representative layers; the sparse autoencoders, selected feature sets, and cell projection were reused without retraining or reselection; niches were selected in each donor with the same sweep grid and score; and author cell types came from the same source. A niche of the analyzed section recurs in a donor if the co-sine between its Pearson-residual cell-type profile and that of the best-matching donor niche is at least 0.8 (22 author labels, and tumor subtypes merged as a second variant); the niche structure replicates if at least two of the three niches recur in every donor. Agreement between model and expression-only niches replicates if the median ARI over models in each donor is no more than 0.10 below that of the analyzed section. The cell-level pattern replicates if, in every donor, sparse features exceed raw hidden dimensions on GSI excess for all three models and reach at least 60% of the single-gene value. The macrophage readout replicates if the myeloid share is predicted with *R*^2^ ≥0.25 in every donor with at least 3,000 macrophages (ridge penalty chosen by inner cross-validation; Lung12, with 1,291 macrophages, is reported but excluded). Seed reproducibility at finer resolution was the median seed-to-seed ARI over the 24 four- and five-niche configurations of the sweep grid; it favors the models in a donor–model case if the model’s median exceeds that of expression-only niches, and replicates if it does so in at least 10 of the 12 cases. A niche recurs at concept level in a donor if its top-25 concept labels overlap those of some donor niche (maximum Jaccard) more than 25 concepts drawn at random from the union of that model’s niche top-25 lists (*P <* 0.05; 1,000 draws), and recurs across donors if it does so in at least three of four donors for at least two of three models. The ligand–receptor co-localization screen was rerun un-changed in each donor on author labels; a CosMx hit replicates if it is a hit at ≥1 transcript in at least three of four donors, and the share of its close pairs in each donor niche was reported descriptively. The recurrence test for expression-only and label-composition niches and the comparison of niche–cell-type agreement with the baselines were added post hoc.

#### Immune infiltration pattern

For each section (the analyzed CosMx section, the four donors, and Xenium at five niches), each model’s tumor niche (CosMx, the niche with the highest tumor-cell share; Xenium, the niche with the highest tumor-like epithelial share) defined a signed distance of every cell to the niche boundary (negative inside). T-cell density was the share of cells that are T cells (CosMx author T-cell types; Xenium marker-based T cells) inside versus outside the niche, and profiles were summarized in 50-unit bins of signed distance (9 µm in CosMx; 50 µm in Xenium). A section was classified as inflamed if the inside/outside ratio exceeded 0.5, excluded at 0.1–0.5, and a desert below 0.1 or when the outside T density was below 1% of cells; agreement of the classification across models is the reported quantity. Expression-only niches were classified with the same rule post hoc. Two tumor-intrinsic readouts of infiltration (the T-cell share around tumor cells predicted from their own expression without lymphocyte markers, and an IFN-*γ* versus hypoxia comparison) are reported in Supplementary Note S3.

#### Exploratory analyses

Eight further within-section exploratory analyses (S1–S5 and C1–C3) are reported in Supplementary Table S19, which also lists the marker-excluded, immune-infiltration, and crop-proximity analyses C4, I1, I2, and D1 (Supplementary Notes S1–S3). Each used a single section; no interface z-score or cross-donor data entered these analyses.

### Statistical analysis

Permutation tests and z-scores were defined within the corresponding niche, crosstalk, and contact-adjusted interface analyses. Multiple testing for niche-associated features was controlled by Benjamini-Hochberg adjustment. ARI and normalized mutual information were used as descriptive partition-agreement metrics. Spearman correlation was used for rank agreement. Randomized stages used the fixed seeds specified in the relevant Methods subsections, and seed-robustness summaries were treated as descriptive rather than specimen-level inferential statistics. Analysis rules (the *l*_1_-weight and feature-retention rules, the concept-F1 summary, the steering protocol, the replication criteria, the criteria of the exploratory analyses, and the further niche, co-localization, marker-excluded and immune-infiltration analyses) were fixed before the corresponding results were computed; analyses added afterwards are labeled post hoc.

Each platform is represented by one analyzed section, so cells and edges are technical or spatial units nested within a specimen, not independent biological replicates; the cross-donor check is reported separately (above).

## Supporting information

supplementary_clean

## Data availability

All spatial data analyzed in this study are public. The CosMx NSCLC formalin-fixed, paraffin-embedded (FFPE) data (NanoString CosMx Spatial Molecular Imager^44^) were obtained from the SpatialCorpus-110M collection of Nicheformer^43^ (Hugging Face dataset `theislab/SpatialCorpus-110M`, revision 5494a62f, file `nanostring_lung_annotated.h5ad`); the analyzed section is Lung13, and the four sections of the cross-donor check are Lung5_Rep1, Lung6, Lung9_Rep1, and Lung12 from the same file. The Xenium section is the 10x Genomics public dataset “Human Renal Cell Carcinoma (FFPE)” (`Xenium_V1_Human_Kidney_FFPE_ Protein_updated`; Xenium Human Multi-Tissue and Cancer Panel with a 28-gene add-on; Xenium Onboard Analysis 4.0). The kidney reference used for label transfer is available from GEO under accession GSE183273. Processed analysis objects are deposited at Zenodo (https://doi.org/10.5281/zenodo.24293514) under a CC BY 4.0 license: the cell-level matrices of the retained sparse features at the six analysis layers (the analyzed CosMx section, the four further CosMx sections, and the Xenium section), the niche assignments of the models and of the expression-only and label-composition baselines, the marker-based and transferred cell annotations of the Xenium section, and the source tables of the figures and Supplementary Tables. The objects are keyed by the cell identifiers of the public source files and contain no expression values, coordinates, or author annotations, which are obtained from the sources above.

## Code availability

Source code for **interpreTFM** is available at https://github.com/maiziezhoulab/interpreTFM, with documentation at https://interpretfm.readthedocs.io. Version 1.0 (archived at Zenodo, https://doi.org/10.5281/zenodo.24298820) provides the complete analysis pipeline, with configurations and tests that reproduce the reported results, a tutorial, and the trained sparse autoencoders of the six analysis layers (Hugging Face, https://huggingface.co/YunfeiHu/interpreTFM). Pre-trained foundation-model weights, activation stores, and processed data are not included in the repository.

## Competing interests

The authors declare that they have no competing interests.

## Funding

This work was supported by the NIGMS Maximizing Investigators’ Research Award (MIRA) R35 GM146960, NSF CAREER DBI-2541736, and Vanderbilt Seeding Success Grant FF_300627.

## Notes

### Competing Interest Statement

The authors have declared no competing interest.

### Summary of Updates

Text revision only. Data, analyses, results, figures and Supplementary files are unchanged. Abstract: rewritten as a structured abstract (Background, Results, Conclusions) of 250 words. It now states the central question, whether a concept-labeled SAE feature represents its concept or only the genes it detects, and the practical conclusion. Introduction: final paragraph shortened. Results: the section on sparse features is split into a gene-level section (concept detection on held-out genes) and a cell-level section (population specificity). Section titles are revised to state each finding, each section opens with the question it addresses, and the interface case examples are moved into the interface section. The immune infiltration paragraph is shortened; its detailed values remain in Supplementary Table S4. Discussion and Conclusions: revised to match the Results, including what the re-initialized model result implies for concept labels and how the cell embedding result compares with independent evaluations of frozen single-cell foundation models. Methods: the transcriptomic interpretability score description is condensed and now cites the Machine Interpretability Score it follows (Zimmermann et al., NeurIPS 2024); the crosstalk methods are reordered to follow the Results; donor details are moved to Study design. References: one reference added and bibliographic details corrected in two entries. Wording edited throughout for clarity and length.

