## supplementary_clean for "interpreTFM: an interpretable framework for decoding transcriptomic foundation models"

Hu et al.

#### Contents

**1 Supplementary Notes** **2**

**2 Supplementary Methods** **3**

**3 Supplementary Figures** **4**

**4 Supplementary Tables** **19**

**Supplementary References** **46**

### 1 Supplementary Notes

**Supplementary Note S1. Concept labels of the niche-defining CosMx features.** We compared the top-25 concept labels of the niche-defining features between models (Methods). In the composition-matched CosMx niches, scGPT–Geneformer and scGPT–C2S-scale each shared 2, 8, and 8 labels in the tumor, macrophage-rich, and fibroblast/CD4-T niches (Jaccard 0.042, 0.190, and 0.190). Geneformer–C2S-scale shared 6, 10, and 11 labels (0.136, 0.250, and 0.282). Unmatched niche pairs scored 0–0.087. Averaged per model pair, matched niches scored 0.14–0.22 and unmatched niche pairs 0.014–0.021. Most shared labels fit the lineage of their niche. The tumor niche has keratin filament. The macrophage-rich niche has MHC class II protein complex, macrophage activation involved in immune response, and chemokine activity. The stromal niche has fibrillar/FACIT collagen trimer, regulation of T cell tolerance induction, and dendritic cell antigen processing and presentation. Labels that are not tissue-specific also occur: forebrain neuron differentiation, lens fiber cell development, synapse pruning, and hepatocyte apoptotic process. These labels name panel-gene sets that the features detect.

In Xenium at five niches, the same comparison gave 0.308, 0.262, and 0.287 for matched niches and 0.073, 0.095, and 0.099 for unmatched niches. The three shared niches had Jaccard 0.28–0.47 (11–16 of 25 labels shared). The matched model-specific niches had 0.04–0.32 (2–12 shared). At three niches the values were 0.360, 0.328, and 0.383 for matched niches and 0.061, 0.073, and 0.084 for unmatched niches.

As an exploratory analysis (Supplementary Table S19, S4), we compared the labels with gene-level enrichment. For each CosMx niche, the concept labels of its 25 defining features were compared with the terms enriched among the niche's top-50 differentially expressed (DE) genes. The overlap was above chance in the macrophage-rich and stromal niches (4–28% of labels, $P \leq 0.035$ in 6 of 9 model-niche pairs) but not in the tumor niche. There the feature labels (keratin filament, cornified envelope, skin barrier) were supported by their genes but absent from the gene-level enrichment. Of the stromal-niche labels that the over-representation analysis (ORA) did not return, 67–85% were not supported by their own genes (Cohen's $d \leq 0.2$). In Xenium at three niches, the overlap was at chance in 8 of 9 model-niche pairs.

**Supplementary Note S2. Tumor-contact signal inside CosMx macrophages.** This analysis is exploratory (Supplementary Table S19, S2). Within macrophages, the features that varied most with distance to the tumor carried keratin and cornified-envelope labels. Their activation fell with distance ($\rho$ −0.23 to −0.25) and followed the macrophage's own PanCK fluorescence ($\rho$ 0.27–0.31). The signal reflects tumor contact or transcript spillover from neighboring cells [1, 2], as for the CD3 immunofluorescence in the macrophage neighborhood analysis and the Xenium interface calls. No feature followed the distance gradient more closely than the mean expression of its label's genes.

**Supplementary Note S3. Tumor-cell expression and local T-cell infiltration.** We ran two exploratory analyses, I1 and I2 (Supplementary Table S19).

*I1.* We asked whether a tumor cell's own expression predicts the T-cell share (T and NK cells) of its 75-µm neighborhood. Tumor cells were the author tumor types in CosMx and the tumor-like epithelial cells in Xenium. The predictors were the cell's panel genes without the top-10 markers of every T, NK, and B type. We used ridge regression with blocked five-fold cross-validation (500-µm blocks). $R^2$ was 0.138 in the analyzed section and 0.142, 0.155, 0.241, and 0.141 in Lung5, Lung6, Lung9, and Lung12. In Xenium it was 0.263. With CD3 immunofluorescence regressed out, $R^2$ was 0.131 in the analyzed section and 0.106–0.222 in the donors. A post hoc control residualized the T-cell share on the non-tumor share of the neighborhood. $R^2$ then fell to 0.006–0.066 in the donors but stayed at 0.160 in the analyzed section and 0.190 in Xenium. In the donors the prediction therefore mainly reflects a tumor cell's position at the tumor edge. We repeated the prediction with the sparse C2S-scale profile after removing the marker-detecting features, which left 15,520–15,976 of 18,432 features, on subsamples of 8,000 cells. It reached $R^2$ 0.094–0.252, similar to the genes on the same cells (0.109–0.254).

*I2.* Following I1, we scored each tumor cell for interferon-$\gamma$ response and for hypoxia. The interferon-$\gamma$ score used the panel genes of GO:0034341 (response to type II interferon). After removal of the top-10 markers of every T/NK, B/plasma, and myeloid type, 34–36 of 40 genes were kept. The hypoxia score used GO:0071456 (cellular response to hypoxia), with 30–31 of 31 genes kept. Each score was related to the T and NK share of the cell's 75-µm neighborhood by partial Spearman correlation, controlling for the neighborhood's non-tumor share and the cell's log transcript count. The 95% CIs came from 1,000 block-bootstrap resamples of 500-µm blocks. The confirmation set was three serial sections not used elsewhere in this study (Lung5_Rep2, Lung5_Rep3, and Lung9_Rep2; 16,555–96,822 tumor cells). Partial $\rho$ was +0.006, −0.008, and +0.040 for the interferon-$\gamma$ score and −0.011, −0.009, and −0.019 for the hypoxia score. In the analyzed section and the four donors the values were +0.010 to +0.052 and −0.083 to +0.003, and in Xenium −0.099 and −0.007. Without covariates, the hypoxia score fell with the T-cell share in every CosMx section ($\rho$ −0.048 to −0.158). With the covariates this association largely disappeared (−0.009 to −0.024, except −0.083 in the analyzed section), consistent with tumor-edge position. No retained concept-labeled feature carried either label, so model support could not be evaluated.

#### 2 Supplementary Methods

**Causal steering check (Supplementary Table S2).** The features tested were the concept-labeled sparse features with a test F1 $\geq$ 0.5 and at least one scorable other member gene. Fired-on genes are defined in the main-text Methods, and the concept's remaining panel genes are its other members. For each feature, a vector was added to the hidden states at the gene positions of held-out cells. In C2S-scale and Geneformer these were the first 25 genes of the cell's rank-ordered input, which were then excluded from scoring. In scGPT they were all input genes. The change in the model's output was compared between the concept's other member genes and non-member panel genes by the area under the receiver operating characteristic curve (AUROC). The output scored was the next-gene log-probability of every panel gene for C2S-scale, the expression predicted for every panel gene by the decoder that reads the cell embedding (MVC head) for scGPT, and the masked-gene log-probability for Geneformer. Three controls were steered at the same norm (scale $\alpha = 4$): a random feature with similar firing frequency, a random direction, and the gene-identity direction of the feature's top fired-on gene. A concept effect beyond the detected genes was defined as both random-control contrasts exceeding 0.05 with CIs excluding 0. A gene-identity contrast whose CI includes 0 means that steering the detected gene's identity direction has the same effect.

#### 3 Supplementary Figures

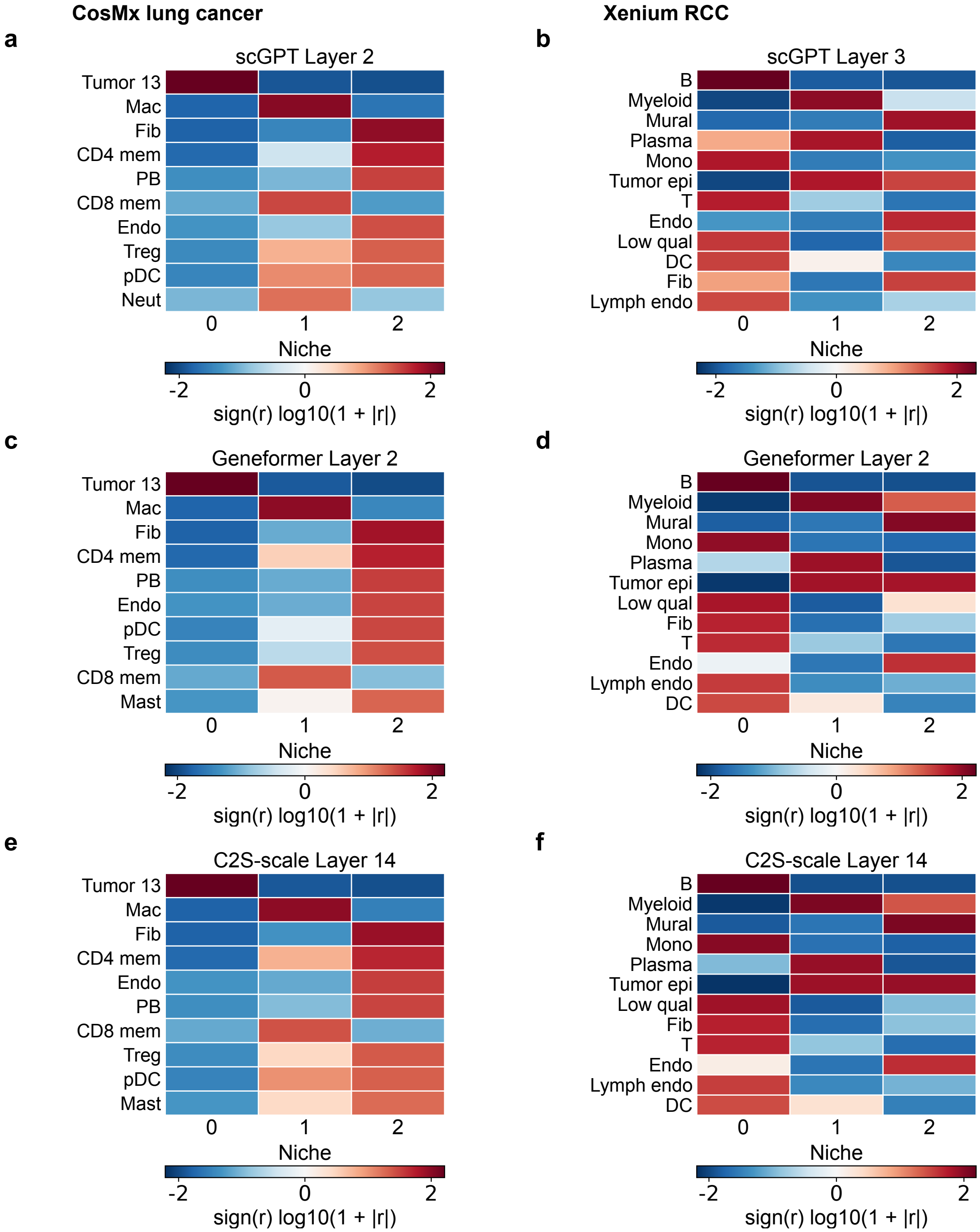


Supplementary Figure S1: **Niche cell-type residuals in CosMx lung cancer and Xenium renal cell carcinoma (RCC). a,c,e,** CosMx. **b,d,f,** Xenium with marker-based labels at three niches (Supplementary Figure S4). Rows show scGPT, Geneformer, and C2S-scale. Each heatmap shows the standardized cell-type enrichment residual r of each niche, plotted as sign(r) times log10(1 + |r|), for the ten CosMx and twelve Xenium labels with the largest positive residual. The color scale is symmetric within each panel. In CosMx, N0 is the tumor 13 niche, N1 the macrophage niche, and N2 the niche of fibroblasts with CD4 memory T, plasmablast, endothelial, Treg, and pDC cells. Niches are matched across models by composition (Methods). Abbreviations as in Figures 3 and 4.

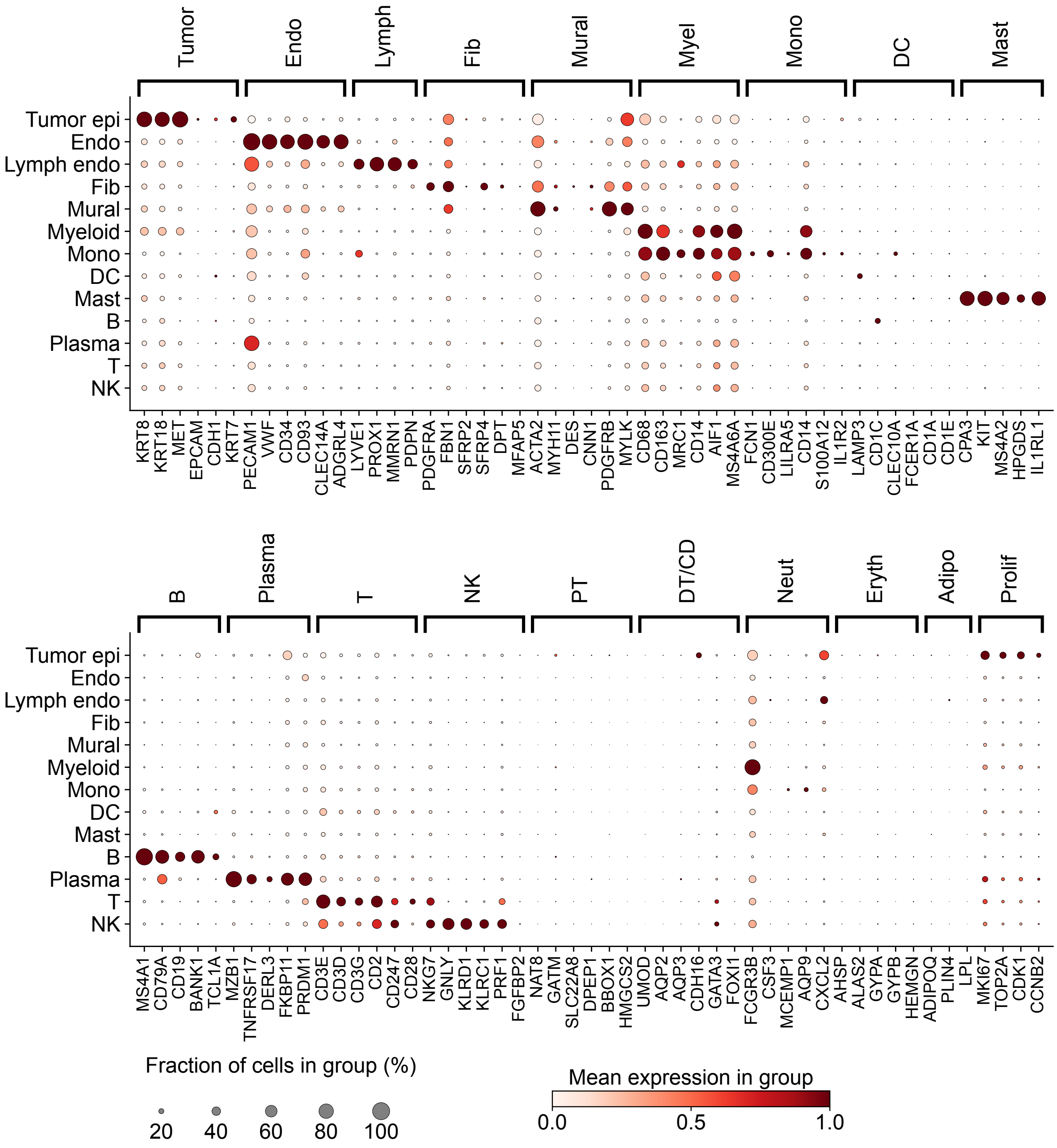


Supplementary Figure S2: **Marker-based annotation of the Xenium RCC section.** Expression of the curated marker sets, restricted to the 399-gene panel, across the Leiden clusters and their assigned labels (13 types). Low-quality cells (8.7% of cells; Methods) are not shown. Marker sets: Lymph, lymphatic endothelial; Myel, myeloid; Mono, monocyte; DC, dendritic cell; PT, proximal tubule; DT/CD, distal tubule/collecting duct; Neut, neutrophil; Eryth, erythrocyte; Adipo, adipocyte; Prolif, proliferation. Other abbreviations as in Figure 4.

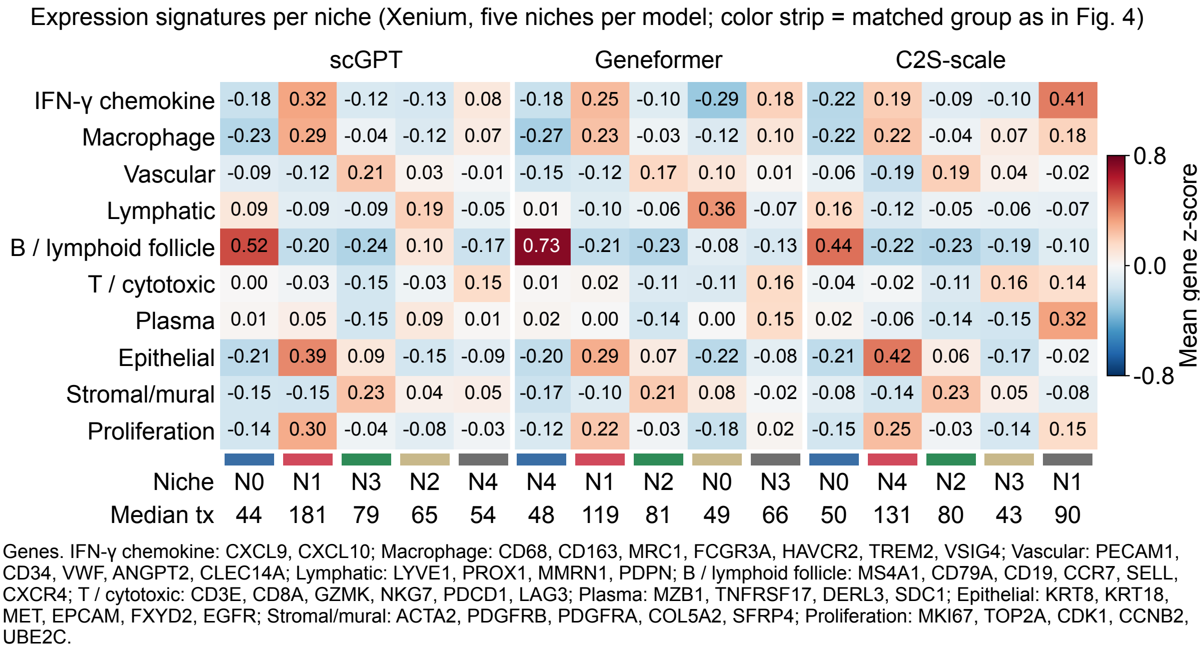


Supplementary Figure S3: **Expression signatures of the five Xenium RCC niches.** Mean gene z-score of each curated signature in each niche, at each model's five-niche configuration. The median number of transcripts per cell is given below each niche, and the signature genes are listed under the panel. Niches are ordered and colored by matched group as in Figure 4. Blue, red, and green mark the B/T-lymphoid, tumor/myeloid, and tumor/vascular niches shared by all three models. Sand and gray mark model-specific niches. Niche numbers are model-specific.

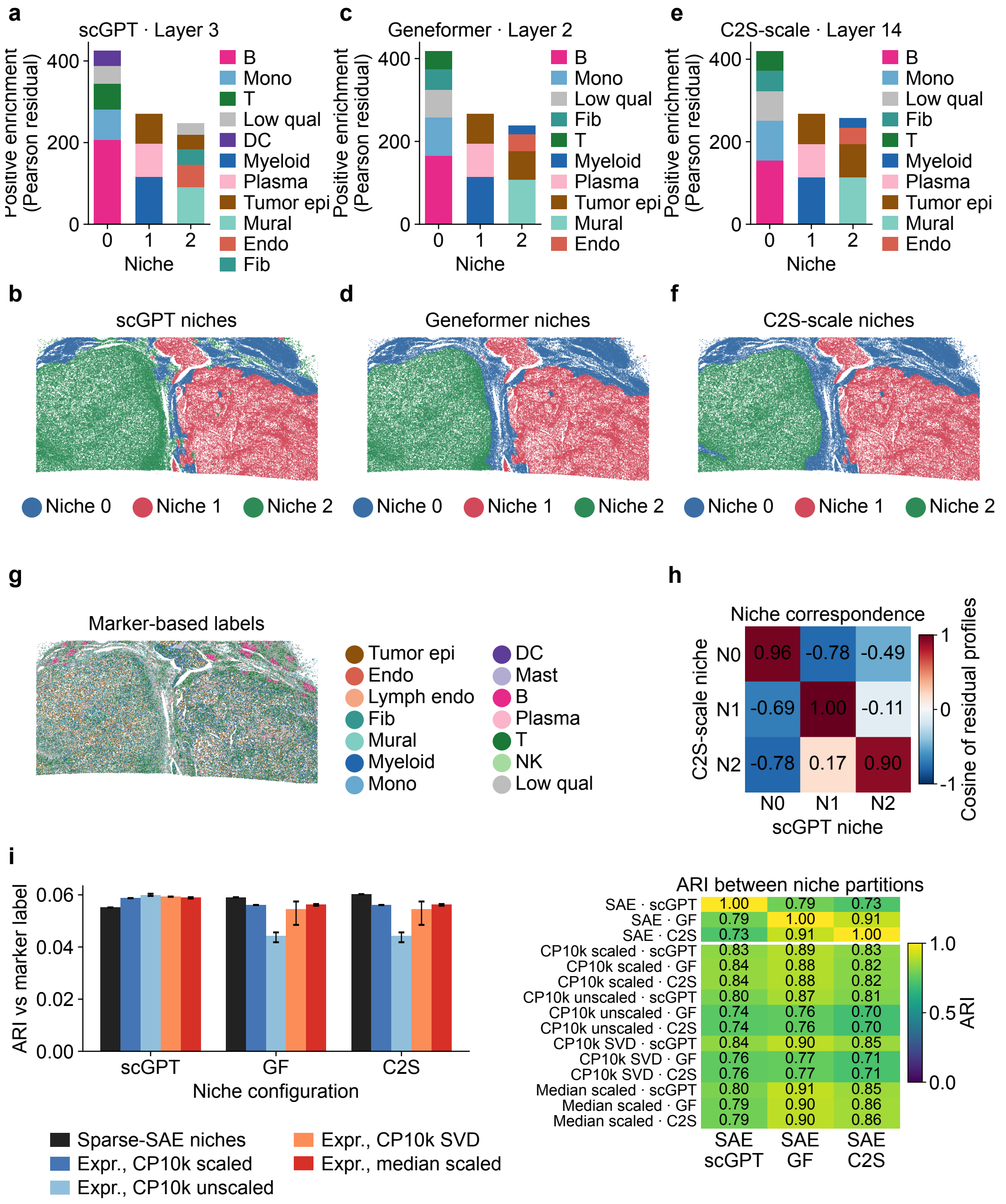


Supplementary Figure S4: **Xenium RCC niches at the highest-scoring configurations (three niches). a,b,** scGPT Layer 3. **c,d,** Geneformer Layer 2. **e,f,** C2S-scale Layer 14 (Methods; Supplementary Table S5). **g,** Whole-section orientation. **h,** Cosine similarity between the niche residual profiles of C2S-scale and scGPT (Supplementary Table S6a). **i,** Adjusted Rand index (ARI) of foundation-model and expression-only niches with the marker-based labels (left) and with each other (right). Panel layout and abbreviations follow Figure 4. Cell labels are marker-based (Supplementary Figure S2). The three territories are a B/T-lymphoid border, a myeloid/tumor/plasma territory, and a tumor/vascular-mural territory. They recur in all three models (cross-model ARI 0.73–0.91).

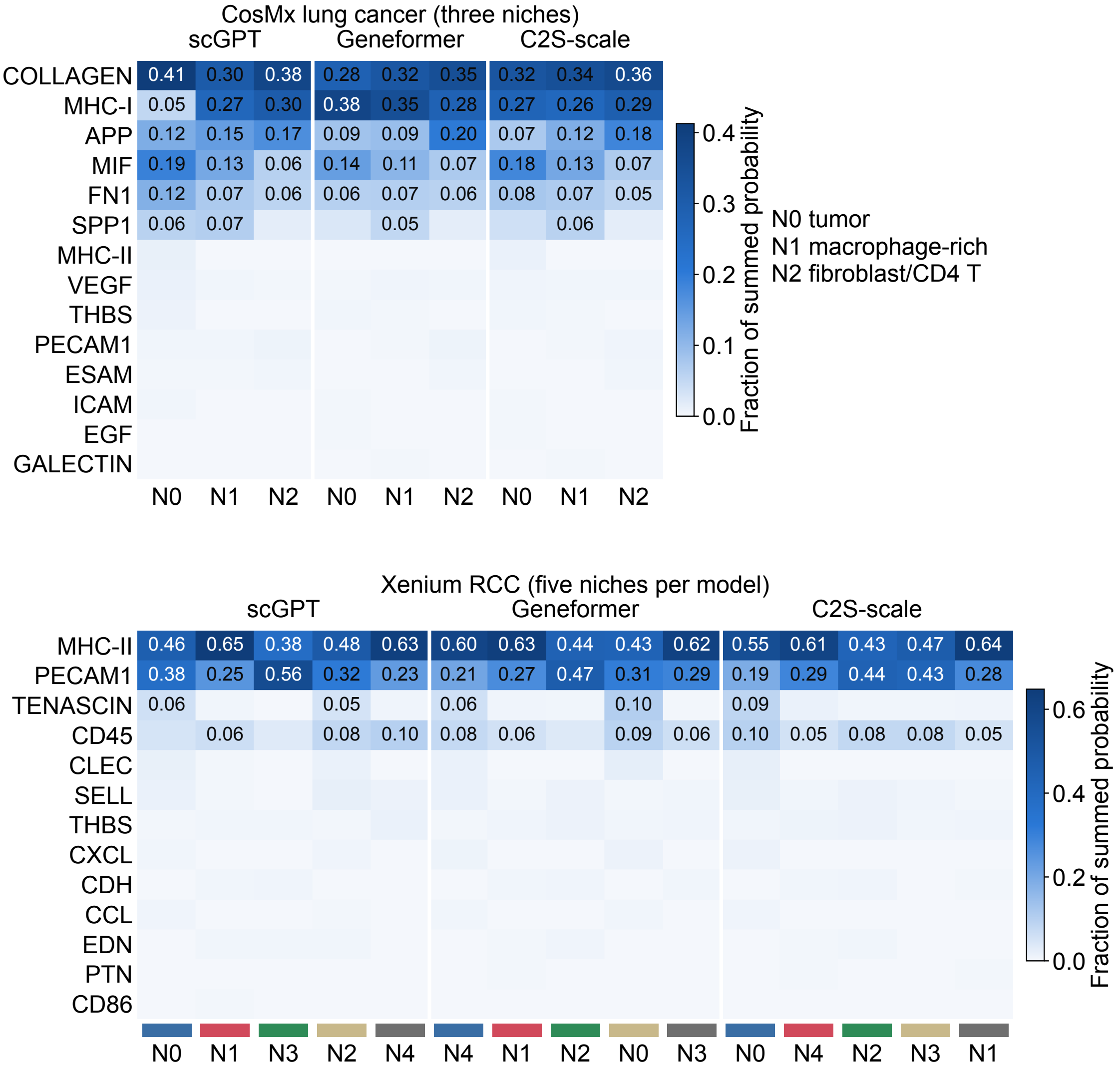


Color strip: matched group as in Fig. 4 (shared niches: blue B/T-lymphoid, red tumor/myeloid, green tumor/vascular; model-specific niches: sand and gray, matched by composition to Geneformer N0 and N3).
Values shown where the share is at least 0.05.

Supplementary Figure S5: **Niche-resolved CellChat pathway profiles (exploratory).** Share of the summed CellChat communication probability carried by each pathway, for each model and niche (Methods). CosMx is shown at three niches. Xenium is shown at five niches per model, ordered and colored by matched group as in Figure 4. The dominant pathways are shared by all niches, and niche-specific pathways have shares of at most 0.09. Agreement of the profiles between models is reported in Supplementary Table S19 (C1).

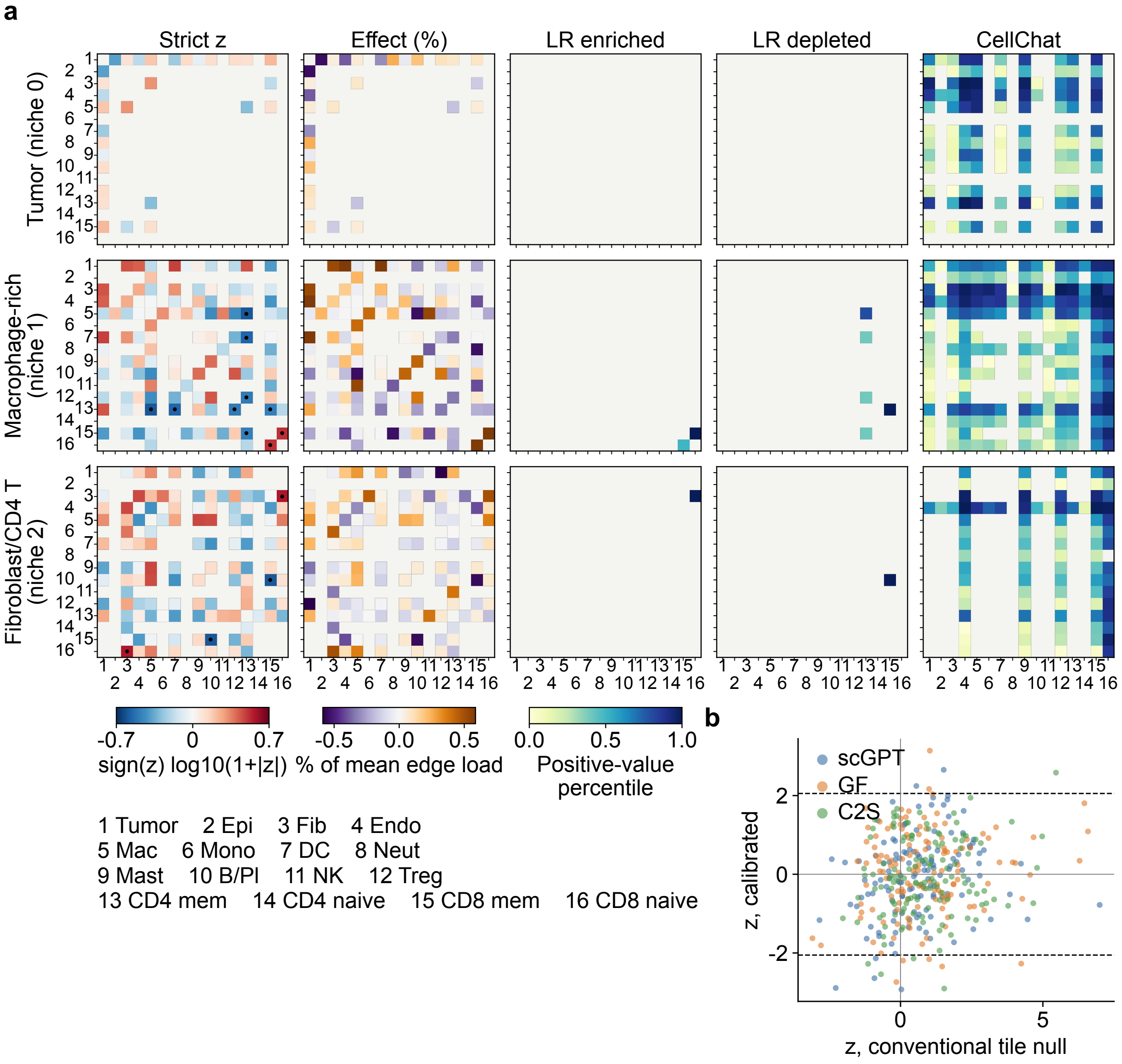


Supplementary Figure S6: **Calibrated interface statistic in CosMx lung cancer: full scGPT layout.** **a,** scGPT Layer 2. Rows are niches 0–2 (tumor, macrophage-rich, and fibroblast/CD4-T niche). The columns show the calibrated z as $\mathrm{sign}(z)\log_{10}(1+|z|)$, with dots where $|z| \geq z^* = 2.00$ (Methods), and the effect size as a percentage of the mean edge program load. They also show the term-supported ligand–receptor (LR) burden on pairs called enriched or depleted and the matched-niche CellChat weight. LR and CellChat values are within-panel percentiles of the positive values, so their absolute scales are not comparable. Cell types are numbered as in the key below the color bars. Light gray marks pairs not tested in that niche. **b,** z under a conventional, uncalibrated 400-unit (72-µm) tile null against the calibrated z, for every niche-by-pair test in the three models. Dashed lines mark $\pm z^*$ (2.05, the largest of the three models). Calibration and call counts are in Supplementary Table S9, and every call is in Supplementary Table S8. Abbreviations as in Figure 3.

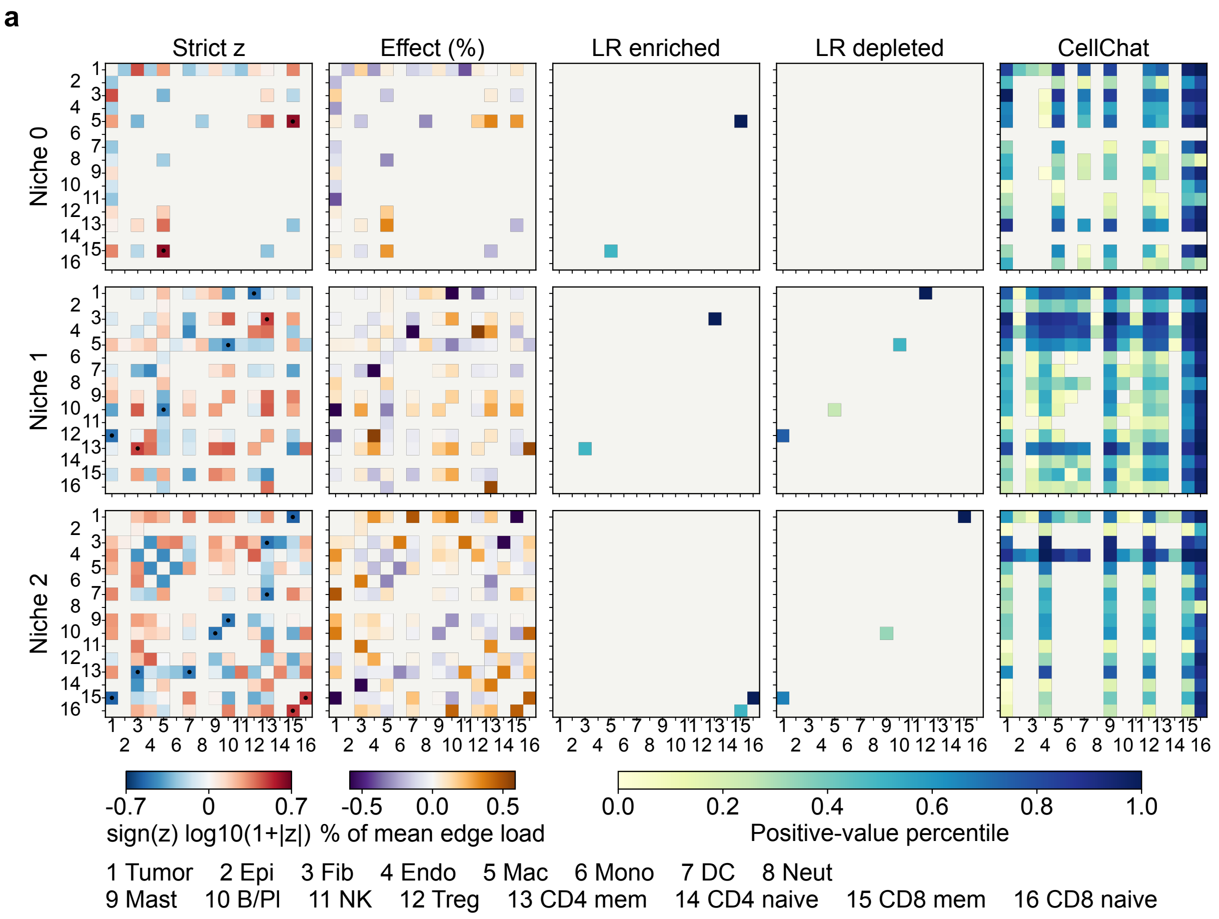


Supplementary Figure S7: **Calibrated interface statistic in CosMx lung cancer: Geneformer.** Geneformer Layer 2, niches 0–2. Layout and statistic as in Supplementary Figure S6a, with $z^* = 2.00$. Niche numbers are model-specific.

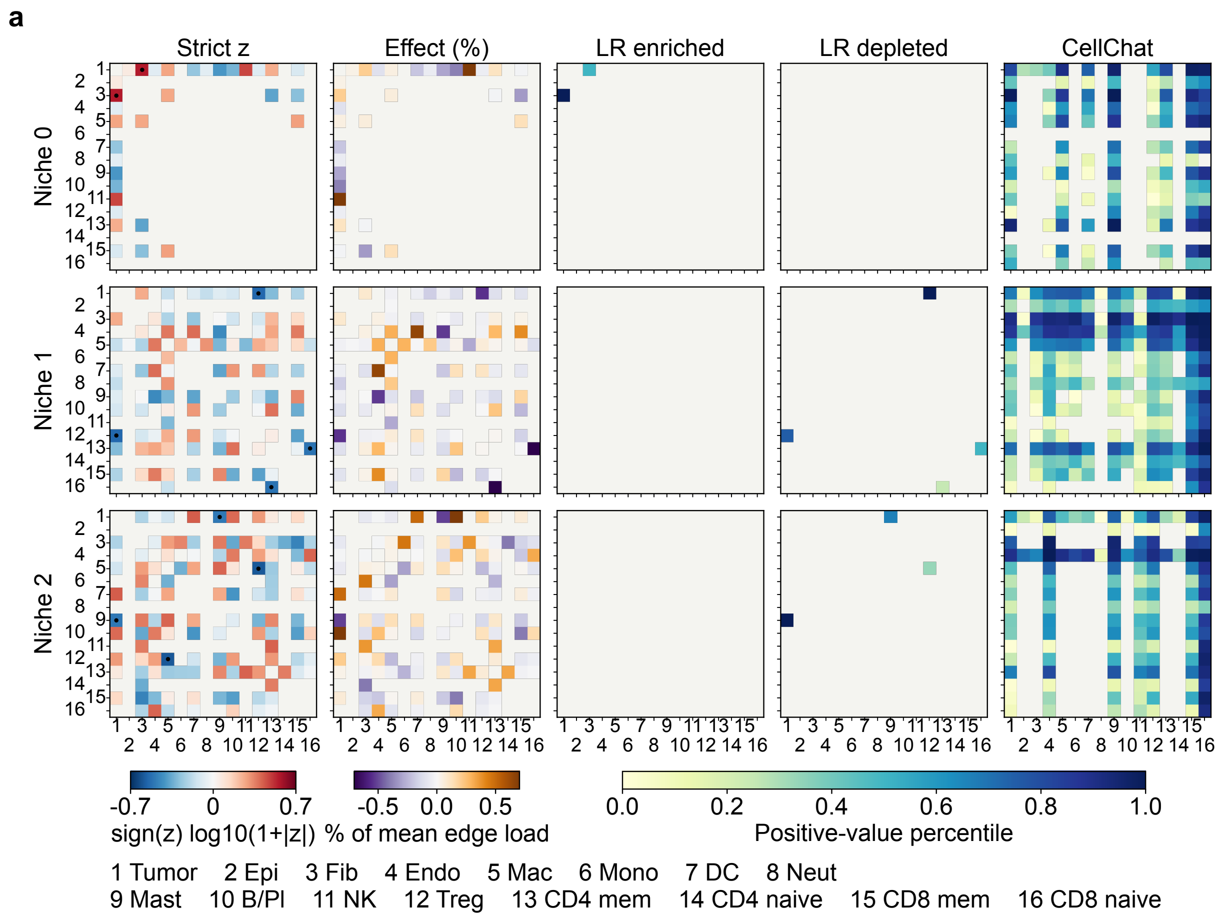


Supplementary Figure S8: **Calibrated interface statistic in CosMx lung cancer: C2S-scale.** C2S-scale Layer 14, niches 0–2. Layout and statistic as in Supplementary Figure S6a, with $z^* = 2.05$. Niche numbers are model-specific.

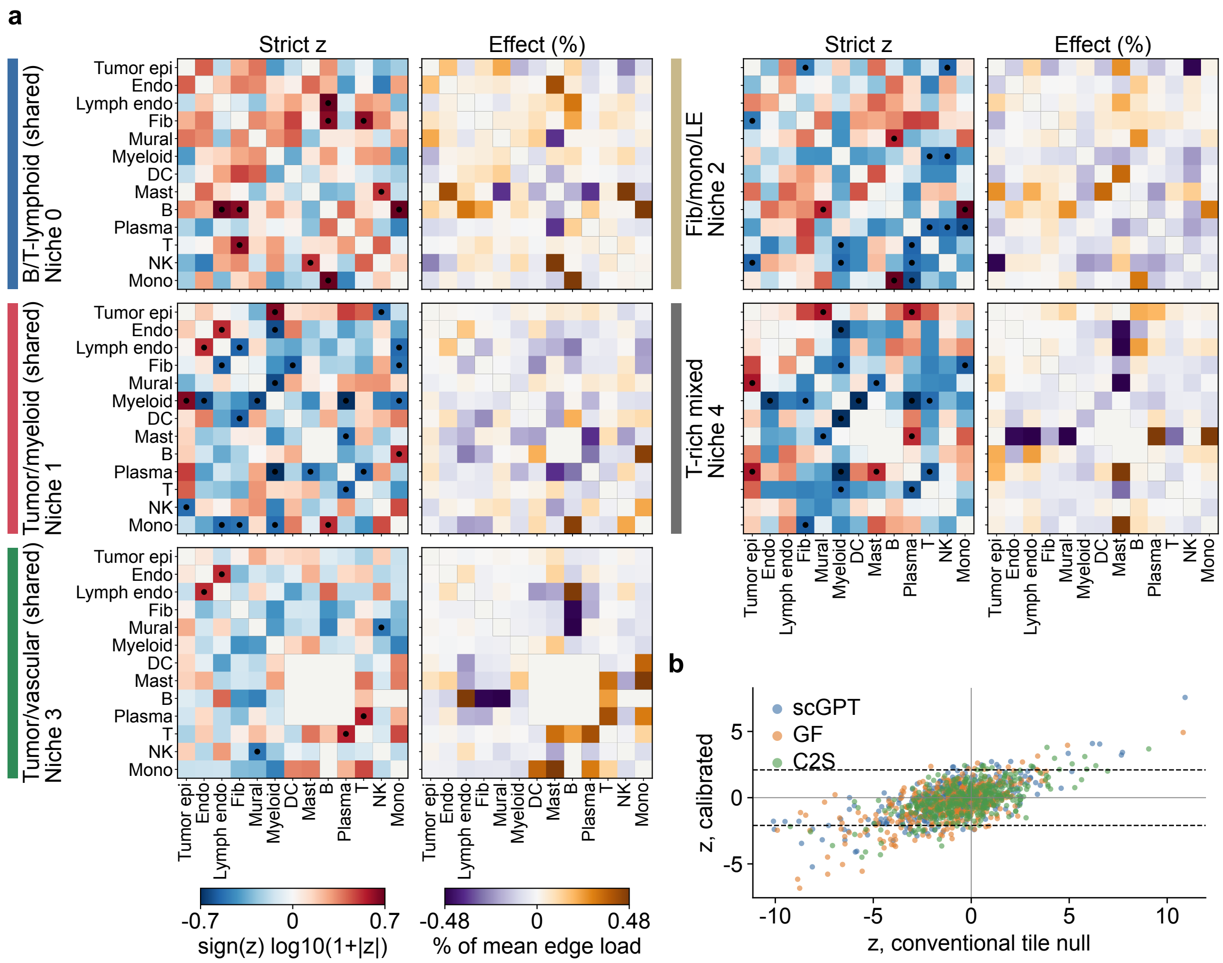


Supplementary Figure S9: **Calibrated interface statistic in Xenium RCC at five niches: full scGPT layout.** **a,** scGPT Layer 3 with marker-based cell labels. Rows are niches, with shared niches on the left and model-specific niches on the right. They are ordered and colored by matched group as in Figure 4 and carry their model-specific numbers. Columns show the calibrated z (dots, $|z| \geq z^* = 2.10$) and the effect size, as in Supplementary Figure S6a. The matched-niche CellChat rows are tabulated by call in Supplementary Table S10. Light gray marks pairs not tested in that niche. **b,** z under a conventional, uncalibrated 400-µm tile null against the calibrated z on the same labels, for every niche-by-pair test in the three models. Dashed lines mark $\pm z^*$ (2.10, the largest of the three models). Contact-level calls follow partner-marker transcripts at the contact, as expected from segmentation spillover [1, 2], and are not interpreted as interface biology. Abbreviations as in Figure 4. Fib/mono/LE, fibroblast/monocyte/lymphatic endothelial; GF, Geneformer; C2S, C2S-scale.

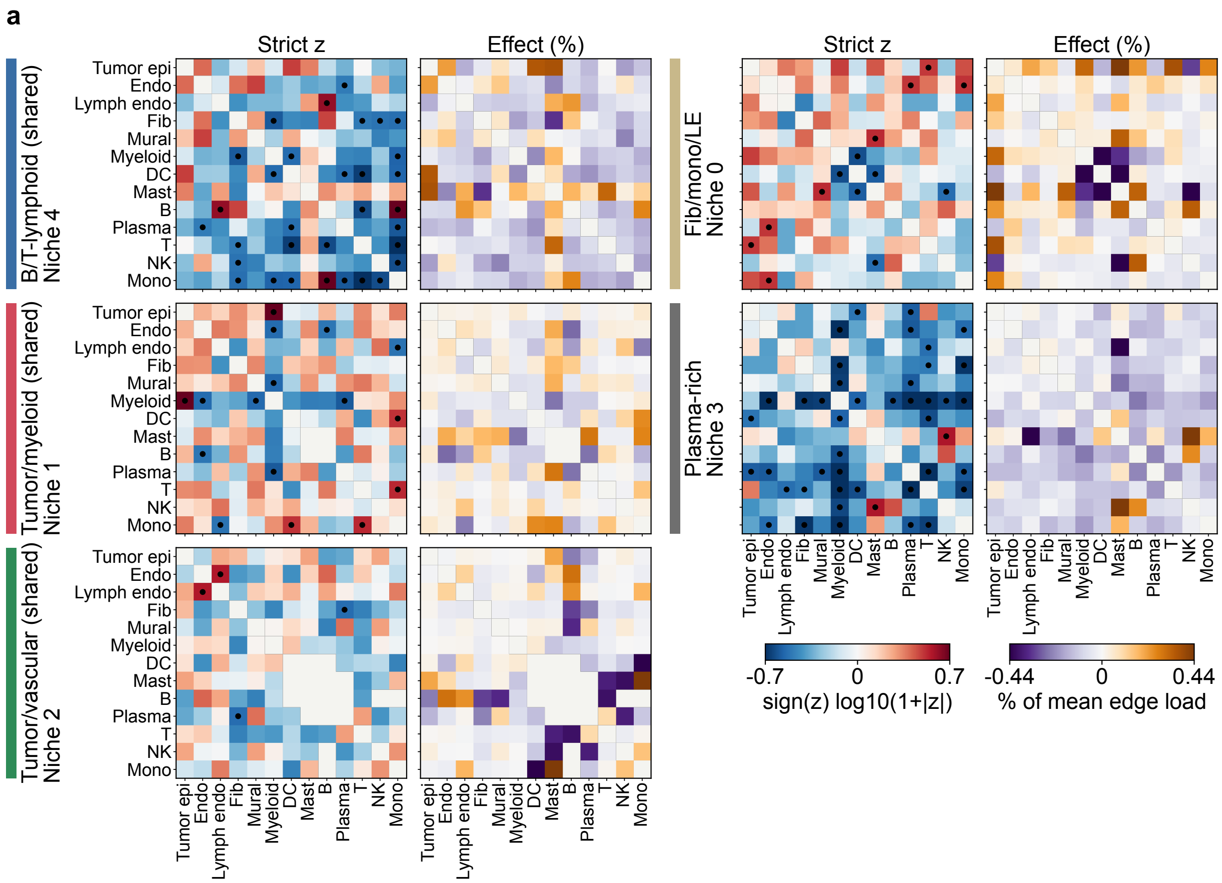


Supplementary Figure S10: **Calibrated interface statistic in Xenium RCC at five niches: Geneformer.** Geneformer Layer 2 with marker-based cell labels. Layout and statistic as in Supplementary Figure S9a, with $z^* = 2.06$. The note on contact-level calls in Supplementary Figure S9 applies here.

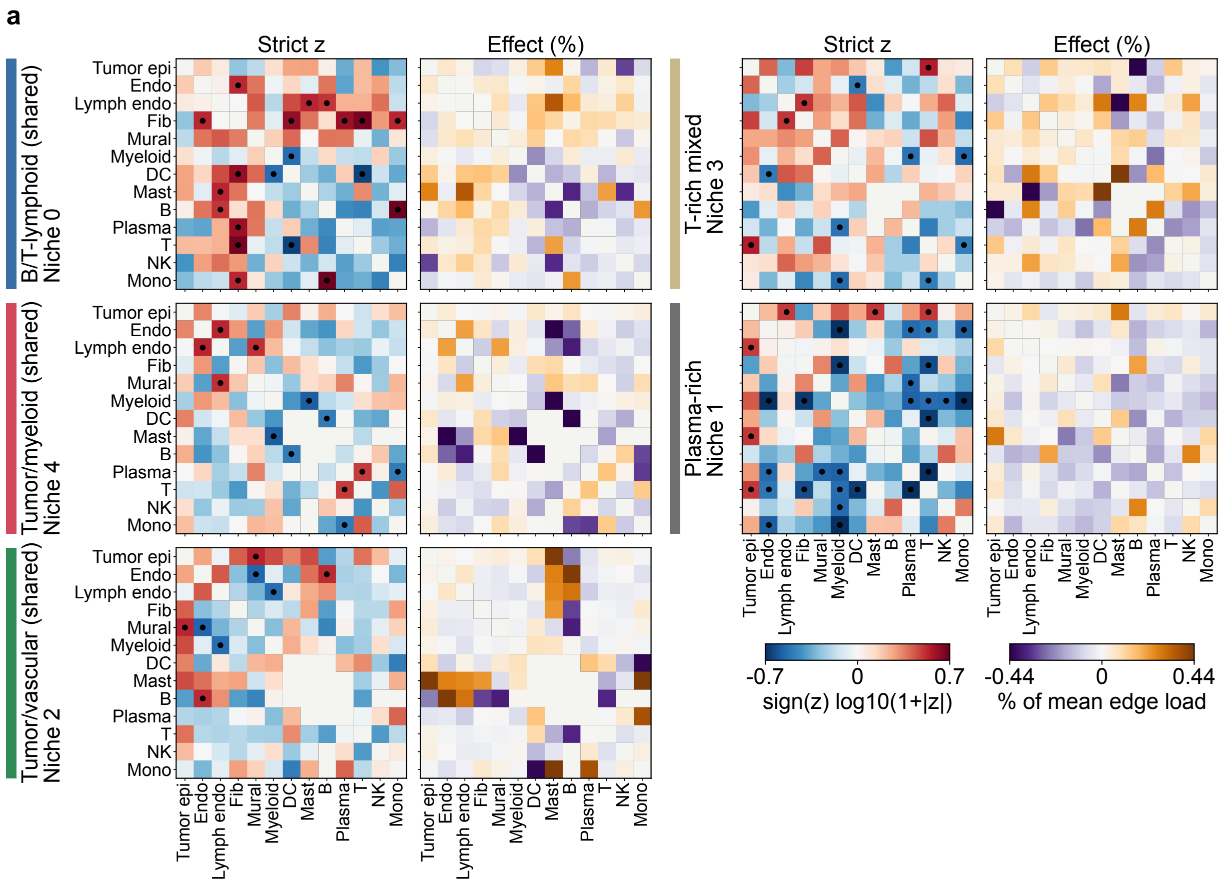


Supplementary Figure S11: **Calibrated interface statistic in Xenium RCC at five niches: C2S-scale.** C2S-scale Layer 14 with marker-based cell labels. Layout and statistic as in Supplementary Figure S9a, with $z^* = 2.05$. The note on contact-level calls in Supplementary Figure S9 applies here.

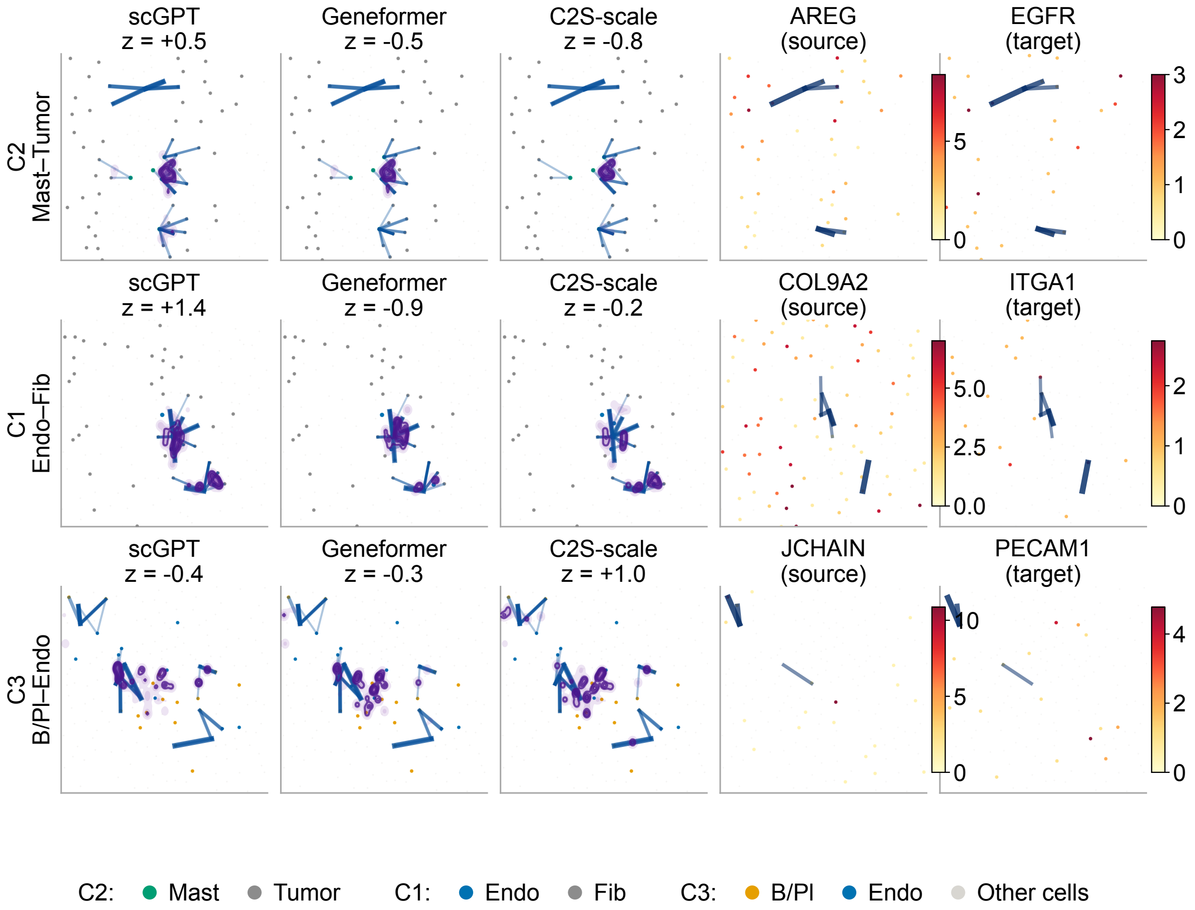


Supplementary Figure S12: **CosMx case crops.** Row labels give the case identifier (Supplementary Table S15) and the cell pair. C2 is mast–tumor AREG/EGFR, C1 is endothelial–fibroblast COL9A2/ITGA1, and C3 is B/plasma–endothelial JCHAIN/PECAM1. The three model columns aggregate all term-supported candidate LR expression/proximity links for the pair and crop. The two expression columns show the source and target gene as log1p transcript counts, with blue links restricted to the displayed gene pair. The key below gives the cell-type colors. In C2 and C1 the more numerous type is gray. Panel titles give the niche-free calibrated z for the pair ($z^* = 2.00$). The crops were chosen for display before the proximity tests, which are reported in Supplementary Figure S13b and Supplementary Table S15. JCHAIN and PECAM1 are cell-type markers, shown as a proximity context. ITGA1 is one component of the nominated ITGA1–ITGB1 complex. Abbreviations as in Figure 3.

**a**

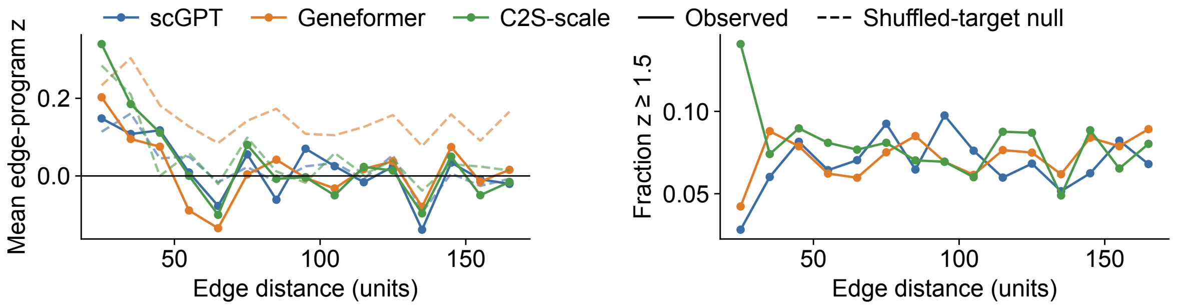


**b** CosMx, author labels

| Crop | Genes | Pair | Src+ | Tgt+ | Null | Obs. | Mean null | Fold | z | p (enr / dep) |
|---|---|---|---|---|---|---|---|---|---|---|
| C2 | AREG/EGFR | Mast–Tumor | 2 | 25 | position | 3 | 4.0 | 0.80 | -0.6 | 0.822 / 0.394 |
| C2 | AREG/EGFR | Mast–Tumor | 2 | 25 | within | 3 | 2.5 | 1.14 | 0.3 | 0.470 / 0.762 |
| C1 | COL9A2/ITGA1 | Endo–Fib | 5 | 8 | position | 4 | 3.1 | 1.22 | 0.4 | 0.399 / 0.743 |
| C1 | COL9A2/ITGA1 | Endo–Fib | 5 | 8 | within | 4 | 4.5 | 0.91 | -0.2 | 0.613 / 0.534 |
| C3 | JCHAIN/PECAM1 | B/Pl–Endo | 8 | 7 | position | 1 | 3.0 | 0.50 | -1.0 | 0.916 / 0.250 |
| C3 | JCHAIN/PECAM1 | B/Pl–Endo | 8 | 7 | within | 1 | 3.1 | 0.49 | -1.0 | 0.910 / 0.245 |

**c** Xenium, marker-based labels

| Crop | Genes | Pair | Src+ | Tgt+ | Null | Obs. | Mean null | Fold | z | p (enr / dep) |
|---|---|---|---|---|---|---|---|---|---|---|
| X2 | EDN1/EDNRB | Endo–Tumor epi | 121 | 170 | position | 492 | 411.8 | 1.19 | 3.5 | 0.001 / 1.000 |
| X2 | EDN1/EDNRB | Endo–Tumor epi | 121 | 170 | within | 492 | 418.5 | 1.18 | 2.6 | 0.011 / 0.992 |

Xenium, transferred labels (reference)

| Crop | Genes | Pair | Src+ | Tgt+ | Null | Obs. | Mean null | Fold | z | p (enr / dep) |
|---|---|---|---|---|---|---|---|---|---|---|
| X2 | EDN1/EDNRB | Endo–PT-like | 251 | 71 | position | 435 | 366.1 | 1.19 | 2.5 | 0.010 / 0.991 |
| X2 | EDN1/EDNRB | Endo–PT-like | 251 | 71 | within | 435 | 350.6 | 1.24 | 3.5 | 0.003 / 0.998 |

Supplementary Figure S13: **Diagnostics for the case studies.** **a,** The CosMx endothelial–fibroblast edge program of Figure 7a by cell–cell distance, in bins of 10 units (1.8 µm). Left, observed mean edge-program z-score and a coordinate-shuffled target-cell null. Right, share of observed edges with $z \geq 1.5$. Bins with fewer than 50 edges are omitted. **b,** Gene-pair proximity within each CosMx crop of Supplementary Figure S12 at 100 units (18 µm). Observed pairs of ligand-positive source and receptor-positive target cells are compared with a target-position null and a within-type null. **c,** The same test for the Xenium EDN1/EDNRB crop of Figure 8c (X2) at 100 µm, on marker-based labels (top) and, for reference, on transferred labels (bottom). PT-like, cells assigned to the reference proximal-tubule class (Supplementary Table S7); Src+ and Tgt+, ligand-positive source and receptor-positive target cells in the crop; Obs., pairs within 100 units; p, upper- and lower-tail permutation P (1,000 permutations). Abbreviations as in Figures 3 and 4.

#### 4 Supplementary Tables

Supplementary Table S1: **Concept alignment and population specificity at the analysis layers.** Concept F1 is the test F1 of the feature selected on validation data, averaged over concepts with 5–100 panel genes (Methods). In the row split, gene-token rows are held out and the same genes occur in validation and test. In the whole-gene split, genes are held out. Gini specificity index (GSI) excess and marker-gene hit rate (MHR@100) are computed as in Figure 2g,h, on the same cells for all representations. Features kept is the number of sparse-SAE features retained for the spatial analyses. Layer in code is the zero-based layer identifier. TIS, transcriptomic interpretability score.

| | | | | | Concept F1, row split | | | Concept F1, whole-gene split | | | GSI excess | | | MHR@100 | | | | |
|---|---|---|---|---|---|---|---|---|---|---|---|---|---|---|---|---|---|---|
| Dataset | Model | Layer in code | Layer | TIS gain | Raw | Untrained SAE | Sparse SAE | Raw | Untrained SAE | Sparse SAE | Raw | Sparse SAE | Genes | Raw | Sparse SAE | Genes | Chance | Features kept |
| CosMx lung cancer | scGPT | layer_1.norm2 | 2 | 0.176 | 0.175 | 0.230 | 0.339 | 0.033 | 0.027 | 0.021 | 0.063 | 0.128 | 0.195 | 0.465 | 0.579 | 0.567 | 0.104 | 194 |
| CosMx lung cancer | Geneformer | layer_1 | 2 | 0.171 | 0.110 | 0.166 | 0.349 | 0.054 | 0.040 | 0.017 | 0.079 | 0.146 | 0.195 | 0.503 | 0.624 | 0.572 | 0.104 | 261 |
| CosMx lung cancer | C2S-scale | layer_13 | 14 | 0.169 | 0.107 | 0.169 | 0.317 | 0.067 | 0.075 | 0.025 | 0.091 | 0.169 | 0.195 | 0.553 | 0.685 | 0.572 | 0.104 | 170 |
| Xenium RCC | scGPT | layer_2.norm2 | 3 | 0.214 | 0.319 | 0.400 | 0.462 | 0.091 | 0.075 | 0.029 | 0.115 | 0.256 | 0.262 | 0.766 | 0.826 | 0.789 | 0.251 | 118 |
| Xenium RCC | Geneformer | layer_1 | 2 | 0.195 | 0.261 | 0.342 | 0.450 | 0.103 | 0.103 | 0.006 | 0.115 | 0.206 | 0.267 | 0.808 | 0.823 | 0.793 | 0.251 | 125 |
| Xenium RCC | C2S-scale | layer_13 | 14 | 0.190 | 0.253 | 0.321 | 0.411 | 0.115 | 0.091 | 0.032 | 0.096 | 0.232 | 0.267 | 0.811 | 0.841 | 0.793 | 0.251 | 94 |

Randomly re-initialized models (Methods): CosMx Layer 2, 30,356 scGPT and 40,480 Geneformer cells, sparse SAE with the same selection rule. Trained-model values are for the same cells.

| Model | Weights | Concept F1, row split | Concept F1, whole-gene split | Concepts with test F1 $\geq 0.5$ |
|---|---|---|---|---|
| scGPT | Trained | 0.333 | 0.023 | 275 |
| scGPT | Re-initialized except the gene-embedding table | 0.366 | 0.013 | 643 |
| scGPT | All weights re-initialized | 0.330 | 0.008 | 350 |
| Geneformer | Trained | 0.355 | 0.017 | 559 |
| Geneformer | Re-initialized except the gene-embedding table | 0.349 | 0.004 | 638 |
| Geneformer | All weights re-initialized | 0.341 | 0.001 | 636 |

Concepts marked at the analysis layers (row split): the number of concepts whose selected unit reached a test F1 $\geq 0.5$ (same concepts as Concept F1), and the number of distinct sparse features that mark them. Validation F1 (whole-gene split) is the mean validation F1 of the sparse feature selected for each concept; its test counterpart is the whole-gene Concept F1 above.

| | | | Row split: concepts with test F1 $\geq 0.5$ | | | | Whole-gene split, sparse SAE |
|---|---|---|---|---|---|---|---|
| Dataset | Model | Layer | Raw | Untrained SAE | Sparse SAE | Sparse features marking them | Validation F1 |
| CosMx lung cancer | scGPT | 2 | 2 | 35 | 315 | 63 | 0.673 |
| CosMx lung cancer | Geneformer | 2 | 0 | 0 | 484 | 111 | 0.662 |
| CosMx lung cancer | C2S-scale | 14 | 0 | 3 | 262 | 63 | 0.618 |
| Xenium RCC | scGPT | 3 | 70 | 249 | 593 | 95 | 0.717 |
| Xenium RCC | Geneformer | 2 | 33 | 109 | 540 | 95 | 0.711 |
| Xenium RCC | C2S-scale | 14 | 40 | 80 | 342 | 66 | 0.679 |

Supplementary Table S2: **Causal steering check of concept-labeled sparse features in CosMx.** The protocol is given in the Supplementary Methods. $D_O$ is the median over features of |AUROC−0.5|, where the area under the receiver operating characteristic curve (AUROC) compares the output change of the concept's other member genes with that of non-member genes. The contrast columns give $D_O$ of the steered feature minus $D_O$ of each control. A median $\text{AUROC}_O$ above 0.5 means that the other member genes moved up relative to non-member genes. Brackets give 95% bootstrap CIs over cells (1,000 resamples). The model output differs between models, so values are compared within a model only. MVC, the scGPT decoder that reads the cell embedding.

| Model | Layer | Cells | Model output | Features | $D_O$, steered | Steered − random feature | Steered − random direction | Steered − gene identity | Median $\text{AUROC}_O$, steered |
|---|---|---|---|---|---|---|---|---|---|
| C2S-scale | Layer 14 | 40 | Next gene | 61 | 0.183<br>[0.160, 0.211] | +0.056<br>[0.026, 0.082] | +0.067<br>[0.031, 0.102] | −0.005<br>[−0.034, 0.031] | 0.622<br>[0.594, 0.648] |
| scGPT | Layer 2 | 200 | MVC expression | 63 | 0.138<br>[0.133, 0.141] | +0.030<br>[0.022, 0.035] | +0.025<br>[0.016, 0.027] | +0.003<br>[−0.006, 0.006] | 0.556<br>[0.545, 0.561] |
| Geneformer | Layer 2 | 200 | Masked gene | 110 | 0.126<br>[0.117, 0.138] | −0.013<br>[−0.021, 0.005] | +0.005<br>[−0.007, 0.023] | −0.008<br>[−0.017, 0.007] | 0.528<br>[0.520, 0.536] |

Supplementary Table S3: **Linear probe of concept membership for unseen genes.** Values are test-gene AUROC (mean ± SD over five whole-gene splits) for concepts with 5–100 panel genes. The probe, the best single dimension, and the controls are defined in the Methods. The standardized version is the primary analysis, and the center-only version is a post hoc sensitivity analysis. †Post hoc rows, with standardized inputs only (–, not computed). Symbol component removed, the representation minus its least-squares fit on the first 50 principal components (PCs) of the gene-symbol trigram profile.

| Dataset | Model | Representation | Linear probe (standardized) | Linear probe (center-only) | Best single dimension |
|---|---|---|---|---|---|
| CosMx lung cancer | scGPT L2 | Raw hidden state | 0.678 ± 0.026 | 0.685 ± 0.024 | 0.588 ± 0.009 |
| CosMx lung cancer | scGPT L2 | Sparse SAE | 0.636 ± 0.018 | 0.680 ± 0.026 | 0.571 ± 0.014 |
| CosMx lung cancer | Geneformer L2 | Raw hidden state | 0.681 ± 0.015 | 0.684 ± 0.014 | 0.581 ± 0.006 |
| CosMx lung cancer | Geneformer L2 | Sparse SAE | 0.620 ± 0.018 | 0.659 ± 0.014 | 0.560 ± 0.009 |
| CosMx lung cancer | C2S-scale L14 | Raw hidden state | 0.745 ± 0.006 | 0.740 ± 0.006 | 0.634 ± 0.003 |
| CosMx lung cancer | C2S-scale L14 | Sparse SAE | 0.666 ± 0.013 | 0.732 ± 0.014 | 0.605 ± 0.010 |
| Xenium RCC | scGPT L3 | Raw hidden state | 0.662 ± 0.025 | 0.662 ± 0.026 | 0.562 ± 0.016 |
| Xenium RCC | scGPT L3 | Sparse SAE | 0.592 ± 0.029 | 0.630 ± 0.026 | 0.536 ± 0.011 |
| Xenium RCC | Geneformer L2 | Raw hidden state | 0.666 ± 0.027 | 0.669 ± 0.025 | 0.579 ± 0.004 |
| Xenium RCC | Geneformer L2 | Sparse SAE | 0.608 ± 0.038 | 0.636 ± 0.015 | 0.571 ± 0.020 |
| Xenium RCC | C2S-scale L14 | Raw hidden state | 0.678 ± 0.036 | 0.673 ± 0.036 | 0.583 ± 0.014 |
| Xenium RCC | C2S-scale L14 | Sparse SAE | 0.613 ± 0.030 | 0.680 ± 0.033 | 0.571 ± 0.041 |
| CosMx lung cancer | scGPT L2 | Re-initialized except gene embeddings† | 0.668 ± 0.020 | – | – |
| CosMx lung cancer | scGPT L2 | Re-initialized, all weights† | 0.500 ± 0.013 | – | – |
| CosMx lung cancer | Geneformer L2 | Re-initialized except gene embeddings† | 0.660 ± 0.009 | – | – |
| CosMx lung cancer | Geneformer L2 | Re-initialized, all weights† | 0.518 ± 0.015 | – | – |
| CosMx lung cancer | scGPT L2 | Raw, symbol component removed† | 0.616 ± 0.027 | – | – |
| CosMx lung cancer | Geneformer L2 | Raw, symbol component removed† | 0.616 ± 0.013 | – | – |
| CosMx lung cancer | C2S-scale L14 | Raw, symbol component removed† | 0.657 ± 0.012 | – | – |
| Xenium RCC | scGPT L3 | Raw, symbol component removed† | 0.594 ± 0.025 | – | – |
| Xenium RCC | Geneformer L2 | Raw, symbol component removed† | 0.583 ± 0.029 | – | – |
| Xenium RCC | C2S-scale L14 | Raw, symbol component removed† | 0.606 ± 0.039 | – | – |
| CosMx lung cancer | Control | Co-expression profile (50 PCs) | 0.603 ± 0.022 | 0.638 ± 0.026 | 0.575 ± 0.018 |
| CosMx lung cancer | Control | Co-expression profile (all genes) | 0.590 ± 0.021 | 0.634 ± 0.028 | 0.608 ± 0.017 |
| CosMx lung cancer | Control | Cell-type expression profile | 0.608 ± 0.025 | 0.609 ± 0.025 | 0.593 ± 0.016 |
| CosMx lung cancer | Control | Mean and detection rate | 0.558 ± 0.012 | 0.537 ± 0.013 | 0.547 ± 0.003 |
| CosMx lung cancer | Control | Random vectors (512 dims) | 0.494 ± 0.009 | 0.493 ± 0.009 | 0.501 ± 0.006 |
| CosMx lung cancer | Control | Gene symbol, character trigrams† | 0.684 ± 0.014 | – | – |
| CosMx lung cancer | Control | Gene symbol (50 PCs)† | 0.676 ± 0.008 | – | – |
| CosMx lung cancer | Control | Co-expression (50 PCs), symbol component removed† | 0.574 ± 0.018 | – | – |
| Xenium RCC | Control | Co-expression profile (50 PCs) | 0.589 ± 0.013 | 0.621 ± 0.017 | 0.564 ± 0.014 |
| Xenium RCC | Control | Co-expression profile (all genes) | 0.593 ± 0.026 | 0.629 ± 0.021 | 0.595 ± 0.007 |
| Xenium RCC | Control | Cell-type expression profile | 0.597 ± 0.011 | 0.599 ± 0.011 | 0.597 ± 0.023 |
| Xenium RCC | Control | Mean and detection rate | 0.556 ± 0.023 | 0.551 ± 0.014 | 0.557 ± 0.019 |
| Xenium RCC | Control | Random vectors (512 dims) | 0.494 ± 0.023 | 0.496 ± 0.021 | 0.503 ± 0.013 |
| Xenium RCC | Control | Gene symbol, character trigrams† | 0.639 ± 0.042 | – | – |
| Xenium RCC | Control | Gene symbol (50 PCs)† | 0.587 ± 0.038 | – | – |
| Xenium RCC | Control | Co-expression (50 PCs), symbol component removed† | 0.557 ± 0.010 | – | – |

Supplementary Table S4: **Cross-donor replication of the CosMx analyses.** One section from each of four further donors was processed with the pipeline of the analyzed section (Lung13) unchanged. The replication criteria are given in the Methods. †, post hoc. **a,** Population specificity at the representative layers, as in Figure 2g–i. Sparse/genes is the sparse GSI excess as a share of the single-gene value. **b,** ARI of niches with author cell types. Cross-model ARI is the range over the three model pairs. Model–expression ARI is the median over models of the ARI with expression-only niches. **c,** Niches of the analyzed section that recur in all four donors, with the 22 author labels and with tumor subtypes merged. Niche 0 is the tumor niche. **d,** Prediction of the T-cell and myeloid shares of a macrophage's neighborhood from its own profile (ridge $R^2$, penalty chosen by inner cross-validation). Lung12 has fewer than 3,000 macrophages and is excluded from the replication criterion. Panels d–g follow on the next pages.

**a** Cell-level population specificity

| | | GSI excess | | | | MHR@100 | | | |
|---|---|---|---|---|---|---|---|---|---|
| Section | Model | Raw | Sparse | Genes | Sparse/genes | Raw | Sparse | Genes | Chance |
| Lung13 (analyzed) | scGPT | 0.063 | 0.128 | 0.195 | 66% | 0.465 | 0.579 | 0.567 | 0.104 |
| | Geneformer | 0.079 | 0.146 | 0.195 | 75% | 0.503 | 0.624 | 0.572 | 0.104 |
| | C2S-scale | 0.091 | 0.169 | 0.195 | 87% | 0.553 | 0.685 | 0.572 | 0.104 |
| Lung5 | scGPT | 0.061 | 0.133 | 0.192 | 69% | 0.654 | 0.727 | 0.839 | 0.104 |
| | Geneformer | 0.077 | 0.151 | 0.192 | 79% | 0.705 | 0.784 | 0.789 | 0.104 |
| | C2S-scale | 0.086 | 0.190 | 0.192 | 99% | 0.733 | 0.838 | 0.789 | 0.104 |
| Lung6 | scGPT | 0.072 | 0.154 | 0.221 | 69% | 0.599 | 0.615 | 0.698 | 0.104 |
| | Geneformer | 0.087 | 0.170 | 0.222 | 77% | 0.644 | 0.729 | 0.709 | 0.104 |
| | C2S-scale | 0.091 | 0.177 | 0.222 | 80% | 0.668 | 0.767 | 0.709 | 0.104 |
| Lung9 | scGPT | 0.061 | 0.119 | 0.190 | 62% | 0.539 | 0.604 | 0.562 | 0.104 |
| | Geneformer | 0.073 | 0.130 | 0.191 | 68% | 0.558 | 0.674 | 0.550 | 0.104 |
| | C2S-scale | 0.077 | 0.146 | 0.191 | 76% | 0.612 | 0.728 | 0.550 | 0.104 |
| Lung12 | scGPT | 0.048 | 0.073 | 0.127 | 57% | 0.465 | 0.593 | 0.565 | 0.104 |
| | Geneformer | 0.054 | 0.087 | 0.128 | 68% | 0.531 | 0.618 | 0.557 | 0.104 |
| | C2S-scale | 0.062 | 0.141 | 0.128 | 110% | 0.596 | 0.717 | 0.557 | 0.104 |

**b** Niche agreement

| | ARI with author cell types | | | | | | |
|---|---|---|---|---|---|---|---|
| Section | scGPT | Geneformer | C2S-scale | Expression-only | Label composition | Cross-model ARI | Model–expression ARI (median) |
| Lung13 (analyzed) | 0.42 | 0.34 | 0.38 | 0.30 | 0.27 | 0.63–0.69 | 0.45 |
| Lung5 | 0.22 | 0.22 | 0.22 | 0.21 | 0.14 | 0.68–0.75 | 0.28 |
| Lung6 | 0.80 | 0.30 | 0.84 | 0.37 | 0.76 | 0.32–0.86 | 0.41 |
| Lung9 | 0.32 | 0.59 | 0.57 | 0.59 | 0.53 | 0.50–0.81 | 0.72 |
| Lung12 | 0.20 | 0.11 | 0.18 | 0.23 | 0.12 | 0.21–0.45 | 0.36 |

**c** Composition-level recurrence of the analyzed section's niches in all four donors

| Partition | 22 labels | Tumor merged | Per niche, tumor merged (Y/n in Lung5, 6, 9, 12) |
|---|---|---|---|
| Sparse SAE, scGPT | 0 of 3 | 1 of 3 | 0:YYYY, 1:nYnn, 2:YYYn |
| Sparse SAE, Geneformer | 0 of 3 | 0 of 3 | 0:YnYY, 1:nnnn, 2:YnYn |
| Sparse SAE, C2S-scale | 0 of 3 | 1 of 3 | 0:YYYY, 1:nYnn, 2:YnYn |
| Expression-only, scGPT configuration† | 0 of 3 | 1 of 3 | 0:YYYY, 1:nnnn, 2:nnnn |
| Expression-only, Geneformer configuration† | 0 of 3 | 0 of 3 | 0:YYYn, 1:nnnn, 2:nnnn |
| Expression-only, C2S-scale configuration† | 0 of 3 | 0 of 3 | 0:YYYn, 1:nnnn, 2:nnnn |
| Expression-only, sweep winner† | 0 of 3 | 1 of 3 | 0:YYYY, 1:nnnn, 2:nnnn |
| Label composition† | 0 of 3 | 0 of 3 | 0:nYnn, 1:YnYY, 2:nnYn |

Best-match cosines (post hoc) for sparse-SAE niches were 0.18–0.72 with 22 labels and 0.43–0.92 with tumor subtypes merged. They were 0.08–0.50 and 0.16–0.94 for expression-only niches, and 0.02–0.80 and 0.22–0.94 for label-composition niches. The same niche compared across the three models within Lung13 gave 0.98–1.00.

**Supplementary Table S4 (continued).** **e,** Seed reproducibility of niches. Values are the median seed-to-seed ARI over the sweep configurations with four or five niches (24 per representation) and, as a control, with three niches (12). The last column of each block counts the models whose median exceeds that of the expression-only niches. The analyzed sections are shown for reference. **f,** Concept-level recurrence (Methods). J* is the maximum Jaccard between a niche's top-25 concept labels and those of any donor niche, given as the range over donors. Donors counts the donors in which the niche recurs ($P < 0.05$ against random sets of 25 concepts; null mean 0.16–0.18). Models counts the models in which it does so in at least three of four donors. Counts in brackets use a frequency-weighted null$^{\dagger}$, with concepts drawn in proportion to the number of niche lists that contain them (null mean 0.21–0.25). **g,** Immune infiltration pattern at the tumor-niche boundary (descriptive; Methods). For each section and partition, the panel gives the tumor niche, its tumor or tumor-like epithelial share, its share of all cells, and the T-cell density inside and outside the niche. Class follows from the inside/outside ratio. Model rows use each model's sweep-selected configuration, and expression-only rows$^{\dagger}$ use the expression-only sweep winner. Configuration codes as in Supplementary Table S5. Density profiles across the boundary are shown in Figure 7c. Panel g follows on the next page.

**d** Macrophage neighborhood prediction ($R^2$)

| | | T-cell share | | | | Myeloid share | | | |
|---|---|---|---|---|---|---|---|---|---|
| Section | Macrophages | Genes | scGPT | Geneformer | C2S-scale | Genes | scGPT | Geneformer | C2S-scale |
| Lung13 (analyzed) | 17,989 | 0.43 | 0.35 | 0.37 | 0.42 | 0.46 | 0.35 | 0.36 | 0.43 |
| Lung5 | 7,409 | 0.26 | 0.18 | 0.19 | 0.25 | 0.38 | 0.28 | 0.27 | 0.36 |
| Lung6 | 6,884 | 0.14 | 0.13 | 0.12 | 0.14 | 0.47 | 0.42 | 0.43 | 0.46 |
| Lung9 | 4,080 | 0.07 | 0.07 | 0.07 | 0.10 | 0.30 | 0.25 | 0.25 | 0.31 |
| Lung12 (excluded) | 1,291 | 0.12 | 0.07 | 0.05 | 0.15 | 0.05 | 0.01 | 0.00 | 0.04 |

In Lung9 the myeloid-share $R^2$ was 0.248 for scGPT and 0.247 for Geneformer before rounding.

**e** Seed reproducibility of niches (median seed-to-seed ARI)

| | Four to five niches | | | | | Three niches (control) | | | | |
|---|---|---|---|---|---|---|---|---|---|---|
| Section | scGPT | Geneformer | C2S-scale | Expression-only | Models above expression-only | scGPT | Geneformer | C2S-scale | Expression-only | Models above expression-only |
| Lung13 (analyzed) | 0.78 | 0.91 | 0.88 | 0.71 | 3/3 | 0.99 | 0.99 | 1.00 | 1.00 | 0/3 |
| Lung5 | 0.99 | 1.00 | 1.00 | 0.79 | 3/3 | 1.00 | 1.00 | 0.99 | 0.66 | 3/3 |
| Lung6 | 0.98 | 0.85 | 0.99 | 0.50 | 3/3 | 0.82 | 0.77 | 0.76 | 0.69 | 3/3 |
| Lung9 | 0.89 | 0.88 | 0.86 | 0.77 | 3/3 | 1.00 | 1.00 | 1.00 | 0.76 | 3/3 |
| Lung12 | 0.99 | 0.98 | 0.89 | 0.69 | 3/3 | 0.99 | 0.95 | 0.99 | 0.85 | 3/3 |
| Xenium RCC (analyzed) | 0.93 | 0.92 | 0.88 | 0.69 | 3/3 | 1.00 | 1.00 | 1.00 | 0.92 | 3/3 |

**f** Concept-level recurrence of the analyzed section's niches

| Niche | scGPT: J*; donors | Geneformer: J*; donors | C2S-scale: J*; donors | Models | Recurs across donors |
|---|---|---|---|---|---|
| 0 tumor | 0.35–0.43; 4/4 (3/4) | 0.32–0.47; 4/4 (4/4) | 0.28–0.52; 4/4 (2/4) | 3/3 (2/3) | yes |
| 1 macrophage-rich | 0.16–0.32; 2/4 (1/4) | 0.25–0.28; 4/4 (0/4) | 0.16–0.39; 2/4 (1/4) | 1/3 (0/3) | no |
| 2 stromal-lymphoid | 0.52–0.72; 4/4 (4/4) | 0.39–0.56; 4/4 (4/4) | 0.43–0.67; 4/4 (4/4) | 3/3 (3/3) | yes |

The donor niche with the highest J* was also the best composition match in 32 of 36 donor–model–niche cases, although none of these matches reached the composition threshold of panel c.

**Supplementary Table S4 (continued).** Panel g is described on the previous page.

**g** Immune infiltration pattern at the tumor-niche boundary

| Section | Partition | Configuration | Tumor niche | Tumor share | Cells in niche (%) | T-cell density (%) | | Ratio in/out | Class |
|---|---|---|---|---|---|---|---|---|---|
| | | | | | | Inside | Outside | | |
| Lung13 (analyzed) | scGPT | r100/xm/kmeans/k3 | 0 | 0.89 | 34.1 | 3.33 | 26.12 | 0.127 | excluded |
| | Geneformer | r120/m/kmeans/k3 | 0 | 0.85 | 34.4 | 3.92 | 25.91 | 0.151 | excluded |
| | C2S-scale | r100/m/kmeans/k3 | 0 | 0.88 | 33.8 | 3.48 | 25.93 | 0.134 | excluded |
| | Expression-only† | r120/m/gmm/k3 | 0 | 0.88 | 32.9 | 3.66 | 25.52 | 0.144 | excluded |
| Lung5 | scGPT | r100/xm/kmeans/k3 | 0 | 0.87 | 20.6 | 0.12 | 12.63 | 0.009 | desert |
| | Geneformer | r120/xm/kmeans/k3 | 0 | 0.91 | 19.8 | 0.11 | 12.51 | 0.009 | desert |
| | C2S-scale | r120/xm/kmeans/k3 | 0 | 0.92 | 19.7 | 0.09 | 12.50 | 0.007 | desert |
| | Expression-only† | r120/xm/gmm/k3 | 0 | 0.97 | 18.4 | 0.00 | 12.33 | 0.000 | desert |
| Lung6 | scGPT | r100/m/kmeans/k3 | 0 | 0.97 | 74.4 | 0.19 | 11.02 | 0.017 | desert |
| | Geneformer | r120/m/gmm/k4 | 0 | 0.99 | 42.2 | 0.03 | 5.11 | 0.005 | desert |
| | C2S-scale | r100/xm/kmeans/k3 | 0 | 0.98 | 74.4 | 0.13 | 11.21 | 0.012 | desert |
| | Expression-only† | r150/xm/gmm/k3 | 0 | 1.00 | 49.0 | 0.00 | 5.81 | 0.000 | desert |
| Lung9 | scGPT | r150/m/gmm/k3 | 0 | 0.98 | 33.9 | 0.36 | 12.99 | 0.028 | desert |
| | Geneformer | r100/xm/kmeans/k3 | 0 | 0.94 | 45.7 | 0.59 | 15.54 | 0.038 | desert |
| | C2S-scale | r100/xm/kmeans/k3 | 0 | 0.97 | 43.4 | 0.46 | 15.03 | 0.030 | desert |
| | Expression-only† | r100/xm/gmm/k3 | 0 | 0.99 | 42.1 | 0.28 | 14.84 | 0.019 | desert |
| Lung12 | scGPT | r120/xm/kmeans/k3 | 0 | 0.69 | 23.4 | 1.36 | 6.19 | 0.220 | excluded |
| | Geneformer | r100/m/kmeans/k3 | 0 | 0.82 | 19.6 | 0.54 | 6.16 | 0.087 | desert |
| | C2S-scale | r100/m/kmeans/k3 | 0 | 0.87 | 18.8 | 0.28 | 6.16 | 0.045 | desert |
| | Expression-only† | r120/xm/gmm/k3 | 0 | 0.81 | 23.8 | 0.41 | 6.51 | 0.063 | desert |
| Xenium RCC | scGPT | r100/xm/gmm/k5 | 1 | 0.35 | 19.1 | 13.28 | 22.57 | 0.588 | inflamed |
| | Geneformer | r120/m/kmeans/k5 | 1 | 0.26 | 22.4 | 18.20 | 21.54 | 0.845 | inflamed |
| | C2S-scale | r100/xm/kmeans/k5 | 4 | 0.35 | 16.4 | 16.04 | 21.72 | 0.739 | inflamed |
| | Expression-only† | r100/m/gmm/k5 | 3 | 0.24 | 29.1 | 19.03 | 21.51 | 0.885 | inflamed |

Supplementary Table S5: **Niche agreement with cell labels and stability of the selected configurations.** ARI and normalized mutual information (NMI) of each model's sweep-selected niches against the CosMx author labels or the Xenium labels. Label composition (Squidpy) is a calibration baseline built from the evaluation labels. Expression-only niches are defined in the Methods. Configuration gives the neighborhood radius r in coordinate units, the feature space, the clustering method, and the number of niches k. In CosMx, r100, r120, and r150 are 18, 21.6, and 27 µm. In Xenium they are 100, 120, and 150 µm. Seed ARI, purity, null purity (tile shuffle, tiles of 4r), and purity z are those of the selected sweep row. Xenium is shown at three niches, which have the highest niche score overall, and at each model's best-scoring five-niche configuration. Layer in code is the zero-based layer identifier, followed by the layer number.

| Dataset | Partition | Layer in code (layer number) | Configuration | Features | ARI | NMI | Seed ARI | Purity | Null purity | Purity z |
|---|---|---|---|---|---|---|---|---|---|---|
| CosMx | scGPT | layer_1.norm2 (2) | r100/xm/kmeans/k3 | 194 | 0.419 | 0.358 | 0.989 | 0.822 | 0.625 | 378.3 |
| CosMx | Geneformer | layer_1 (2) | r120/m/kmeans/k3 | 261 | 0.342 | 0.292 | 0.998 | 0.832 | 0.608 | 390.6 |
| CosMx | C2S-scale | layer_13 (14) | r100/m/kmeans/k3 | 170 | 0.377 | 0.323 | 0.998 | 0.847 | 0.632 | 338.5 |
| CosMx | Expression-only (sweep winner) | – | r120/m/gmm/k3 | 50 | 0.301 | 0.317 | 1.000 | 0.881 | 0.678 | 307.7 |
| CosMx | Expression-only (scGPT config.) | – | r100/xm/kmeans/k3 | 50 | 0.330 | 0.351 | | | | |
| CosMx | Expression-only (Geneformer config.) | – | r120/m/kmeans/k3 | 50 | 0.268 | 0.269 | | | | |
| CosMx | Expression-only (C2S-scale config.) | – | r100/m/kmeans/k3 | 50 | 0.287 | 0.292 | | | | |
| CosMx | Label composition (Squidpy) | – | k3 | – | 0.269 | 0.289 | | | | |
| *Xenium, marker-based labels, highest-scoring configurations (three niches)* | | | | | | | | | | |
| Xenium | scGPT | layer_2.norm2 (3) | r100/m/gmm/k3 | 118 | 0.055 | 0.120 | 1.000 | 0.972 | 0.911 | 474.0 |
| Xenium | Geneformer | layer_1 (2) | r150/m/gmm/k3 | 125 | 0.059 | 0.123 | 1.000 | 0.969 | 0.903 | 606.9 |
| Xenium | C2S-scale | layer_13 (14) | r150/m/gmm/k3 | 94 | 0.060 | 0.124 | 1.000 | 0.965 | 0.895 | 607.5 |
| Xenium | Expression-only (sweep winner) | – | r100/m/gmm/k3 | 50 | 0.059 | 0.125 | 1.000 | 0.975 | 0.924 | 520.7 |
| Xenium | Expression-only (scGPT config.) | – | r100/m/gmm/k3 | 50 | 0.059 | 0.125 | | | | |
| Xenium | Expression-only (Geneformer config.) | – | r150/m/gmm/k3 | 50 | 0.057 | 0.116 | | | | |
| Xenium | Expression-only (C2S-scale config.) | – | r150/m/gmm/k3 | 50 | 0.057 | 0.116 | | | | |
| Xenium | Label composition (Squidpy) | – | r100/k3 | – | 0.078 | 0.151 | | | | |
| *Xenium, marker-based labels, highest-scoring five-niche configurations (main text)* | | | | | | | | | | |
| Xenium | scGPT | layer_2.norm2 (3) | r100/xm/gmm/k5 | 118 | 0.081 | 0.144 | 0.999 | 0.531 | 0.484 | 389.2 |
| Xenium | Geneformer | layer_1 (2) | r120/m/kmeans/k5 | 125 | 0.077 | 0.141 | 0.999 | 0.876 | 0.681 | 1982.9 |
| Xenium | C2S-scale | layer_13 (14) | r100/xm/kmeans/k5 | 94 | 0.068 | 0.145 | 0.996 | 0.866 | 0.699 | 1004.5 |
| Xenium | Expression-only (sweep winner)† | – | r100/m/gmm/k5 | 50 | 0.075 | 0.143 | 0.966 | 0.925 | 0.782 | 889.8 |
| Xenium | Expression-only (scGPT config.) | – | r100/xm/gmm/k5 | 50 | 0.090 | 0.182 | | | | |
| Xenium | Expression-only (Geneformer config.) | – | r120/m/kmeans/k5 | 50 | 0.052 | 0.105 | | | | |
| Xenium | Expression-only (C2S-scale config.) | – | r100/xm/kmeans/k5 | 50 | 0.034 | 0.095 | | | | |
| Xenium | Label composition (Squidpy) | – | r100/k5 | – | 0.074 | 0.156 | | | | |
| Xenium | Label composition (Squidpy) | – | r120/k5 | – | 0.085 | 0.153 | | | | |
| *Xenium, transferred (TACCO) labels, secondary; three niches* | | | | | | | | | | |
| Xenium | scGPT | layer_2.norm2 (3) | r100/m/gmm/k3 | 118 | 0.022 | 0.054 | | | | |
| Xenium | Geneformer | layer_1 (2) | r150/m/gmm/k3 | 125 | 0.022 | 0.052 | | | | |
| Xenium | C2S-scale | layer_13 (14) | r150/m/gmm/k3 | 94 | 0.025 | 0.054 | | | | |
| Xenium | Label composition (Squidpy) | – | k3 | – | 0.042 | 0.056 | | | | |
| *Xenium, transferred (TACCO) labels, secondary; five niches* | | | | | | | | | | |
| Xenium | scGPT | layer_2.norm2 (3) | r100/xm/gmm/k5 | 118 | 0.026 | 0.061 | | | | |
| Xenium | Geneformer | layer_1 (2) | r120/m/kmeans/k5 | 125 | 0.031 | 0.069 | | | | |
| Xenium | C2S-scale | layer_13 (14) | r100/xm/kmeans/k5 | 94 | 0.034 | 0.069 | | | | |
| Xenium | Label composition (Squidpy) | – | k5 | – | 0.049 | | | | | |

The niche score caps purity z at 10 (Methods). Purity z exceeded this cap in every five-niche configuration (329–1,983), so the ranking among five-niche configurations was set by seed ARI and separability. †Post hoc. For every model, each of the three niches shared by all models has a best match among the niches of the expression-only five-niche winner at residual-profile cosine 0.88–0.99. At six and seven niches (outside the sweep grid; post hoc), mean cross-model ARI was 0.63 and 0.66, against 0.44 at five niches, with five and seven niches shared by all three models. The Geneformer and C2S-scale partitions were less stable across seeds (seed-to-seed ARI 0.84–0.87, against ≥0.996 at five niches).

Supplementary Table S6: **Cross-model niche correspondence in Xenium (marker-based labels). a,** Three niches (Supplementary Figure S4h). Cosine similarity of niche residual profiles between C2S-scale niches (rows) and scGPT niches (columns). The matched niches are n0 (lymphoid border), n1 (myeloid/tumor/plasma), and n2 (tumor/vascular-mural). **b,** Five niches (Figure 4). Residual-profile cosine of the composition-matched niches of each model pair, grouped as in Figure 4, and the ARI between the two partitions. Brackets give the niche numbers in the first and second model. For scGPT–C2S-scale, the composition and cell-overlap matchings disagree for the two model-specific niches (Figure 4h).

**a** Three niches

| | scGPT n0 | scGPT n1 | scGPT n2 |
|---|---|---|---|
| C2S n0 | 0.96 | −0.78 | −0.49 |
| C2S n1 | −0.69 | 1.00 | −0.11 |
| C2S n2 | −0.78 | 0.17 | 0.90 |

**b** Five niches

| Matched group | scGPT–Geneformer | scGPT–C2S-scale | Geneformer–C2S-scale |
|---|---|---|---|
| B/T-lymphoid (shared) | 0.97 (n0/n4) | 0.96 (n0/n0) | 0.91 (n4/n0) |
| Tumor/myeloid (shared) | 0.97 (n1/n1) | 0.97 (n1/n4) | 0.97 (n1/n4) |
| Tumor/vascular (shared) | 0.99 (n3/n2) | 0.99 (n3/n2) | 1.00 (n2/n2) |
| Model-specific (fibroblast/monocyte/LE match) | 0.82 (n2/n0) | 0.18 (n2/n3) | 0.56 (n0/n3) |
| Model-specific (plasma-rich match) | 0.76 (n4/n3) | 0.42 (n4/n1) | 0.88 (n3/n1) |
| Partition ARI | 0.33 | 0.38 | 0.60 |

Supplementary Table S7: **Marker support of the transferred (TACCO) Xenium labels.** Own marker detected is the percentage of cells with at least one transcript of any of the label's own panel markers. PTPRC and KRT8 detection are shown for comparison. Labels without a marker on the panel cannot be checked (n/a). Transferred labels name the most similar class of a tumor-naive kidney reference (GSE183273). On this RCC section they indicate similarity to that class, so PT is written PT-like in Supplementary Figure S13c and Supplementary Table S16. EC, endothelial; IMM, immune; VSM/P, vascular smooth muscle/pericyte; FIB, fibroblast; TAL, thick ascending limb; PC, principal cell; IC, intercalated cell; PT, proximal tubule; POD, podocyte; DTL, descending thin limb; CNT, connecting tubule; DCT, distal convoluted tubule; ATL, ascending thin limb; PEC, parietal epithelial cell.

| Transferred label | Cells | % of section | Own markers on panel | Own marker detected (%) | PTPRC detected (%) | KRT8 detected (%) |
|---|---|---|---|---|---|---|
| EC | 68340 | 14.7 | PECAM1, VWF, CD34 | 85.0 | 24.7 | 23.0 |
| IMM | 106349 | 22.9 | PTPRC, CD3E, CD68 | 94.7 | 84.2 | 18.1 |
| VSM/P | 23874 | 5.1 | ACTA2, MYH11, PDGFRB | 89.7 | 14.9 | 28.5 |
| FIB | 63602 | 13.7 | PDGFRA | 21.3 | 27.8 | 38.3 |
| TAL | 65248 | 14.0 | UMOD | 0.2 | 55.5 | 21.1 |
| PC | 32881 | 7.1 | AQP2, AQP3 | 1.1 | 50.5 | 47.2 |
| IC | 32901 | 7.1 | SLC4A1, FOXI1 | 1.8 | 79.2 | 23.5 |
| PT | 31133 | 6.7 | none | n/a | 61.3 | 65.0 |
| POD | 18427 | 4.0 | none | n/a | 47.6 | 31.6 |
| DTL | 7412 | 1.6 | none | n/a | 43.4 | 91.6 |
| CNT | 5804 | 1.2 | none | n/a | 73.4 | 21.9 |
| DCT | 4912 | 1.1 | none | n/a | 66.0 | 14.9 |
| ATL | 2272 | 0.5 | none | n/a | 21.7 | 9.4 |
| PEC | 2065 | 0.4 | none | n/a | 17.4 | 70.2 |

Supplementary Table S8: **Calibrated interface calls.** Every niche-by-cell-pair test with $|z| \geq z^*$ is listed, for CosMx (author labels, three niches) and Xenium (marker-based labels, five niches per model). z is the calibrated interface statistic (Methods). Edges are cell pairs of the two types within 120 units (21.6 µm in CosMx, 120 µm in Xenium). Effect is a percentage of the mean unadjusted edge program load. LR rows are the matched-niche CellChat rows of the pair, all of which receive the pair's call. Xenium niches carry their model-specific number. L, M, and V mark the B/T-lymphoid, tumor/myeloid, and tumor/vascular niches shared by all three models (Figure 4). The tables of all tests are deposited with the processed data (Zenodo; source tables table_S8_all_tests_cosmx_and_xenium_k3.csv and table_S8_all_tests_xenium_marker_k5.csv). Notes on the CosMx calls follow the table.

| Dataset | Labels | Model | Niche | Cell pair | Edges | z | Effect (%) | $z^*$ | Call | LR rows | Term-supp. LR rows |
|---|---|---|---|---|---|---|---|---|---|---|---|
| CosMx | author | scGPT | 2 | fibroblast – t cd8 naive | 487 | 2.65 | 0.50 | 2.00 | enriched | 5 | 5 |
| CosMx | author | scGPT | 2 | t cd8 memory – b plasma | 178 | -2.89 | -0.55 | 2.00 | depleted | 8 | 8 |
| CosMx | author | scGPT | 1 | macrophage – t cd4 memory | 8606 | -2.93 | -0.17 | 2.00 | depleted | 11 | 3 |
| CosMx | author | scGPT | 1 | t cd4 memory – t cd8 memory | 2887 | -2.02 | -0.22 | 2.00 | depleted | 14 | 6 |
| CosMx | author | scGPT | 1 | t cd4 memory – treg | 1352 | -2.14 | -0.34 | 2.00 | depleted | 8 | 1 |
| CosMx | author | scGPT | 1 | t cd4 memory – dendritic | 1384 | -2.64 | -0.33 | 2.00 | depleted | 8 | 1 |
| CosMx | author | scGPT | 1 | t cd8 memory – t cd8 naive | 53 | 2.24 | 1.48 | 2.00 | enriched | 10 | 9 |
| CosMx | author | Geneformer | 0 | macrophage – t cd8 memory | 372 | 3.14 | 0.30 | 2.00 | enriched | 16 | 4 |
| CosMx | author | Geneformer | 2 | tumor – t cd8 memory | 178 | -2.74 | -0.64 | 2.00 | depleted | 12 | 12 |
| CosMx | author | Geneformer | 2 | fibroblast – t cd4 memory | 21055 | -2.27 | -0.08 | 2.00 | depleted | 0 | 0 |
| CosMx | author | Geneformer | 2 | t cd4 memory – dendritic | 4579 | -2.16 | -0.15 | 2.00 | depleted | 0 | 0 |
| CosMx | author | Geneformer | 2 | t cd8 memory – t cd8 naive | 56 | 2.17 | 0.45 | 2.00 | enriched | 8 | 8 |
| CosMx | author | Geneformer | 2 | b plasma – mast | 699 | -2.19 | -0.28 | 2.00 | depleted | 7 | 7 |
| CosMx | author | Geneformer | 1 | tumor – treg | 169 | -2.34 | -0.35 | 2.00 | depleted | 12 | 12 |
| CosMx | author | Geneformer | 1 | fibroblast – t cd4 memory | 2744 | 2.04 | 0.11 | 2.00 | enriched | 17 | 17 |
| CosMx | author | Geneformer | 1 | macrophage – b plasma | 487 | -2.01 | -0.32 | 2.00 | depleted | 3 | 3 |
| CosMx | author | C2S-scale | 0 | tumor – fibroblast | 1922 | 2.58 | 0.20 | 2.05 | enriched | 27 | 27 |
| CosMx | author | C2S-scale | 2 | tumor – mast | 116 | -2.14 | -0.52 | 2.05 | depleted | 15 | 14 |
| CosMx | author | C2S-scale | 2 | macrophage – treg | 1326 | -2.90 | -0.23 | 2.05 | depleted | 9 | 8 |
| CosMx | author | C2S-scale | 1 | tumor – treg | 167 | -2.54 | -0.54 | 2.05 | depleted | 11 | 11 |
| CosMx | author | C2S-scale | 1 | t cd4 memory – t cd8 naive | 76 | -2.28 | -0.86 | 2.05 | depleted | 12 | 12 |
| Xenium | marker | scGPT | 0 (L) | monocyte – B cell | 412415 | 7.57 | 0.48 | 2.10 | enriched | 3 | 3 |
| Xenium | marker | scGPT | 0 (L) | lymphatic endothelial – B cell | 212600 | 4.09 | 0.31 | 2.10 | enriched | 0 | 0 |
| Xenium | marker | scGPT | 0 (L) | fibroblast – B cell | 310667 | 3.34 | 0.22 | 2.10 | enriched | 1 | 1 |
| Xenium | marker | scGPT | 0 (L) | T cell – fibroblast | 720790 | 3.22 | 0.08 | 2.10 | enriched | 3 | 3 |
| Xenium | marker | scGPT | 0 (L) | NK – mast | 69 | 2.24 | 1.37 | 2.10 | enriched | 1 | 1 |
| Xenium | marker | scGPT | 1 (M) | tumor-like epithelial – myeloid | 1406179 | 3.42 | 0.03 | 2.10 | enriched | 4 | 4 |
| Xenium | marker | scGPT | 1 (M) | endothelial – lymphatic endothelial | 9000 | 2.32 | 0.15 | 2.10 | enriched | 3 | 1 |
| Xenium | marker | scGPT | 1 (M) | monocyte – B cell | 876 | 2.31 | 0.46 | 2.10 | enriched | 5 | 5 |

Continued on next page

Supplementary Table S8 (continued)

| Dataset | Labels | Model | Niche | Cell pair | Edges | z | Effect (%) | $z^*$ | Call | LR rows | Term-supp. LR rows |
|---|---|---|---|---|---|---|---|---|---|---|---|
| Xenium | marker | scGPT | 1 (M) | fibroblast – monocyte | 4421 | -2.29 | -0.20 | 2.10 | depleted | 7 | 7 |
| Xenium | marker | scGPT | 1 (M) | mast – plasma | 621 | -2.37 | -0.37 | 2.10 | depleted | 0 | 0 |
| Xenium | marker | scGPT | 1 (M) | myeloid – monocyte | 72895 | -2.41 | -0.07 | 2.10 | depleted | 7 | 7 |
| Xenium | marker | scGPT | 1 (M) | NK – tumor-like epithelial | 22513 | -2.41 | -0.09 | 2.10 | depleted | 1 | 1 |
| Xenium | marker | scGPT | 1 (M) | monocyte – lymphatic endothelial | 3510 | -2.49 | -0.21 | 2.10 | depleted | 10 | 8 |
| Xenium | marker | scGPT | 1 (M) | T cell – plasma | 105796 | -2.54 | -0.11 | 2.10 | depleted | 2 | 0 |
| Xenium | marker | scGPT | 1 (M) | fibroblast – lymphatic endothelial | 6896 | -2.58 | -0.16 | 2.10 | depleted | 3 | 1 |
| Xenium | marker | scGPT | 1 (M) | fibroblast – dendritic | 2705 | -2.72 | -0.24 | 2.10 | depleted | 6 | 6 |
| Xenium | marker | scGPT | 1 (M) | myeloid – endothelial | 125690 | -3.18 | -0.08 | 2.10 | depleted | 6 | 4 |
| Xenium | marker | scGPT | 1 (M) | myeloid – mural | 86924 | -3.20 | -0.07 | 2.10 | depleted | 5 | 5 |
| Xenium | marker | scGPT | 1 (M) | myeloid – plasma | 198449 | -5.22 | -0.16 | 2.10 | depleted | 8 | 6 |
| Xenium | marker | scGPT | 3 (V) | T cell – plasma | 936 | 2.50 | 0.40 | 2.10 | enriched | 2 | 0 |
| Xenium | marker | scGPT | 3 (V) | endothelial – lymphatic endothelial | 33761 | 2.33 | 0.09 | 2.10 | enriched | 2 | 0 |
| Xenium | marker | scGPT | 3 (V) | NK – mural | 26375 | -2.19 | -0.09 | 2.10 | depleted | 0 | 0 |
| Xenium | marker | scGPT | 2 | monocyte – B cell | 68839 | 4.05 | 0.31 | 2.10 | enriched | 3 | 1 |
| Xenium | marker | scGPT | 2 | mural – B cell | 17401 | 2.48 | 0.25 | 2.10 | enriched | 1 | 0 |
| Xenium | marker | scGPT | 2 | tumor-like epithelial – fibroblast | 18112 | -2.10 | -0.17 | 2.10 | depleted | 0 | 0 |
| Xenium | marker | scGPT | 2 | T cell – myeloid | 60778 | -2.20 | -0.09 | 2.10 | depleted | 2 | 0 |
| Xenium | marker | scGPT | 2 | NK – myeloid | 4282 | -2.45 | -0.22 | 2.10 | depleted | 2 | 0 |
| Xenium | marker | scGPT | 2 | NK – plasma | 10130 | -2.46 | -0.20 | 2.10 | depleted | 1 | 1 |
| Xenium | marker | scGPT | 2 | T cell – plasma | 126552 | -2.58 | -0.09 | 2.10 | depleted | 1 | 1 |
| Xenium | marker | scGPT | 2 | NK – tumor-like epithelial | 1927 | -2.62 | -0.46 | 2.10 | depleted | 0 | 0 |
| Xenium | marker | scGPT | 2 | monocyte – plasma | 55047 | -3.25 | -0.13 | 2.10 | depleted | 6 | 4 |
| Xenium | marker | scGPT | 4 | tumor-like epithelial – mural | 36711 | 2.73 | 0.12 | 2.10 | enriched | 0 | 0 |
| Xenium | marker | scGPT | 4 | tumor-like epithelial – plasma | 33613 | 2.72 | 0.19 | 2.10 | enriched | 0 | 0 |
| Xenium | marker | scGPT | 4 | mast – plasma | 81 | 2.33 | 1.07 | 2.10 | enriched | 0 | 0 |
| Xenium | marker | scGPT | 4 | myeloid – fibroblast | 347199 | -2.37 | -0.05 | 2.10 | depleted | 2 | 2 |
| Xenium | marker | scGPT | 4 | T cell – myeloid | 985223 | -2.39 | -0.04 | 2.10 | depleted | 2 | 2 |
| Xenium | marker | scGPT | 4 | mural – mast | 93 | -2.40 | -0.72 | 2.10 | depleted | 0 | 0 |
| Xenium | marker | scGPT | 4 | fibroblast – monocyte | 125129 | -2.81 | -0.10 | 2.10 | depleted | 3 | 3 |
| Xenium | marker | scGPT | 4 | T cell – plasma | 248863 | -2.89 | -0.09 | 2.10 | depleted | 0 | 0 |
| Xenium | marker | scGPT | 4 | myeloid – endothelial | 213230 | -3.45 | -0.08 | 2.10 | depleted | 2 | 2 |
| Xenium | marker | scGPT | 4 | myeloid – plasma | 166904 | -4.05 | -0.13 | 2.10 | depleted | 2 | 2 |
| Xenium | marker | scGPT | 4 | myeloid – dendritic | 35496 | -4.10 | -0.16 | 2.10 | depleted | 6 | 6 |
| Xenium | marker | Geneformer | 4 (L) | monocyte – B cell | 755903 | 3.96 | 0.26 | 2.06 | enriched | 3 | 3 |
| Xenium | marker | Geneformer | 4 (L) | lymphatic endothelial – B cell | 375418 | 3.14 | 0.23 | 2.06 | enriched | 1 | 1 |
| Xenium | marker | Geneformer | 4 (L) | endothelial – plasma | 77634 | -2.09 | -0.13 | 2.06 | depleted | 3 | 1 |
| Xenium | marker | Geneformer | 4 (L) | T cell – fibroblast | 990730 | -2.14 | -0.08 | 2.06 | depleted | 1 | 1 |
| Xenium | marker | Geneformer | 4 (L) | NK – fibroblast | 42375 | -2.28 | -0.17 | 2.06 | depleted | 0 | 0 |
| Xenium | marker | Geneformer | 4 (L) | dendritic – plasma | 48431 | -2.36 | -0.21 | 2.06 | depleted | 1 | 1 |
| Xenium | marker | Geneformer | 4 (L) | fibroblast – monocyte | 235873 | -2.36 | -0.09 | 2.06 | depleted | 3 | 3 |
| Xenium | marker | Geneformer | 4 (L) | myeloid – dendritic | 19419 | -2.45 | -0.23 | 2.06 | depleted | 6 | 6 |

Continued on next page

Supplementary Table S8 (continued)

| Dataset | Labels | Model | Niche | Cell pair | Edges | z | Effect (%) | $z^*$ | Call | LR rows | Term-supp. LR rows |
|---|---|---|---|---|---|---|---|---|---|---|---|
| Xenium | marker | Geneformer | 4 (L) | monocyte – dendritic | 105067 | -2.60 | -0.14 | 2.06 | depleted | 7 | 7 |
| Xenium | marker | Geneformer | 4 (L) | myeloid – monocyte | 46525 | -2.62 | -0.17 | 2.06 | depleted | 7 | 7 |
| Xenium | marker | Geneformer | 4 (L) | myeloid – fibroblast | 35035 | -2.87 | -0.19 | 2.06 | depleted | 2 | 2 |
| Xenium | marker | Geneformer | 4 (L) | monocyte – plasma | 132700 | -2.93 | -0.17 | 2.06 | depleted | 6 | 4 |
| Xenium | marker | Geneformer | 4 (L) | NK – monocyte | 58968 | -3.17 | -0.19 | 2.06 | depleted | 3 | 3 |
| Xenium | marker | Geneformer | 4 (L) | T cell – B cell | 2861061 | -3.22 | -0.17 | 2.06 | depleted | 2 | 2 |
| Xenium | marker | Geneformer | 4 (L) | T cell – dendritic | 572959 | -3.54 | -0.15 | 2.06 | depleted | 3 | 3 |
| Xenium | marker | Geneformer | 4 (L) | T cell – monocyte | 1412892 | -5.05 | -0.15 | 2.06 | depleted | 6 | 6 |
| Xenium | marker | Geneformer | 1 (M) | tumor-like epithelial – myeloid | 1928426 | 4.92 | 0.05 | 2.06 | enriched | 5 | 5 |
| Xenium | marker | Geneformer | 1 (M) | T cell – monocyte | 54348 | 2.36 | 0.15 | 2.06 | enriched | 5 | 4 |
| Xenium | marker | Geneformer | 1 (M) | monocyte – dendritic | 3316 | 2.20 | 0.25 | 2.06 | enriched | 7 | 6 |
| Xenium | marker | Geneformer | 1 (M) | endothelial – B cell | 1228 | -2.19 | -0.28 | 2.06 | depleted | 1 | 1 |
| Xenium | marker | Geneformer | 1 (M) | monocyte – lymphatic endothelial | 6985 | -2.27 | -0.25 | 2.06 | depleted | 10 | 6 |
| Xenium | marker | Geneformer | 1 (M) | myeloid – endothelial | 254000 | -2.46 | -0.07 | 2.06 | depleted | 4 | 2 |
| Xenium | marker | Geneformer | 1 (M) | myeloid – mural | 259459 | -2.47 | -0.07 | 2.06 | depleted | 5 | 5 |
| Xenium | marker | Geneformer | 1 (M) | myeloid – plasma | 237637 | -2.85 | -0.11 | 2.06 | depleted | 4 | 2 |
| Xenium | marker | Geneformer | 2 (V) | endothelial – lymphatic endothelial | 80859 | 2.61 | 0.12 | 2.06 | enriched | 2 | 0 |
| Xenium | marker | Geneformer | 2 (V) | fibroblast – plasma | 3051 | -2.40 | -0.24 | 2.06 | depleted | 2 | 2 |
| Xenium | marker | Geneformer | 0 | mural – mast | 846 | 2.31 | 0.34 | 2.06 | enriched | 0 | 0 |
| Xenium | marker | Geneformer | 0 | endothelial – monocyte | 230512 | 2.19 | 0.08 | 2.06 | enriched | 2 | 0 |
| Xenium | marker | Geneformer | 0 | endothelial – plasma | 77329 | 2.14 | 0.13 | 2.06 | enriched | 3 | 3 |
| Xenium | marker | Geneformer | 0 | T cell – tumor-like epithelial | 13939 | 2.13 | 0.37 | 2.06 | enriched | 1 | 1 |
| Xenium | marker | Geneformer | 0 | NK – mast | 261 | -2.39 | -0.72 | 2.06 | depleted | 1 | 1 |
| Xenium | marker | Geneformer | 0 | dendritic – mast | 269 | -2.77 | -0.95 | 2.06 | depleted | 2 | 0 |
| Xenium | marker | Geneformer | 0 | myeloid – dendritic | 4105 | -3.10 | -0.56 | 2.06 | depleted | 6 | 0 |
| Xenium | marker | Geneformer | 3 | NK – mast | 555 | 2.69 | 0.63 | 2.06 | enriched | 0 | 0 |
| Xenium | marker | Geneformer | 3 | T cell – fibroblast | 737222 | -2.08 | -0.06 | 2.06 | depleted | 0 | 0 |
| Xenium | marker | Geneformer | 3 | tumor-like epithelial – dendritic | 28322 | -2.22 | -0.16 | 2.06 | depleted | 4 | 4 |
| Xenium | marker | Geneformer | 3 | T cell – lymphatic endothelial | 205780 | -2.37 | -0.10 | 2.06 | depleted | 1 | 1 |
| Xenium | marker | Geneformer | 3 | endothelial – monocyte | 110572 | -2.45 | -0.10 | 2.06 | depleted | 5 | 5 |
| Xenium | marker | Geneformer | 3 | myeloid – dendritic | 106150 | -2.63 | -0.13 | 2.06 | depleted | 6 | 6 |
| Xenium | marker | Geneformer | 3 | tumor-like epithelial – plasma | 179912 | -2.79 | -0.14 | 2.06 | depleted | 0 | 0 |
| Xenium | marker | Geneformer | 3 | mural – plasma | 115461 | -2.82 | -0.16 | 2.06 | depleted | 2 | 2 |
| Xenium | marker | Geneformer | 3 | endothelial – plasma | 155600 | -2.87 | -0.14 | 2.06 | depleted | 2 | 2 |
| Xenium | marker | Geneformer | 3 | monocyte – plasma | 109314 | -2.92 | -0.16 | 2.06 | depleted | 5 | 5 |
| Xenium | marker | Geneformer | 3 | myeloid – B cell | 34717 | -3.15 | -0.25 | 2.06 | depleted | 2 | 2 |
| Xenium | marker | Geneformer | 3 | T cell – dendritic | 170156 | -3.16 | -0.13 | 2.06 | depleted | 2 | 2 |
| Xenium | marker | Geneformer | 3 | NK – myeloid | 118619 | -3.31 | -0.14 | 2.06 | depleted | 2 | 2 |
| Xenium | marker | Geneformer | 3 | T cell – monocyte | 502968 | -3.49 | -0.12 | 2.06 | depleted | 3 | 3 |
| Xenium | marker | Geneformer | 3 | myeloid – mural | 293301 | -3.58 | -0.10 | 2.06 | depleted | 2 | 2 |
| Xenium | marker | Geneformer | 3 | myeloid – fibroblast | 492454 | -3.68 | -0.09 | 2.06 | depleted | 2 | 2 |
| Xenium | marker | Geneformer | 3 | fibroblast – monocyte | 140001 | -3.79 | -0.15 | 2.06 | depleted | 3 | 3 |

Continued on next page

Supplementary Table S8 (continued)

| Dataset | Labels | Model | Niche | Cell pair | Edges | z | Effect (%) | $z^*$ | Call | LR rows | Term-supp. LR rows |
|---|---|---|---|---|---|---|---|---|---|---|---|
| Xenium | marker | Geneformer | 3 | myeloid – endothelial | 419036 | -4.49 | -0.10 | 2.06 | depleted | 4 | 4 |
| Xenium | marker | Geneformer | 3 | myeloid – plasma | 554426 | -5.55 | -0.18 | 2.06 | depleted | 4 | 4 |
| Xenium | marker | Geneformer | 3 | T cell – plasma | 680492 | -5.78 | -0.21 | 2.06 | depleted | 0 | 0 |
| Xenium | marker | Geneformer | 3 | myeloid – monocyte | 294028 | -6.16 | -0.20 | 2.06 | depleted | 7 | 7 |
| Xenium | marker | Geneformer | 3 | T cell – myeloid | 1769080 | -6.84 | -0.13 | 2.06 | depleted | 3 | 3 |
| Xenium | marker | C2S-scale | 0 (L) | monocyte – B cell | 836332 | 3.80 | 0.23 | 2.05 | enriched | 3 | 1 |
| Xenium | marker | C2S-scale | 0 (L) | T cell – fibroblast | 1708722 | 3.68 | 0.10 | 2.05 | enriched | 3 | 1 |
| Xenium | marker | C2S-scale | 0 (L) | fibroblast – dendritic | 122765 | 3.10 | 0.15 | 2.05 | enriched | 3 | 1 |
| Xenium | marker | C2S-scale | 0 (L) | fibroblast – plasma | 295026 | 2.97 | 0.10 | 2.05 | enriched | 2 | 1 |
| Xenium | marker | C2S-scale | 0 (L) | mast – lymphatic endothelial | 3666 | 2.43 | 0.35 | 2.05 | enriched | 1 | 1 |
| Xenium | marker | C2S-scale | 0 (L) | fibroblast – monocyte | 694656 | 2.42 | 0.05 | 2.05 | enriched | 3 | 1 |
| Xenium | marker | C2S-scale | 0 (L) | endothelial – fibroblast | 500711 | 2.38 | 0.05 | 2.05 | enriched | 0 | 0 |
| Xenium | marker | C2S-scale | 0 (L) | lymphatic endothelial – B cell | 447236 | 2.15 | 0.17 | 2.05 | enriched | 1 | 1 |
| Xenium | marker | C2S-scale | 0 (L) | myeloid – dendritic | 22241 | -2.42 | -0.22 | 2.05 | depleted | 6 | 0 |
| Xenium | marker | C2S-scale | 0 (L) | T cell – dendritic | 638570 | -3.45 | -0.16 | 2.05 | depleted | 5 | 0 |
| Xenium | marker | C2S-scale | 4 (M) | endothelial – lymphatic endothelial | 7716 | 2.43 | 0.22 | 2.05 | enriched | 2 | 0 |
| Xenium | marker | C2S-scale | 4 (M) | mural – lymphatic endothelial | 9813 | 2.26 | 0.22 | 2.05 | enriched | 2 | 0 |
| Xenium | marker | C2S-scale | 4 (M) | T cell – plasma | 32167 | 2.10 | 0.15 | 2.05 | enriched | 0 | 0 |
| Xenium | marker | C2S-scale | 4 (M) | dendritic – B cell | 190 | -2.09 | -0.75 | 2.05 | depleted | 2 | 0 |
| Xenium | marker | C2S-scale | 4 (M) | monocyte – plasma | 1319 | -2.10 | -0.33 | 2.05 | depleted | 3 | 0 |
| Xenium | marker | C2S-scale | 4 (M) | myeloid – mast | 616 | -2.77 | -0.55 | 2.05 | depleted | 1 | 0 |
| Xenium | marker | C2S-scale | 2 (V) | tumor-like epithelial – mural | 1050163 | 2.39 | 0.04 | 2.05 | enriched | 1 | 1 |
| Xenium | marker | C2S-scale | 2 (V) | endothelial – B cell | 402 | 2.34 | 0.45 | 2.05 | enriched | 1 | 1 |
| Xenium | marker | C2S-scale | 2 (V) | myeloid – lymphatic endothelial | 159916 | -2.39 | -0.07 | 2.05 | depleted | 6 | 5 |
| Xenium | marker | C2S-scale | 2 (V) | endothelial – mural | 618510 | -2.58 | -0.05 | 2.05 | depleted | 0 | 0 |
| Xenium | marker | C2S-scale | 3 | T cell – tumor-like epithelial | 168786 | 2.55 | 0.14 | 2.05 | enriched | 0 | 0 |
| Xenium | marker | C2S-scale | 3 | fibroblast – lymphatic endothelial | 56951 | 2.17 | 0.14 | 2.05 | enriched | 1 | 1 |
| Xenium | marker | C2S-scale | 3 | T cell – monocyte | 356411 | -2.05 | -0.09 | 2.05 | depleted | 3 | 3 |
| Xenium | marker | C2S-scale | 3 | endothelial – dendritic | 15471 | -2.13 | -0.18 | 2.05 | depleted | 0 | 0 |
| Xenium | marker | C2S-scale | 3 | myeloid – plasma | 8539 | -2.27 | -0.27 | 2.05 | depleted | 4 | 4 |
| Xenium | marker | C2S-scale | 3 | myeloid – monocyte | 168798 | -2.44 | -0.12 | 2.05 | depleted | 7 | 7 |
| Xenium | marker | C2S-scale | 1 | T cell – tumor-like epithelial | 413045 | 2.13 | 0.06 | 2.05 | enriched | 2 | 2 |
| Xenium | marker | C2S-scale | 1 | tumor-like epithelial – mast | 1693 | 2.10 | 0.28 | 2.05 | enriched | 0 | 0 |
| Xenium | marker | C2S-scale | 1 | tumor-like epithelial – lymphatic endothelial | 38789 | 2.07 | 0.10 | 2.05 | enriched | 3 | 3 |
| Xenium | marker | C2S-scale | 1 | T cell – endothelial | 305032 | -2.18 | -0.08 | 2.05 | depleted | 0 | 0 |
| Xenium | marker | C2S-scale | 1 | T cell – myeloid | 1441513 | -2.39 | -0.04 | 2.05 | depleted | 7 | 7 |
| Xenium | marker | C2S-scale | 1 | endothelial – plasma | 193135 | -2.44 | -0.10 | 2.05 | depleted | 2 | 2 |
| Xenium | marker | C2S-scale | 1 | NK – myeloid | 106383 | -2.49 | -0.12 | 2.05 | depleted | 2 | 2 |
| Xenium | marker | C2S-scale | 1 | mural – plasma | 146910 | -2.54 | -0.14 | 2.05 | depleted | 3 | 3 |
| Xenium | marker | C2S-scale | 1 | myeloid – plasma | 771817 | -2.66 | -0.08 | 2.05 | depleted | 4 | 4 |
| Xenium | marker | C2S-scale | 1 | endothelial – monocyte | 49193 | -2.77 | -0.15 | 2.05 | depleted | 6 | 6 |

Continued on next page

Supplementary Table S8 (continued)

| Dataset | Labels | Model | Niche | Cell pair | Edges | z | Effect (%) | $z^*$ | Call | LR rows | Term-supp. LR rows |
|---|---|---|---|---|---|---|---|---|---|---|---|
| Xenium | marker | C2S-scale | 1 | T cell – fibroblast | 320322 | -3.14 | -0.09 | 2.05 | depleted | 0 | 0 |
| Xenium | marker | C2S-scale | 1 | T cell – dendritic | 146348 | -3.62 | -0.16 | 2.05 | depleted | 4 | 4 |
| Xenium | marker | C2S-scale | 1 | myeloid – fibroblast | 302274 | -3.65 | -0.10 | 2.05 | depleted | 6 | 6 |
| Xenium | marker | C2S-scale | 1 | T cell – plasma | 828156 | -4.10 | -0.13 | 2.05 | depleted | 0 | 0 |
| Xenium | marker | C2S-scale | 1 | myeloid – endothelial | 281896 | -4.31 | -0.13 | 2.05 | depleted | 5 | 5 |
| Xenium | marker | C2S-scale | 1 | myeloid – monocyte | 195575 | -4.50 | -0.15 | 2.05 | depleted | 7 | 7 |

In CosMx, between-model Spearman correlations of the niche-free $z$ were 0.15–0.47. For the 13 CosMx cell pairs called in at least one model, the niche-free $z$ did not change from the contacts with the fewest to those with the most partner-marker transcripts (median tercile gradient 0.07, 95% CI $-0.77$ to 0.58). The CosMx calls therefore do not show the spillover pattern [1, 2], although they are weak (tercile $z$ within $\pm 3.5$).

Supplementary Table S9: **Calibration of the interface statistic and call counts.** $|z| > 2$ is the share of niche-by-pair tests with $|z| > 2$ under a conventional, uncalibrated 400-unit tile null and under the calibrated 120-unit block null (Methods). In CosMx these are 72 and 21.6 µm, and in Xenium 400 and 120 µm. Median and maximum |effect| are over all tests. The smooth-field false-positive rate (FPR) is the share of pairs with $|z| > 2$ when contact-free smooth random fields, matched to the autocorrelation of the score, are passed through each null (5% nominal). Noise FPR is the same for independent noise under the calibrated block null. Xenium is shown at three niches and at the five niches reported in the main text.

| | | | | | $\|z\| > 2$ (%) | | | | | | Smooth-field FPR (%) | | |
|---|---|---|---|---|---|---|---|---|---|---|---|---|---|
| Dataset | Labels | Niches | Model | Tests | Tile null | Calibrated block null | Enriched / depleted | $z^*$ | Median \|effect\| (%) | Max \|effect\| (%) | Tile null | Calibrated block null | Noise FPR, calibrated (%) |
| CosMx | author | 3 | scGPT | 129 | 17.8 | 5.4 | 2 / 5 | 2.00 | 0.13 | 1.48 | 6.3 | 4.8 | 6.2 |
| CosMx | author | 3 | Geneformer | 132 | 18.9 | 6.8 | 3 / 6 | 2.00 | 0.10 | 1.01 | 5.7 | 4.9 | 4.5 |
| CosMx | author | 3 | C2S-scale | 125 | 20.8 | 4.0 | 1 / 4 | 2.05 | 0.10 | 0.93 | 6.8 | 5.9 | 2.4 |
| Xenium | transferred | 3 | scGPT | 270 | 47.8 | 18.5 | 18 / 29 | 2.06 | 0.04 | 1.02 | 12.4 | 6.0 | 5.2 |
| Xenium | transferred | 3 | Geneformer | 271 | 59.4 | 24.7 | 7 / 53 | 2.19 | 0.07 | 0.91 | 15.9 | 6.6 | 5.5 |
| Xenium | transferred | 3 | C2S-scale | 271 | 51.3 | 20.7 | 10 / 46 | 2.02 | 0.06 | 1.07 | 13.2 | 5.4 | 4.4 |
| Xenium | marker | 3 | scGPT | 233 | 30.5 | 15.5 | 9 / 25 | 2.04 | 0.05 | 0.43 | 9.9 | 5.6 | 6.4 |
| Xenium | marker | 3 | Geneformer | 231 | 43.7 | 19.9 | 3 / 40 | 2.07 | 0.09 | 0.56 | 13.5 | 5.8 | 6.1 |
| Xenium | marker | 3 | C2S-scale | 230 | 37.0 | 19.1 | 9 / 33 | 2.05 | 0.06 | 0.63 | 12.0 | 5.3 | 4.8 |
| Xenium | transferred | 5 | scGPT | 444 | 38.3 | 13.7 | 30 / 25 | 2.05 | 0.06 | 1.67 | 9.8 | 5.6 | 4.1 |
| Xenium | transferred | 5 | Geneformer | 446 | 40.8 | 18.4 | 8 / 66 | 2.12 | 0.08 | 1.87 | 15.0 | 6.6 | 5.6 |
| Xenium | transferred | 5 | C2S-scale | 448 | 27.9 | 12.1 | 12 / 38 | 2.08 | 0.06 | 1.13 | 13.0 | 6.1 | 4.9 |
| Xenium | marker | 5 | scGPT | 376 | 37.2 | 12.2 | 15 / 27 | 2.10 | 0.05 | 1.37 | 8.2 | 5.9 | 7.2 |
| Xenium | marker | 5 | Geneformer | 382 | 33.5 | 15.2 | 11 / 44 | 2.06 | 0.07 | 0.95 | 11.1 | 5.7 | 5.8 |
| Xenium | marker | 5 | C2S-scale | 376 | 31.1 | 12.0 | 18 / 24 | 2.05 | 0.06 | 0.86 | 12.2 | 5.9 | 4.5 |

Supplementary Table S10: **Matched-niche CellChat LR rows by the calibrated call of their cell pair.** Counts of CellChat LR rows, with term-supported rows in brackets. Untested rows are homotypic or belong to pairs with fewer than 50 edges in that niche. Xenium uses marker-based labels and is shown at three niches and at the five niches reported in the main text.

| Dataset | Model | All LR rows | On enriched pairs | On depleted pairs | Not called | Untested |
|---|---|---|---|---|---|---|
| CosMx | scGPT | 2016 (1670) | 15 (14) | 49 (19) | 1006 (828) | 946 (809) |
| CosMx | Geneformer | 1817 (1389) | 41 (29) | 34 (34) | 1002 (826) | 740 (500) |
| CosMx | C2S-scale | 2051 (1928) | 27 (27) | 47 (45) | 1042 (1008) | 935 (848) |
| Xenium, three niches | scGPT | 565 (508) | 17 (17) | 95 (83) | 402 (364) | 51 (44) |
| Xenium, three niches | Geneformer | 497 (408) | 9 (9) | 128 (106) | 312 (257) | 48 (36) |
| Xenium, three niches | C2S-scale | 513 (343) | 17 (11) | 104 (39) | 340 (253) | 52 (40) |
| Xenium, five niches | scGPT | 879 (628) | 28 (19) | 84 (68) | 680 (479) | 87 (62) |
| Xenium, five niches | Geneformer | 838 (599) | 29 (23) | 144 (124) | 579 (396) | 86 (56) |
| Xenium, five niches | C2S-scale | 847 (535) | 28 (15) | 83 (65) | 644 (400) | 92 (55) |

The scGPT macrophage–tumor pair in the CosMx tumor niche had 36 CellChat rows (28 term-supported) without enriched program burden ($z = +0.53$).

Supplementary Table S11: **Interface calls after removal of marker-detecting features (exploratory; Supplementary Table S19, C4).** The cell pairs are those called by the niche-free calibrated statistic in at least one model: 30 in Xenium (marker-based labels) and 13 in CosMx (author labels). Retained features that detect one of the top-10 panel markers of either cell type were removed, and the statistic and $z^*$ were recomputed (Methods). Values are given as scGPT/Geneformer/C2S-scale. Before removal the models had 118/125/94 retained features in Xenium and 194/261/170 in CosMx. Models holding counts the models with $|z| \geq z^*$ and the sign of the all-feature call. A pair is spillover-robust if at least two of three models hold.

| Dataset | Cell pair | Call, all features | Models calling | Niche-free $z$, all features | Niche-free $z$, marker-excluded | Features removed | $z^*$, marker-excluded | Models holding | Spillover-robust |
|---|---|---|---|---|---|---|---|---|---|
| Xenium | lymphatic endothelial – B cell | enriched | 3 | +3.34/+3.82/+2.35 | +2.29/+2.95/+1.16 | 15/11/11 | 2.28/2.30/2.30 | 2/3 | yes |
| Xenium | monocyte – B cell | enriched | 3 | +7.24/+3.24/+2.85 | +4.45/+2.51/+0.48 | 19/15/13 | 2.28/2.30/2.30 | 2/3 | yes |
| Xenium | T cell – tumor-like epithelial | enriched | 3 | +3.47/+3.93/+5.49 | +1.12/+1.41/+1.57 | 20/18/19 | 2.23/2.23/2.23 | 0/3 | no |
| Xenium | tumor-like epithelial – dendritic | enriched | 1 | +0.23/+0.70/+2.83 | −1.19/−0.59/+1.76 | 23/22/23 | 2.23/2.23/2.23 | 0/3 | no |
| Xenium | tumor-like epithelial – fibroblast | enriched | 1 | +2.59/+0.69/+0.41 | +0.56/−0.97/−2.07 | 18/18/15 | 2.23/2.23/2.23 | 0/3 | no |
| Xenium | tumor-like epithelial – myeloid | enriched | 3 | +4.78/+6.57/+7.12 | −2.20/+1.89/+0.93 | 23/24/22 | 2.23/2.23/2.23 | 0/3 | no |
| Xenium | endothelial – plasma | depleted | 2 | −1.29/−2.42/−2.61 | −1.20/−2.18/−2.54 | 6/7/8 | 2.23/2.30/2.30 | 1/3 | no |
| Xenium | fibroblast – monocyte | depleted | 2 | −3.02/−3.73/+0.02 | −4.43/−4.34/−2.70 | 19/16/12 | 2.23/2.28/2.30 | 3/3 | yes |
| Xenium | monocyte – dendritic | depleted | 1 | −0.54/−2.93/−1.41 | −1.46/−2.85/−2.91 | 17/15/15 | 2.23/2.30/2.30 | 2/3 | yes |
| Xenium | monocyte – plasma | depleted | 2 | −3.13/−5.36/−2.12 | −4.49/−5.82/−4.50 | 15/13/12 | 2.23/2.30/2.30 | 3/3 | yes |
| Xenium | mural – monocyte | depleted | 1 | −1.09/−2.60/−0.69 | −2.07/−2.79/−1.03 | 23/19/17 | 2.23/2.26/2.28 | 1/3 | no |
| Xenium | mural – plasma | depleted | 3 | −2.40/−2.59/−2.60 | −3.43/−3.50/−3.53 | 16/14/14 | 2.23/2.28/2.30 | 3/3 | yes |
| Xenium | myeloid – dendritic | depleted | 1 | −1.56/−3.22/−1.57 | −1.44/−3.53/−3.18 | 19/18/17 | 2.23/2.26/2.28 | 2/3 | yes |
| Xenium | myeloid – endothelial | depleted | 3 | −2.23/−4.72/−2.56 | −2.94/−5.22/−4.31 | 15/16/14 | 2.23/2.26/2.28 | 3/3 | yes |
| Xenium | myeloid – fibroblast | depleted | 3 | −2.68/−8.08/−5.40 | −6.16/−9.99/−8.20 | 20/19/14 | 2.23/2.23/2.28 | 3/3 | yes |
| Xenium | myeloid – mast | depleted | 3 | −2.25/−3.91/−2.65 | −2.91/−4.19/−2.81 | 20/22/19 | 2.23/2.23/2.28 | 3/3 | yes |
| Xenium | myeloid – monocyte | depleted | 3 | −3.49/−6.79/−6.24 | −3.93/−7.04/−7.77 | 16/15/12 | 2.23/2.23/2.28 | 3/3 | yes |
| Xenium | myeloid – mural | depleted | 2 | −2.24/−3.42/−1.83 | −3.83/−4.45/−3.99 | 24/22/19 | 2.23/2.23/2.23 | 3/3 | yes |
| Xenium | myeloid – plasma | depleted | 3 | −6.31/−7.87/−6.76 | −6.43/−9.53/−9.42 | 16/16/14 | 2.23/2.28/2.28 | 3/3 | yes |
| Xenium | NK – fibroblast | depleted | 1 | −1.12/−3.50/−0.69 | −1.04/−4.01/−1.37 | 14/12/10 | 2.28/2.30/2.30 | 1/3 | no |
| Xenium | NK – monocyte | depleted | 1 | −0.93/−3.60/−2.06 | −1.90/−3.46/−2.57 | 17/14/13 | 2.26/2.30/2.30 | 2/3 | yes |
| Xenium | NK – myeloid | depleted | 3 | −2.55/−4.10/−3.11 | −3.01/−4.37/−4.24 | 18/17/15 | 2.23/2.28/2.28 | 3/3 | yes |
| Xenium | T cell – B cell | depleted | 2 | −0.47/−3.70/−2.80 | −2.75/−3.80/−2.74 | 16/13/13 | 2.28/2.30/2.30 | 3/3 | yes |
| Xenium | T cell – dendritic | depleted | 2 | −2.22/−4.55/−5.67 | −2.93/−3.85/−5.82 | 21/16/18 | 2.28/2.30/2.30 | 3/3 | yes |
| Xenium | T cell – endothelial | depleted | 1 | −1.07/−2.65/−1.68 | −2.79/−2.37/−3.99 | 12/10/12 | 2.26/2.30/2.30 | 3/3 | yes |
| Xenium | T cell – fibroblast | depleted | 1 | +0.29/−2.80/+0.13 | −2.32/−3.68/−4.20 | 17/13/11 | 2.28/2.30/2.30 | 3/3 | yes |
| Xenium | T cell – monocyte | depleted | 2 | −1.66/−7.67/−3.19 | −5.98/−8.44/−8.55 | 20/15/15 | 2.28/2.30/2.30 | 3/3 | yes |
| Xenium | T cell – myeloid | depleted | 1 | −1.57/−5.83/−0.88 | −6.17/−9.14/−5.32 | 21/18/17 | 2.28/2.28/2.30 | 3/3 | yes |
| Xenium | T cell – NK | depleted | 2 | −0.59/−3.10/−2.38 | −2.08/−4.18/−2.74 | 11/8/10 | 2.28/2.30/2.30 | 2/3 | yes |
| Xenium | T cell – plasma | depleted | 3 | −3.39/−7.63/−5.79 | −7.65/−9.98/−8.68 | 13/10/12 | 2.28/2.30/2.30 | 3/3 | yes |
| CosMx | fibroblast – t cd8 naive | enriched | 1 | +2.32/+0.61/−1.02 | +1.87/+0.54/−0.93 | 38/23/16 | 2.00/2.00/2.00 | 0/3 | no |
| CosMx | t cd4 memory – nk | enriched | 1 | +1.41/+0.58/+2.59 | +1.22/+0.29/+2.37 | 40/27/22 | 2.00/2.00/2.00 | 1/3 | no |
| CosMx | t cd8 memory – t cd8 naive | enriched | 2 | +2.79/+2.25/+1.48 | +2.39/+1.38/+1.40 | 34/14/11 | 2.00/2.00/2.00 | 1/3 | no |
| CosMx | treg – endothelial | enriched | 1 | +0.77/+2.43/+0.74 | +1.21/+2.35/+0.93 | 39/10/15 | 2.00/2.00/2.00 | 1/3 | no |
| CosMx | tumor – t cd4 memory | enriched | 1 | +1.13/+1.27/+2.53 | +0.26/+0.53/+0.56 | 72/59/54 | 2.00/2.00/2.00 | 0/3 | no |
| CosMx | b plasma – dendritic | depleted | 1 | −2.93/−0.52/−0.89 | −3.10/−0.75/−0.70 | 29/27/24 | 2.00/2.00/2.00 | 1/3 | no |
| CosMx | macrophage – endothelial | depleted | 1 | −2.05/−1.47/−0.33 | −1.52/−1.49/−0.39 | 34/20/23 | 2.00/2.00/2.00 | 0/3 | no |
| CosMx | macrophage – treg | depleted | 2 | −2.67/−0.97/−2.39 | −2.36/−0.87/−2.35 | 27/18/20 | 2.00/2.00/2.00 | 2/3 | yes |
| CosMx | t cd4 memory – dendritic | depleted | 1 | −3.19/−1.97/−1.26 | −3.51/−2.61/−2.05 | 28/22/22 | 2.00/2.00/2.00 | 3/3 | yes |
| CosMx | t cd4 memory – t cd8 memory | depleted | 1 | −2.56/−1.90/+0.58 | −2.94/−2.83/+0.17 | 34/22/17 | 2.00/2.00/2.00 | 2/3 | yes |
| CosMx | t cd4 memory – t cd8 naive | depleted | 1 | −2.24/+1.05/−0.41 | −1.75/+0.93/−0.24 | 36/21/15 | 2.00/2.00/2.00 | 0/3 | no |
| CosMx | tumor – dendritic | depleted | 1 | −1.45/−0.36/−2.29 | −1.28/−0.64/−2.50 | 67/63/56 | 2.00/2.00/2.00 | 1/3 | no |
| CosMx | tumor – epithelial non tumor | depleted | 1 | −2.74/−1.80/−1.73 | −3.02/−1.34/−0.93 | 65/56/50 | 2.00/2.00/2.00 | 1/3 | no |

Partner-marker gradient of the niche-free $z$ ($z$ in the top minus the bottom partner-marker tercile, divided by $|z|$; median over pairs). In Xenium the marker-excluded score had a gradient of −6.92 (95% CI −9.28 to −5.35), versus 0.71 (−0.41 to 1.24) for the full score. The Xenium calls that remain are therefore not spillover-free [1, 2]. In CosMx the marker-excluded gradient was −1.07 (−2.06 to 0.08), and that of the full score 0.07. Before marker removal, the niche-free $z$ of T cell–tumor-like epithelial contacts in Xenium was −17.3, −14.5, and −34.7 (scGPT, Geneformer, C2S-scale) in the lowest and +22.8, +20.5, and +39.8 in the highest partner-marker tercile, and 117 of the 139 niche-level Xenium calls at five niches (84%) had opposite signs in the lowest and highest tercile.

Supplementary Table S12: **Model-free LR co-localization screen (exploratory).** The screen covers every CellChatDB pair with all genes on the panel (Methods). Ligand+ and Receptor+ are the numbers of positive cells at ≥1 transcript. Fold is (obs + 1)/(null mean + 1) for ligand-positive–receptor-positive cell pairs within 100 coordinate units (18 µm in CosMx, 100 µm in Xenium) against a within-type null, at ≥1 and at ≥3 transcripts. q is Benjamini–Hochberg adjusted per dataset from a normal approximation, so values far below 0.05 are not to be read literally. A hit has $q < 0.05$ and fold ≥1.2. Robust, hit at ≥1 and ≥3 transcripts; enriched (≥1 only), hit at ≥1 transcript and not testable at ≥3; enriched (≥3 only), hit at ≥3 transcripts only; depleted, $q < 0.05$ and fold ≤1/1.2 at ≥1 transcript; n.t., not testable at ≥3 (fewer than 100 positive cells). Shown are the pairs that are a hit at either threshold or depleted, and EDN1–EDNRB for reference; the full screen (397 CosMx and 25 Xenium pairs) is deposited with the processed data (source tables table_S12_lr_screen_cosmx.csv and table_S12_lr_screen_xenium.csv).

| Dataset | Interaction | Pathway | Ligand+ | Receptor+ | Fold ≥1 | q ≥1 | Fold ≥3 | q ≥3 | Call |
|---|---|---|---|---|---|---|---|---|---|
| CosMx | CEACAM6 CEACAM6 | CEACAM | 11332 | 11332 | 1.36 | 0.0e+00 | 3.18 | 0.0e+00 | robust |
| CosMx | CDH1 CDH1 | CDH | 13703 | 13703 | 1.12 | 1.2e-90 | 1.70 | 6.5e-68 | enriched (≥3 only) |
| CosMx | PECAM1 PECAM1 | PECAM1 | 6907 | 6907 | 1.30 | 1.5e-90 | 2.21 | 5.6e-29 | robust |
| CosMx | HLA-DRA CD4 | MHC-II | 43418 | 9475 | 1.09 | 8.3e-81 | 1.32 | 3.3e-11 | enriched (≥3 only) |
| CosMx | HLA-DPA1 CD4 | MHC-II | 44271 | 9475 | 1.09 | 3.8e-80 | 1.38 | 8.7e-14 | enriched (≥3 only) |
| CosMx | ESAM ESAM | ESAM | 5410 | 5410 | 1.33 | 4.7e-80 | 2.25 | 3.7e-20 | robust |
| CosMx | HLA-DPB1 CD4 | MHC-II | 35705 | 9475 | 1.11 | 5.4e-80 | 1.61 | 5.6e-26 | enriched (≥3 only) |
| CosMx | HLA-DQB1 CD4 | MHC-II | 34583 | 9475 | 1.10 | 3.9e-73 | 1.43 | 1.6e-08 | enriched (≥3 only) |
| CosMx | CXCL9 CXCR3 | CXCL | 16041 | 4940 | 1.20 | 5.6e-62 | n.t. | | enriched (≥1 only) |
| CosMx | HLA-DRB5 CD4 | MHC-II | 25426 | 9475 | 1.12 | 1.6e-61 | 1.70 | 1.8e-13 | enriched (≥3 only) |
| CosMx | MIF CD74 CD44 | MIF | 44164 | 21731 | 1.05 | 5.8e-58 | 1.35 | 4.1e-77 | enriched (≥3 only) |
| CosMx | CCL5 CCR5 | CCL | 15499 | 4986 | 1.21 | 2.3e-55 | n.t. | | enriched (≥1 only) |
| CosMx | COL4A2 ITGA1 ITGB1 | COLLAGEN | 10820 | 4504 | 1.22 | 1.1e-48 | n.t. | | enriched (≥1 only) |
| CosMx | COL4A1 ITGA1 ITGB1 | COLLAGEN | 6408 | 4504 | 1.32 | 1.5e-45 | n.t. | | enriched (≥1 only) |
| CosMx | HLA-DQA1 CD4 | MHC-II | 40335 | 9475 | 1.08 | 1.8e-43 | 1.30 | 1.4e-08 | enriched (≥3 only) |
| CosMx | LGALS9 CD45 | GALECTIN | 11660 | 13693 | 1.11 | 9.1e-43 | 1.34 | 3.7e-02 | enriched (≥3 only) |
| CosMx | JAG1 NOTCH3 | NOTCH | 7625 | 9177 | 1.16 | 1.2e-41 | 2.80 | 4.1e-12 | enriched (≥3 only) |
| CosMx | HLA-DRB1 CD4 | MHC-II | 52201 | 9475 | 1.06 | 1.0e-36 | 1.21 | 3.6e-08 | enriched (≥3 only) |
| CosMx | LGALS9 CD44 | GALECTIN | 11660 | 23510 | 1.07 | 2.8e-36 | 1.83 | 1.0e-17 | enriched (≥3 only) |
| CosMx | CCL5 CCR1 | CCL | 15499 | 8990 | 1.12 | 1.7e-35 | 1.39 | 2.1e-03 | enriched (≥3 only) |
| CosMx | CCL3 CCR1 | CCL | 6765 | 8990 | 1.15 | 5.6e-25 | 2.24 | 1.1e-08 | enriched (≥3 only) |
| CosMx | SPP1 CD44 | SPP1 | 36506 | 23510 | 1.04 | 2.3e-24 | 1.24 | 1.3e-49 | enriched (≥3 only) |
| CosMx | HLA-E CD8A | MHC-I | 44702 | 6130 | 1.06 | 5.7e-23 | 1.23 | 5.1e-07 | enriched (≥3 only) |
| CosMx | ICAM1 ITGAX ITGB2 | ICAM | 23030 | 5664 | 1.09 | 7.7e-23 | 1.51 | 1.5e-05 | enriched (≥3 only) |

Continued on next page

Supplementary Table S12 (continued)

| Dataset | Interaction | Pathway | Ligand+ | Receptor+ | Fold ≥1 | q ≥1 | Fold ≥3 | q ≥3 | Call |
|---|---|---|---|---|---|---|---|---|---|
| CosMx | IL16 CD4 | IL16 | 11393 | 9475 | 1.09 | 6.5e-19 | 1.71 | 1.8e-02 | enriched (≥3 only) |
| CosMx | CCL8 CCR1 | CCL | 5137 | 8990 | 1.14 | 8.5e-19 | 2.22 | 4.2e-04 | enriched (≥3 only) |
| CosMx | APP TNFRSF21 | APP | 23713 | 9522 | 1.06 | 4.7e-16 | 1.33 | 1.5e-07 | enriched (≥3 only) |
| CosMx | CD80 CD274 | CD80 | 6171 | 9954 | 1.10 | 1.4e-14 | 2.62 | 2.9e-05 | enriched (≥3 only) |
| CosMx | PDGFA PDGFRB | PDGF | 8552 | 6854 | 1.11 | 1.8e-12 | 2.53 | 9.7e-06 | enriched (≥3 only) |
| CosMx | EFNB1 EPHB4 | EPHB | 8310 | 11598 | 1.07 | 8.0e-12 | 1.45 | 3.1e-02 | enriched (≥3 only) |
| CosMx | CSF1 CSF1R | CSF | 13755 | 7655 | 1.07 | 2.8e-10 | 1.68 | 2.3e-02 | enriched (≥3 only) |
| CosMx | CCL3L1 CCR1 | CCL | 6437 | 8990 | 1.09 | 1.6e-09 | 2.13 | 1.5e-05 | enriched (≥3 only) |
| CosMx | COL9A2 CD44 | COLLAGEN | 42641 | 23510 | 1.02 | 4.5e-09 | 1.22 | 2.4e-27 | enriched (≥3 only) |
| CosMx | HLA-C CD8A | MHC-I | 68189 | 6130 | 1.03 | 1.9e-07 | 1.23 | 1.7e-20 | enriched (≥3 only) |
| CosMx | CDH5 CDH5 | CDH5 | 5054 | 5054 | 1.10 | 2.3e-06 | 2.91 | 2.8e-09 | enriched (≥3 only) |
| CosMx | CD55 ADGRE5 | ADGRE | 24398 | 9013 | 1.03 | 4.0e-06 | 1.47 | 2.0e-06 | enriched (≥3 only) |
| CosMx | CD80 CD28 | CD80 | 6171 | 6538 | 1.07 | 3.0e-04 | 2.27 | 2.3e-02 | enriched (≥3 only) |
| Xenium | CCL19 CCR7 | CCL | 18873 | 33756 | 2.58 | 0.0e+00 | 4.51 | 0.0e+00 | robust |
| Xenium | CDH1 CDH1 | CDH | 12828 | 12828 | 1.67 | 0.0e+00 | 2.55 | 0.0e+00 | robust |
| Xenium | CLEC2C KLRB1 | CLEC | 52097 | 32951 | 1.47 | 0.0e+00 | 1.62 | 0.0e+00 | robust |
| Xenium | HLA-DRB5 CD4 | MHC-II | 118663 | 127528 | 1.07 | 0.0e+00 | 1.32 | 0.0e+00 | enriched (≥3 only) |
| Xenium | PTN SDC1 | PTN | 24550 | 46014 | 1.13 | 4.0e-250 | 1.33 | 4.7e-96 | enriched (≥3 only) |
| Xenium | NCAM1 NCAM1 | NCAM | 5015 | 5015 | 1.20 | 1.0e-83 | 1.53 | 2.7e-05 | enriched (≥3 only) |
| Xenium | CD274 PDCD1 | PD-L1 | 17748 | 14585 | 1.07 | 5.5e-36 | 1.34 | 3.1e-08 | enriched (≥3 only) |
| Xenium | CD70 CD27 | CD70 | 11375 | 48157 | 1.05 | 1.7e-13 | 1.34 | 1.7e-14 | enriched (≥3 only) |
| Xenium | EDN1 EDNRB | EDN | 44512 | 58531 | 1.01 | 7.0e-06 | 1.16 | 1.6e-94 | |
| Xenium | SELL CD34 | SELL | 49201 | 70480 | 0.81 | 0.0e+00 | 0.60 | 0.0e+00 | depleted |
| Xenium | CXCL10 ACKR1 | CXCL | 67722 | 16326 | 0.77 | 0.0e+00 | 0.40 | 0.0e+00 | depleted |
| Xenium | CXCL6 ACKR1 | CXCL | 128386 | 16326 | 0.62 | 0.0e+00 | 0.26 | 0.0e+00 | depleted |

Supplementary Table S13: **Replication of the CosMx LR co-localization hits in four further donors.** The screen of Supplementary Table S12 was run unchanged in each donor (Methods). Targets are the three robust hits and the four heterotypic hits at ≥1 transcript of the analyzed section (Lung13). **a,** Fold at ≥1 transcript within 100 units (18 µm), with the Benjamini–Hochberg q, adjusted per section, in brackets. *, hit ($q < 0.05$ and fold ≥1.2). Same direction counts donors with fold > 1 and $q < 0.05$. A target replicates if it is a hit at ≥1 transcript in at least three of four donors. **b,** Niche context (descriptive). For each model and donor niche, the target's share of close pairs was divided by the niche's share of cells. Each model's peak niche is shown, with its leading cell type in brackets.

**a** Fold at ≥1 transcript (q) and replication; tx, transcripts

| Target | Lung13 (analyzed) | Lung5 | Lung6 | Lung9 | Lung12 | Donor hits, ≥1 tx | Same direction | Donor hits, ≥3 tx | Replicates |
|---|---|---|---|---|---|---|---|---|---|
| CEACAM6–CEACAM6 | 1.36* (0.0e+00) | 1.10 (0.0e+00) | 1.70* (1.1e-87) | 1.16 (3.5e-08) | 1.81* (0.0e+00) | 2/4 | 4/4 | 2/4 | no |
| ESAM–ESAM | 1.33* (4.7e-80) | 1.25* (3.1e-107) | 1.30* (1.5e-25) | 1.12 (1.9e-11) | 1.05 (1.1e-03) | 2/4 | 4/4 | 3/4 | no |
| PECAM1–PECAM1 | 1.30* (1.5e-90) | 1.22* (4.3e-168) | 1.28* (2.4e-37) | 1.18 (2.8e-64) | 1.10 (3.1e-12) | 2/4 | 4/4 | 4/4 | no |
| CXCL9–CXCR3 | 1.20* (5.6e-62) | 1.08 (9.9e-04) | 1.04 (3.0e-01) | 1.07 (2.6e-03) | 1.04 (1.9e-02) | 0/4 | 3/4 | 0/4 | no |
| CCL5–CCR5 | 1.21* (2.3e-55) | 1.08 (3.9e-08) | 1.16 (2.1e-04) | 1.06 (5.8e-04) | 1.08 (3.4e-06) | 0/4 | 4/4 | 0/4 | no |
| COL4A2–ITGA1/ITGB1 | 1.22* (1.1e-48) | 1.20* (4.1e-190) | 1.21* (2.0e-104) | 1.15 (2.8e-113) | 1.16 (1.1e-38) | 2/4 | 4/4 | 2/4 | no |
| COL4A1–ITGA1/ITGB1 | 1.32* (1.5e-45) | 1.24* (3.3e-187) | 1.23* (1.4e-96) | 1.17 (2.9e-64) | 1.25* (3.7e-56) | 3/4 | 4/4 | 3/4 | yes |

The same code on the analyzed section reproduces Supplementary Table S12. Close pairs of COL4A1–ITGA1/ITGB1 were mostly between tumor cells in Lung6 and Lung9 (80% and 74%), so its co-localization is not a demonstrated cross-type interface.

**Supplementary Table S13 (continued).** Panel b is described in the caption on the previous page.

**b** Peak niche per model: close-pair share over cell share (leading cell type)

| Donor | Target | scGPT | Geneformer | C2S-scale |
|---|---|---|---|---|
| Lung5 | CEACAM6–CEACAM6 | 4.73 (tumor 5) | 4.93 (tumor 5) | 4.96 (tumor 5) |
| Lung5 | ESAM–ESAM | 1.67 (fibroblast) | 1.68 (fibroblast) | 1.69 (fibroblast) |
| Lung5 | PECAM1–PECAM1 | 1.87 (fibroblast) | 1.79 (fibroblast) | 1.86 (fibroblast) |
| Lung5 | CXCL9–CXCR3 | 1.25 (tumor 5) | 1.19 (tumor 5) | 1.19 (tumor 5) |
| Lung5 | CCL5–CCR5 | 1.20 (neutrophil) | 1.29 (neutrophil) | 1.32 (neutrophil) |
| Lung5 | COL4A2–ITGA1/ITGB1 | 2.04 (fibroblast) | 1.88 (fibroblast) | 2.01 (fibroblast) |
| Lung5 | COL4A1–ITGA1/ITGB1 | 2.09 (fibroblast) | 1.95 (fibroblast) | 2.09 (fibroblast) |
| Lung6 | CEACAM6–CEACAM6 | 1.26 (fibroblast) | 1.44 (macrophage) | 1.21 (epithelial) |
| Lung6 | ESAM–ESAM | 3.45 (fibroblast) | 2.89 (macrophage) | 3.09 (epithelial) |
| Lung6 | PECAM1–PECAM1 | 4.01 (fibroblast) | 3.83 (macrophage) | 3.78 (epithelial) |
| Lung6 | CXCL9–CXCR3 | 1.04 (tumor 6) | 1.44 (tumor 6) | 1.04 (tumor 6) |
| Lung6 | CCL5–CCR5 | 1.90 (macrophage) | 2.37 (macrophage) | 2.05 (epithelial) |
| Lung6 | COL4A2–ITGA1/ITGB1 | 1.16 (fibroblast) | 1.69 (tumor 6) | 1.15 (tumor 6) |
| Lung6 | COL4A1–ITGA1/ITGB1 | 1.15 (tumor 6) | 1.71 (tumor 6) | 1.16 (tumor 6) |
| Lung9 | CEACAM6–CEACAM6 | 1.38 (neutrophil) | 1.26 (neutrophil) | 1.26 (tumor 9) |
| Lung9 | ESAM–ESAM | 1.29 (tumor 9) | 1.13 (neutrophil) | 1.14 (tumor 9) |
| Lung9 | PECAM1–PECAM1 | 1.80 (tumor 9) | 1.44 (tumor 9) | 1.52 (tumor 9) |
| Lung9 | CXCL9–CXCR3 | 1.74 (tumor 9) | 1.47 (tumor 9) | 1.52 (tumor 9) |
| Lung9 | CCL5–CCR5 | 1.23 (neutrophil) | 1.26 (neutrophil) | 1.19 (neutrophil) |
| Lung9 | COL4A2–ITGA1/ITGB1 | 2.23 (tumor 9) | 1.79 (tumor 9) | 1.88 (tumor 9) |
| Lung9 | COL4A1–ITGA1/ITGB1 | 2.13 (tumor 9) | 1.70 (tumor 9) | 1.79 (tumor 9) |
| Lung12 | CEACAM6–CEACAM6 | 3.89 (tumor 12) | 4.56 (tumor 12) | 4.80 (tumor 12) |
| Lung12 | ESAM–ESAM | 1.91 (tumor 12) | 1.63 (tumor 12) | 1.72 (tumor 12) |
| Lung12 | PECAM1–PECAM1 | 1.27 (tumor 12) | 1.38 (neutrophil) | 1.07 (neutrophil) |
| Lung12 | CXCL9–CXCR3 | 1.59 (tumor 12) | 1.30 (tumor 12) | 1.37 (tumor 12) |
| Lung12 | CCL5–CCR5 | 1.32 (tumor 12) | 1.33 (neutrophil) | 1.13 (neutrophil) |
| Lung12 | COL4A2–ITGA1/ITGB1 | 1.58 (fibroblast) | 1.15 (fibroblast) | 1.70 (fibroblast) |
| Lung12 | COL4A1–ITGA1/ITGB1 | 1.57 (fibroblast) | 1.13 (fibroblast) | 1.73 (fibroblast) |

The three models' peak niches have the same leading cell type in 19 of 28 donor–target cases (post hoc tally).

Supplementary Table S14: **Edge support of the residual maps in Figures 7 and 8.** Edges are cell pairs of the two groups within 138 units in CosMx and 120 units in Xenium. Distances are in coordinate units. In CosMx these are pixels of 0.18 µm, so 138 units are 24.8 µm, the median edge is 17.3 µm, and the 90th percentile is 23.5 µm. In Xenium they are µm. Edge geometry is independent of the features. Residuals use each model's count-adjusted program score. The residual display limit is the 98th percentile of |residual| pooled for the panel in CosMx and the 99th percentile per model in Xenium.

| Dataset | Model | Cell-group pair | Edges | Edges with residual | Median distance | 90th percentile distance | Residual display limit |
|---|---|---|---|---|---|---|---|
| CosMx | scGPT | Endothelial/fibroblast | 6060 | 5477 | 96.143 | 130.311 | 2.36 |
| CosMx | Geneformer | Endothelial/fibroblast | 6060 | 5477 | 96.143 | 130.311 | 2.36 |
| CosMx | C2S-scale | Endothelial/fibroblast | 6060 | 5477 | 96.143 | 130.311 | 2.36 |
| Xenium | scGPT | Endothelial/tumor-like epithelial | 1108170 | 1107452 | 87.130 | 114.208 | 2.69 |
| Xenium | Geneformer | Endothelial/tumor-like epithelial | 1108170 | 1107452 | 87.130 | 114.208 | 2.77 |
| Xenium | C2S-scale | Endothelial/tumor-like epithelial | 1108170 | 1107452 | 87.130 | 114.208 | 2.78 |

Supplementary Table S15: **CosMx case-study evidence.** Pairs ≤100 is the number of ligand-positive source and receptor-positive target cell pairs within 100 units (18 µm; CosMx pixels of 0.18 µm) in the crop. Folds are given against each null (1,000 permutations per null), with the upper- and lower-tail P. z is the niche-free calibrated statistic for the cell pair. Supplementary Figure S12 displays 5 links for C2, 7 for C1, and 3 for C3. JCHAIN/PECAM1 is a marker pair and not an LR system. ITGA1 is one component of the nominated receptor complex.

| Case | Labels | Cell pair | Genes | Pairs ≤100 | Fold, position null (P upper; P lower) | Fold, within-type null (P upper; P lower) | scGPT z | Geneformer z | C2S-scale z | $z^*$ | Note |
|---|---|---|---|---|---|---|---|---|---|---|---|
| C2 | author | mast/tumor | AREG/EGFR | 3 | 0.80 (0.822; 0.394) | 1.14 (0.470; 0.762) | 0.53 | -0.45 | -0.77 | 2.00 | shown |
| C1 | author | endothelial/fibroblast | COL9A2/ITGA1 | 4 | 1.22 (0.399; 0.743) | 0.91 (0.613; 0.534) | 1.42 | -0.94 | -0.21 | 2.00 | shown |
| C3 | author | b plasma/endothelial | JCHAIN/PECAM1 | 1 | 0.50 (0.916; 0.250) | 0.49 (0.910; 0.245) | -0.38 | -0.34 | 0.96 | 2.00 | shown |

Supplementary Table S16: **Xenium case-study evidence on marker-based labels (primary) and transferred labels (reference).** Columns as in Supplementary Table S15, with pairs counted within 100 µm (Xenium coordinates are µm). $z^*$ is given as the range over models and label sets. PT-like denotes cells assigned to the reference proximal-tubule class (Supplementary Table S7). The X2 crop was selected for display before the proximity tests, which are also shown in Supplementary Figure S13c.

| Case | Labels | Cell pair | Genes | Pairs ≤100 | Fold, position null (P upper; P lower) | Fold, within-type null (P upper; P lower) | scGPT z | Geneformer z | C2S-scale z | $z^*$ | Note |
|---|---|---|---|---|---|---|---|---|---|---|---|
| X2 | marker | endothelial/tumor-like epithelial | EDN1/EDNRB | 492 | 1.19 (0.001; 1.000) | 1.18 (0.011; 0.992) | 0.77 | 0.31 | 0.99 | 2.23–2.34 | Figure 8c |
| X2 | transferred | endothelial/PT-like | EDN1/EDNRB | 435 | 1.19 (0.010; 0.991) | 1.24 (0.003; 0.998) | 0.11 | -1.97 | 0.61 | 2.23–2.34 | reference |

Supplementary Table S17: **Prediction of neighborhood composition from a macrophage's own expression in CosMx (exploratory).** Targets are the T-cell and myeloid shares of the macrophage's 75-µm neighborhood, in the analyzed section only. Ridge regression was evaluated by blocked five-fold cross-validation with 500-µm blocks (Methods). CD3-IF is the CD3 immunofluorescence inside the macrophage. $R^2$ uses a fixed ridge penalty ($\alpha = 10$). With the penalty chosen by inner cross-validation over $\alpha = 1$ to $10^6$, as in Figure 7b and Supplementary Table S4d, the genes give $R^2$ 0.43 (T share) and 0.46 (myeloid share).

| Macrophages | Mean T share | Mean myeloid share | $R^2$ genes → T share | $R^2$ genes → myeloid | $R^2$ intrinsic genes → T | $\rho$ CD3-IF vs T share | $R^2$ CD3-IF alone | $R^2$ intrinsic after CD3-IF |
|---|---|---|---|---|---|---|---|---|
| 17989 | 0.159 | 0.352 | 0.419 | 0.445 | 0.416 | 0.40 | 0.103 | 0.288 |

Supplementary Table S18: **Sparse C2S-scale cell profiles and genes as predictors of macrophage neighborhood composition in CosMx (exploratory; 17,989 macrophages).** $p$ is the number of predictors. The fixed penalty ($\alpha = 10$) was the primary setting and the $\alpha$ grid the sensitivity setting. The fixed penalty gives negative $R^2$ when predictors outnumber macrophages, so the grid values are reported. The grid setting chooses $\alpha$ per target from a grid scaled by training size (relative $\alpha$ in brackets). For the latent rows the chosen $\alpha$ is at the edge of the grid, so those values are lower bounds. Figure 7b and Supplementary Table S4d choose $\alpha$ by inner cross-validation over $\alpha = 1$ to $10^6$, which gives 0.43 and 0.46 for the genes and 0.42 and 0.43 for the sparse C2S-scale profile (T and myeloid share), within 0.004 of the grid values here.

| Predictors | $p$ | $R^2$ T share ($\alpha = 10$) | $R^2$ myeloid ($\alpha = 10$) | $R^2$ T share ($\alpha$ grid) | $R^2$ myeloid ($\alpha$ grid) |
|---|---|---|---|---|---|
| genes (960 panel genes) | 960 | 0.419 | 0.445 | 0.431 (1.0) | 0.452 (0.1) |
| sparse SAE, C2S-scale Layer 14 (layer_13 in the code), latents active in ≥1% of macrophages (18,432) | 18432 | -3.946 | -3.730 | 0.421 (10.0) | 0.426 (10.0) |
| sparse SAE, the 170 retained spatial features | 170 | 0.273 | 0.265 | 0.274 (0.1) | 0.266 (0.01) |
| genes + sparse SAE (≥1%) | 19392 | -3.018 | -2.906 | 0.426 (10.0) | 0.433 (10.0) |
| intrinsic genes only | 938 | 0.416 | 0.445 | 0.426 (1.0) | 0.451 (0.1) |

Supplementary Table S19: **Exploratory analyses.** S1–S5 and C1–C3 use single sections, without interface z-scores or cross-donor data. C1 was repeated with the same criterion at the five Xenium niches of the main text. C4 uses the calibrated interface statistic (Supplementary Table S11). I1 uses the four further CosMx donors, and I2 uses three serial sections not used elsewhere as its confirmation set (Supplementary Note S3). D1 is descriptive and had no criterion.

| Analysis | Question | Criterion | Result |
|---|---|---|---|
| S1 | Are concept-labeled features spatially smoother than the mean expression of their label's panel genes? | Median Moran's I difference $> 0$, 95% CI excluding 0, in $\geq 2$ of 3 models | Features less autocorrelated: median $\Delta I$ $-0.08$ to $-0.09$ (CosMx), $-0.04$ to $-0.06$ (Xenium); CIs exclude 0 |
| S2 | Do features inside CosMx macrophages track distance to tumor better than label genes? | p90 $\lvert\rho\rvert$ features > gene sets in $\geq 2$ of 3 models, CI excluding 0 in $\geq 1$ | Below gene sets in 3/3; top features carry keratin labels and track the macrophage's own PanCK ($\rho$ 0.27–0.31): spillover [1, 2] |
| S3 | Do features inside Xenium tumor-like epithelium track distance to vessels better than label genes? | As S2, judged on tumor-like epithelium | Below gene sets in 3/3; weak gradients ($\lvert\rho\rvert \leq 0.24$) carried by single genes |
| S4 | Do niche-defining feature labels match gene-level niche enrichment (ORA of top-50 DE genes)? | Overlap above a random-label null ($P < 0.05$) in $\geq 2$ of 3 niches for $\geq 2$ of 3 CosMx models | CosMx: 4–28% of labels overlap, $P \leq 0.035$ in 6 of 9 model-niche pairs (macrophage-rich and stromal niches); Xenium at chance in 8 of 9 |
| S5 | Is a within-type two-state split on features more spatially coherent than on label genes? | Moran's I difference $> 0.05$, CI excluding 0, in $\geq 2$ of 3 models | Not in any type; splits follow transcript depth ($\lvert\rho\rvert$ 0.43–0.82) |
| C1 | Do matched-niche CellChat pathway profiles agree across models, with recurrent niche-specific pathways? | (i) matched > unmatched Spearman for all model pairs; (ii) $\geq 1$ recurrent niche-specific pathway in $\geq 2$ of 3 niches | CosMx: 0.88–0.94 vs 0.79–0.80, specific pathways in the tumor niche only; Xenium, three niches: 0.94–1.00 vs 0.71–0.73, specific pathways in 2 niches; Xenium, five niches: 0.87–0.91 vs 0.69–0.75, no pathway specific to the same niche in all three models (CCL, B/T-lymphoid, and PTN, tumor/myeloid, in two) |
| C2 | Is concept-label support of CellChat rows niche-specific? | Own-minus-swapped support $\geq 0.10$ in $\geq 2$ of 3 niches for $\geq 2$ of 3 CosMx models | Reached in 2 models. Support depended on which ligand and receptor genes a niche's term list contained rather than on the pair (17–99% of rows per niche in CosMx; 8–99.5% in Xenium at five niches, 32–100% at three) and transferred between niches; reported as a descriptor |
| C3 | Which panel LR pairs colocalize beyond a within-type null? | $\geq 1$ robust enriched hit per dataset ($q < 0.05$, fold $\geq 1.2$ at $\geq 1$ and $\geq 3$ transcripts) | CosMx: CEACAM6, ESAM, PECAM1 (homophilic); Xenium: CCL19–CCR7, CLEC2C–KLRB1, CDH1–CDH1; EDN1–EDNRB fold 1.01 |
| C4 | Do interface calls survive removal of the features that detect either partner's markers? | Spillover-robust pair: $\lvert z\rvert \geq z^*$ with the sign of the all-feature call in $\geq 2$ of 3 models | Xenium: 23 of 30 pairs, 21 of them depletions; all four tumor-like epithelial enrichments (two called by all three models) held in none of the three models; the reduced score remains partner-patterned. CosMx: 3 of 13 pairs, all depletions (Supplementary Table S11) |

Continued on next page

Supplementary Table S19 (continued)

| Analysis | Question | Criterion | Result |
|---|---|---|---|
| I1 | Does a tumor cell's own expression, without T, NK, and B markers, predict the T-cell share of its 75-µm neighborhood? | $R^2 \geq 0.15$ in the analyzed section and $\geq$0.10 in $\geq$3 of 4 donors | $R^2$ 0.138 in the analyzed section; 0.14–0.24 in the donors (4 of 4 $\geq$0.10); in the donors the signal largely reflects tumor-edge position (post hoc; Supplementary Note S3) |
| I2 | Do interferon-$\gamma$-response and hypoxia scores of tumor cells track local T-cell infiltration? | Partial $\rho \geq +0.10$ (interferon-$\gamma$) and $\leq -0.10$ (hypoxia), CIs excluding 0, in $\geq$2 of 3 serial sections | Interferon-$\gamma$ $-0.008$ to $+0.040$; hypoxia $-0.019$ to $-0.009$ in the three confirmation sections (Supplementary Note S3) |
| D1 | Are ligand- and receptor-positive cells closer than expected in the CosMx case-study crops (Supplementary Figure S12)? | None (descriptive; crops chosen for display before the tests) | 1–4 pairs within 100 units (18 µm) per crop; fold 0.49–1.22, all upper-tail $P > 0.39$ under a target-position and a within-type null; no crop's cell pair model-supported (niche-free $z$ $-0.94$ to $+1.42$; $z^* = 2.0$; Supplementary Figure S13b, Table S15) |

Supplementary Table S20: **Co-localization screen at matched neighbor counts.** The screen of Supplementary Table S12 was repeated at the radius where the mean number of other cells within reach equals that of the other dataset at 100 coordinate units (Methods). In Xenium this radius is 15.4 µm, which holds 7.2 cells on average, as do 100 pixels (18 µm) in CosMx. In CosMx it is 652.8 pixels (117.5 µm), which holds 287 cells, as do 100 µm in Xenium. Radii were interpolated on the neighbor-count curve and rounded to 0.1 unit. Fold is at ≥1 transcript unless stated. *, hit; robust and n.t. as in Supplementary Table S12; tx, transcripts. Folds just below the 1.2 cut are shown with four decimals. **a,** Robust hits of the 100-unit screen, the new hits, and EDN1–EDNRB for reference. **b,** The seven CosMx targets of Supplementary Table S13 in the analyzed section (Lung13) and the four donors at the matched radius. Donor hits are counted at ≥1 transcript, at 100 units → the matched radius. Same direction counts donors with fold > 1 and $q < 0.05$. **c,** Xenium five-niche placement as in Figure 5c: the ratio of a hit's share of close pairs in its peak niche to that niche's share of cells, at 100 µm → 15.4 µm.

**a** Section-level screen

| Dataset | Pair | Fold, 100 units | Fold, matched | Fold ≥3 tx, matched | Robust, matched |
|---|---|---|---|---|---|
| CosMx | CEACAM6–CEACAM6 | 1.36 (robust) | 1.27* | 2.44* | yes |
| CosMx | ESAM–ESAM | 1.33 (robust) | 1.13 | 1.56* | no (≥3 only) |
| CosMx | PECAM1–PECAM1 | 1.30 (robust) | 1.16 | 1.75* | no (≥3 only) |
| CosMx | INHBA/INHBB (activin AB)–ACVR1B/ACVR2A | 1.04 | 1.22* | n.t. | no (≥1 only) |
| Xenium | CCL19–CCR7 | 2.58 (robust) | 2.62* | 4.34* | yes |
| Xenium | CLEC2C–KLRB1 | 1.47 (robust) | 1.61* | 1.79* | yes |
| Xenium | CDH1–CDH1 | 1.67 (robust) | 1.98* | 2.40* | yes |
| Xenium | NCAM1–NCAM1 | 1.1994 | 2.16* | 2.72* | yes |
| Xenium | EDN1–EDNRB | 1.01 | 0.85 | 0.66 | no |

At 15.4 µm, EDN1–EDNRB is in the depleted direction but above the depletion cut (fold ≤1/1.2; Supplementary Table S12). In CosMx the number of hits at ≥3 transcripts rose from 33 to 52, none of them robust. Hit sets are specific to the radius, and the main analysis uses 100 coordinate units.

**b** CosMx donors at 117.5 µm

| Target | Lung13 (analyzed) | Lung5 | Lung6 | Lung9 | Lung12 | Donor hits, 100 → matched | Same direction |
|---|---|---|---|---|---|---|---|
| CEACAM6–CEACAM6 | 1.27* | 1.09 | 1.1998 | 1.09 | 1.50* | 2/4 → 1/4 | 4/4 |
| ESAM–ESAM | 1.13 | 1.10 | 1.19 | 1.09 | 1.08 | 2/4 → 0/4 | 4/4 |
| PECAM1–PECAM1 | 1.16 | 1.09 | 1.17 | 1.13 | 1.07 | 2/4 → 0/4 | 4/4 |
| CXCL9–CXCR3 | 1.14 | 1.11 | 1.07 | 1.08 | 1.09 | 0/4 → 0/4 | 4/4 |
| CCL5–CCR5 | 1.14 | 1.08 | 1.14 | 1.08 | 1.10 | 0/4 → 0/4 | 4/4 |
| COL4A2–ITGA1/ITGB1 | 1.10 | 1.12 | 1.22* | 1.12 | 1.14 | 2/4 → 1/4 | 4/4 |
| COL4A1–ITGA1/ITGB1 | 1.16 | 1.13 | 1.24* | 1.15 | 1.19 | 3/4 → 1/4 | 4/4 |

**c** Xenium niche placement at 15.4 µm

| Pair | Peak niche | scGPT | Geneformer | C2S-scale |
|---|---|---|---|---|
| CCL19–CCR7 | B/T-lymphoid | 3.20 → 3.26 | 4.34 → 4.24 | 3.10 → 3.08 |
| CLEC2C–KLRB1 | B/T-lymphoid | 2.83 → 2.92 | 3.73 → 3.68 | 2.71 → 2.80 |
| CDH1–CDH1 | Tumor/myeloid | 2.94 → 2.34 | 2.42 → 1.93 | 2.94 → 2.25 |
| NCAM1–NCAM1 | B/T-lymphoid | 1.64 | 1.80 | 1.74 |
